# Single-cell transcriptome analysis reveals heterogeneity and a dynamic regenerative response of quiescent radial glia in adult zebrafish brain

**DOI:** 10.1101/2022.07.27.501663

**Authors:** Manana Kutsia, Yuki Takeuchi, Nishtha Ranawat, Ichiro Masai

**Affiliations:** Developmental Neurobiology Unit, Okinawa Institute of Science and Technology Graduate University, Tancha 1919-1, Onna, Okinawa 904-0495, Japan

**Keywords:** radial glia, adult neural stem cell, neural regeneration, microglia, zebrafish

## Abstract

In zebrafish telencephalon, radial glial cells (RGs) show a remarkable ability to regenerate damaged neural tissue by re-initiating cell proliferation to produce neural precursors to rebuild the lost neural circuit. However, it is not fully understood how RGs respond to brain damage to initiate this regenerative response. Here we applied single-cell transcriptomics to RGs in adult zebrafish telencephalon and identified five RG subtypes, which are classified into four quiescent RGs (qRGs) and one proliferating RG (pRG). The four qRGs differentially express distinct subsets of qRG markers, suggesting heterogeneity of qRG in zebrafish adult brain. Interestingly, one qRG subtype shows high expression of ribosomal proteins, and its fraction increases in response to brain damage. Consistently, the mTOR pathway is activated in RGs near the injury site. It was reported that inflammatory responses of brain-resident immune cells, microglia, are required for inducing regenerative responses of RGs in zebrafish. Genetical elimination of microglia not only suppressed the damage-induced regenerative response of RGs, but also decreased the fraction of the ribosomal expression-enriched qRGs. Our pseudo-time analysis suggests that putative dormant RGs produce ribosomal expression-enriched qRGs through activation of ribosomal genesis, as well as suppression of cholesterol biogenesis, and pRGs through activation of the JAK/STAT pathway. Our findings reveal heterogeneity of qRGs in adult zebrafish brain and their dynamic regenerative response to brain damage.

## Introduction

In mammals, most neurons in the brain are generated during embryonic development; however, production of new neurons is sustained throughout adulthood in only two regions of the telencephalon, the ventricular/subventricular zone (V/SVZ) of the lateral ventricle and the subgranular zone (SGZ) of the dentate gyrus of the hippocampus (Kriegstein and Alvarez-Buylla, 2009; Zhao et al., 2008). In the SGZ, radial glial stem cells (Type 1 cells) generate amplifying progenitor cells (Type 2 cells) and then neuroblasts (Type 3 cells), which eventually generate granule interneurons. Dentate gyrus granule neurons extend axons called mossy fibers to target pyramidal neurons in the CA3 and CA2 hippocampal regions (Cope and Gould, 2019). In the V/SVZ, neural stem cells (B1 cells) generate transit-amplifying cells (C cells). C cells subsequently generate neuroblasts (A cells), which migrate toward the olfactory bulb, where they differentiate into local interneurons (Otsuki and Brand, 2020).

In contrast to the limited neurogenic capability of adult mammals, teleosts such as zebrafish show remarkably higher proliferative and regenerative potential in the nervous system (Grandel and Brand, 2013; Lindsey and Tropepe, 2006). In zebrafish, composition of neurogenic niches and the cellular nature of neurogenic progenitors have been intensively studied. Classic analysis of zebrafish adult brain with proliferative markers identified 16 different proliferative zones, which produce newborn neurons (Adolf et al., 2006; Grandel et al., 2006). In zebrafish, two distinct neurogenic proliferation zones are formed at the ventricular surface of the dorsal and ventral telencephalon, respectively (Barbosa and Ninkovic, 2016; Jurisch-Yaksi et al., 2020). In the dorsal telencephalon, called the pallium, radial glial cells (RGs) are located at the ventricular surface and extend long radial processes, whose endfeet reach the pial surface close to a blood vessel at the lateral margin of the dorsal telencephalon (Ganz et al., 2010; Marz et al., 2010). These pallial ventricular RGs highly express glial markers such as GFAP, Vimentin and S100β, and generate newborn neurons. In the lateral domain of the pallium, the same process operates, but RGs are generated during juvenile and adult stages from neuroepithelial progenitor cells located at the interface between the pallium and subpallium (Dirian et al., 2014), suggesting that the source of RGs is different from that of RGs in the medial and dorsal pallium. On the other hand, in the ventral telencephalon, called the subpallium, neuroepithelial-like progenitor cells are located in the ventricular region (Ganz et al., 2010; Marz et al., 2010). They rapidly proliferate to generate neuroblasts migrating toward the olfactory bulb, so this neurogenic niche may share characteristics with glial marker-negative proliferating neuroblasts in mouse V/SVZ. Thus, there is heterogeneity in progenitor cell types in the adult zebrafish telencephalon.

In general, severe trauma brain injury (TBI) causes direct damage to neural tissue in the form of primary lesions (Xiong et al., 2013). Following these primary injuries, lasting damage is sustained through a series of complex events called secondary injuries (Jassam et al., 2017). In mammals, brain damage induces acute inflammation, microglial activation, and reactive astrogliosis, leading to scar formation, which limits spread of damage into unaffected CNS area. However, significant regeneration does not occur. In contrast, stab injuries to the zebrafish telencephalon cause an acute inflammatory response and reactive gliosis; however, neither persistent gliosis, fibrotic scar formation nor chronic inflammation occurs (Baumgart et al., 2012; Kishimoto et al., 2012; Kroehne et al., 2011). Rather, RGs upregulate cell proliferation at 2-3 days post-lesion (dpl), and subsequently promote neurogenesis, which leads to rebuild neural circuits of damaged tissue.

Quantitative clonal analysis of RGs in zebrafish dorsal pallium revealed that RGs comprise two groups of neural stem cells (NSCs): deeply quiescent and self-renewing neural stem cells called reservoir NSCs and neural stem cells called operational NSCs, which promote neurogenesis. Operational NSCs undergo various cell division modes: symmetric gliogenic division, asymmetric neurogenic division, symmetric neurogenic division, and direct conversion to neurons without cell division, in normal homeostasis (Rothenaigner et al., 2011; Than-Trong et al., 2020). Furthermore, *in vivo* live imaging of RGs in zebrafish dorsal pallium confirmed that neurons are generated by both direct conversion from NSCs to post-mitotic neurons and asymmetric neurogenic division to generate newborn neurons or via intermediate progenitors (Barbosa et al., 2015). Interestingly, in injured telencephalon, RGs generate neural progenitors migrating toward injured sites by expanding a new type of symmetric cell division, which depletes the pool of NSCs, but generates two neural progenitor cells that are never observed under normal homeostatic conditions. Thus, compared with NSCs in mouse V/SVZ and SGZ, zebrafish dorsal pallium NSCs show unique modes of cell division in intact and injured conditions (Barbosa and Ninkovic, 2016).

In zebrafish, the process controlling NSC activation from quiescent RGs (qRGs) to proliferating RGs (pRGs) and non-glial proliferating neuroblasts is regulated by the Notch signaling pathway. In the pallium of zebrafish telencephalon, Notch 3 is prominently expressed in qRGs and pRGs, whereas Notch 1a/1b are expressed in pRGs and non-glial progenitor cells (Alunni et al., 2013; de Oliveira-Carlos et al., 2013). Indeed, Notch 3 inhibits qRGs from entering symmetric gliogenic cell division, which limits amplifying neurogenic division, whereas Notch1a stimulates qRGs to differentiate into pRGs (Alunni et al., 2013). RNA-seq analysis confirmed that Notch 3 and its downstream transcription factor Hey1 promote stemness and quiescence of qRGs in the pallium of the telencephalon (Than-Trong et al., 2018). Recently, single-cell transcriptomic analysis has been applied to NSCs in mouse brain (Basak et al., 2018; Dulken et al., 2017; Llorens-Bobadilla et al., 2015) and RGs in zebrafish telencephalon (Cosacak et al., 2019; Lange et al., 2020). scRNA-seq analysis of RGs using zebrafish transgenic lines *Tg[her4.1:mCherry; elavl3:GFP]*, which differentially labels RGs, newborn neurons, and mature neurons, revealed the diversity of RG progeny, consisting of qRGs, pRGs, newborn neurons, and mature neurons (Lange et al., 2020). This study also identified 47 genes that are differentially expressed in qRGs and pRGs. Another scRNA-seq analysis of RGs using zebrafish *Tg[her4.1:GFP]*, identified nine RG clusters. However, the diversity of these clusters is likely to represent heterogeneity of RGs corresponding to different ventricular domains of the telencephalon (Cosacak et al., 2019). Thus, it is still unknown how many cellular states can be specified in the NSC activation process from qRGs to pRGs in normal intact and injured zebrafish telencephalon.

Inflammation occurs rapidly following brain injury and involves activation of different types of immune cells. Although the immune system is beneficial for clearance of neurotoxic debris induced by brain damage, several mouse studies suggest that inflammation is largely unfavorable for neural precursor cell proliferation, which is required for tissue regeneration. Microglial activation in the brain reduces hippocampal neurogenesis, and treatment of anti-inflammatory drugs restores it (Ekdahl et al., 2003; Monje et al., 2003). Such detrimental immune responses are involved in brain damage pathogenesis and become an obstacle to recovery in mammals. However, surprisingly, in zebrafish, the immune response after traumatic brain injury is required to initiate the regenerative response of RGs in the telencephalon (Kyritsis et al., 2012), suggesting an interaction between RGs and immune cells, including microglia. However, it remains to be seen how immune cells initiate regenerative responses of RGs.

In this study, to understand how RGs respond to brain damage to initiate a regenerative response, we applied single-cell transcriptomics to RGs in zebrafish adult telencephalon using the zebrafish transgenic line *Tg[GFAP:dTomato]*. We identified five RG subtypes, which are classified into four quiescent RGs (qRG) and one proliferating RG (pRG). The four qRGs express distinct subsets of qRG markers, suggesting heterogeneity of qRGs in adult zebrafish brain. Interestingly, one qRG subtype shows high expression of ribosomal proteins, and its fraction increases in response to brain damage. Consistently, the mTOR pathway is activated in RGs near the injury site. Next, to investigate the role of microglia in regenerative responses of RGs, we eliminated microglia. We confirmed that microglial elimination suppresses damage-induced regenerative responses of RGs. Interestingly, our scRNA-seq analysis revealed that the fraction of ribosomal expression-enriched qRGs is markedly decreased in microglia-depleted injury brain, indicating that the transition of ribosomal expression-enriched qRGs depends on microglia function. Our pseudo-time analysis suggests that putative dormant RGs produce the ribosomal expression-enriched qRGs through activation of ribosomal genesis as well as suppression of cholesterol biogenesis, and pRGs through activation of the JAK/STAT pathway. Our findings reveal heterogeneity of qRGs in adult zebrafish brain and their dynamic regenerative responses to brain damage.

## Results

### TBI induces a proliferative response of RGs in the telencephalon of adult zebrafish

In zebrafish telencephalon, RGs function as neural stem cells to regenerate damaged neural tissue by re-entering cell cycle and producing neural progenitor cells (Barbosa and Ninkovic, 2016; Kishimoto et al., 2012; Kroehne et al., 2011; Than-Trong and Bally-Cuif, 2015). To understand this mechanism, stab-mediated injury was employed in the dorsal region of a hemisphere of the telencephalon of adult zebrafish, 3 – 7 months post-fertilization (mpf) (Fig. 1A). After BrdU treatment for 12 h, injured fish were sacrificed at 1, 3, 7 and 14 dpl, and the telencephalon was dissected and fixed with paraformaldehyde (PFA) (Fig. 1B). Using a vibratome, 100-μm slices covering the telencephalon along the anterior and posterior axes were prepared and labeled with anti-GFAP and ani-BrdU antibodies. GFAP is a glial maker and is expressed in all RGs of the zebrafish telencephalon (Lam et al., 2009). RG cell bodies are located in the ventricular zone of the telencephalon and extend processes to the basal area of the brain (Ganz et al., 2010; Marz et al., 2010) (Fig. 1C). Double labeling of the telencephalon with anti-BrdU and anti-GFAP antibodies visualized proliferating RGs, which re-entered cell-cycle progression (Fig. 1D). We examined BrdU^+^ RGs in telencephalic slices at 1, 3, 7, and 14 dpl (Fig. 1E) and counted the total number of BrdU^+^ RGs (Fig. 1F). The number of BrdU^+^ RGs was significantly higher at 3 dpl than that of non-injured controls. However, there was no significant difference at other time points between injured samples and non-injured controls. We also confirmed that the number of BrdU^+^ RGs was significantly higher in lesioned hemispheres than in control intact hemispheres (Fig. 1G). These data suggest that cell proliferation is activated in RGs at 3 dpl, which is consistent with previous reports (Kishimoto et al., 2012; Kroehne et al., 2011). The telencephalon is divided into two major areas, the dorsal telencephalic area (D) and the ventral telencephalic area (V) (Ganz et al., 2010; Than-Trong and Bally-Cuif, 2015) (Fig. 1C). Accordingly, the ventricular zone of D is divided into three areas, the medial (Dm), dorsal (Dd), and lateral zones of D (Dl). Similarly, the ventricular zone of V is divided into two areas, the dorsal nucleus (Vd) and the ventral nucleus of V (Vv). We examined which ventricular zone area most actively responds to TBI. RGs in the Dd region responded significantly to TBI and incorporated BrdU, whereas RGs in other areas did not (Fig. 1H). This is reasonable, because the Dd region is closest to the site of the administered stab injury. Taken together, TBI induces a regenerative response of RGs in adult zebrafish telencephalon.

**Figure 1.**
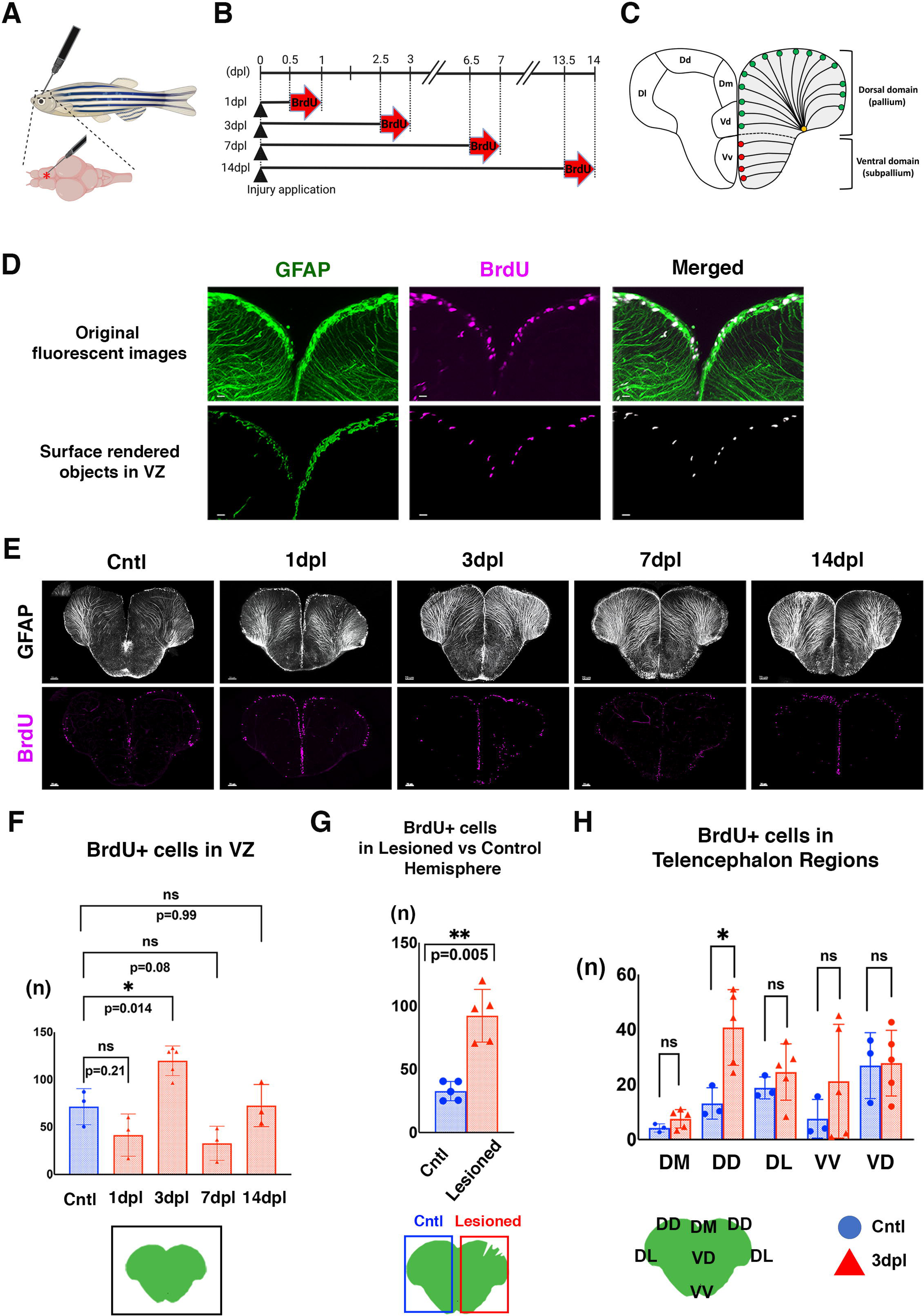
Stab injuries activate regenerative responses of RGs in adult zebrafish telencephalon. (A) Injury application method. Using a needle, stab injury is introduced in the right hemisphere of the dorsal telencephalon (pallium) in adult zebrafish. Image was created using Biorender (https://biorender.com). (B) Experimental design to investigate temporal profiles of regenerative responses of RGs to TBI. After TBI, BrdU is applied for 12 h prior to analysis at 1, 3, 7, and 14 dpl. Image was created using Biorender (https://biorender.com). (C) Schematic drawing of zebrafish adult telencephalon. In zebrafish, two distinct neurogenic proliferation zones were formed in ventricular surfaces of the dorsal and ventral telencephalon, respectively. In the dorsal domain, called the pallium, an RG cell body (green circle) is located at the ventricular surface and extends a long radial process, the endfoot of which reaches the pial surface close to a blood vessel (orange circle) at the lateral margin of the dorsal telencephalon. In the ventral domain, called the subpallium, neuroepithelial-like progenitors (red circles) are located at the ventricular zone. (D) Double labeling of the telencephalon with anti-GFAP (green) and anti-BrdU (magenta) antibodies. Top and bottom panels indicate original fluorescent images and surface-rendered object images, respectively. Only surface-rendered objects located in the ventricular zone (VZ) were used for analysis. Scales: 20μm. (E) Double labeling of the telencephalon with anti-GFAP (white) and anti-BrdU (magenta) antibodies at 1, 3, 7, and 14 dpl. The left-most panel indicates non-injured telencephalon. Scales: 70μm. (F) The number of BrdU+ cells in the ventricular zone of both hemispheres. The number is significantly higher at 3 dpl than in non-injured control. There is no significant difference between non-injured controls and other time points. mean±SD. p*<0.05; one way ANOVA Dunnett’s multiple comparison. (G) Comparison of the number of BrdU^+^ cells between lesioned and intact hemispheres at 3 dpl. The number was significantly higher in lesioned hemispheres. mean±SD. p**<0.01; paired t-test. (H) The number of BrdU^+^ cells in different ventricular domains of the telencephalon: Dm, Dd, Dl, Vv and Vd domains. The number was significantly higher in the Dd domain at 3 dpl than in non-injured control brains. There was no significant difference in other domains between 3 dpl and non-injured controls, although the number was higher in Dm, Dl and Vv at 3 dpl. mean±SD. p*<0.05; unpaired students’ t-test. **Source Data 1.** Data for Figure 1FGH.

### scRNA-seq analysis on non-injured controls and 3-dpl RGs revealed five distinct populations of RGs

To identify the mechanism that regulates the regenerative response of RGs, we applied scRNA-seq analysis to RGs in 3-dpl and non-injured control telencephalons (Fig. 2A). Stab injuries were inflicted in telencephalons of 5-mpf adult zebrafish carrying a transgene *Tg[gfap:dTomato]*, which expresses dimeric Tomato (dTomato) in RGs under the control of the *gfap* promoter (Satou et al., 2012; Than-Trong et al., 2020). At 3 dpl, telencephalons were homogenized with papain to prepare cell suspensions, which were subsequently subjected to flow cytometry-assisted cell sorting (Fig. 2-figure supplement 1AC). We used a 10x genomic system to prepare scRNA-seq libraries. After read sequencing was done, reads were mapped to the zebrafish genomic database GRCz11. Next, we evaluated nUMI, nGene, and mitochondrial genes in both datasets separately to ensure that all data were comparable (Fig. 2-figure supplement 1D). After removing cells that exhibited unique gene counts below the minimum threshold (200 genes) or above the maximum threshold (2500 genes), or mitochondrial genes above the maximum threshold (5% counts), we obtained 567 and 747 cells from non-injured control and 3-dpl samples, respectively. We conducted unbiased clustering of the datasets from non-injured control and 3-dpl samples separately, using the Seurat package (Butler et al., 2018). In the non-injured control sample, cells were classified into 6 clusters (Fig. 2-figure supplement 2A). Based on a previous report on zebrafish genes specifically expressed in qRGs, pRGs, neuroblasts, newborn neurons (NBNs), ependymal cells, and oligodendrocyte precursor cells (OPCs) (Lange et al., 2020), 6 clusters corresponded to 3 types of qRGs, pRG/Neuroblast/NBN, ependymal cells/OPC, and probably pineal gland (Fig. 2-figure supplement 2B, Supplementary table 1). On the other hand, in the 3-dpl sample, cells were classified into 7 clusters (Fig. 2-figure supplement 3A), which correspond to 4 types of qRGs, qRG/OPC, pRG/Neuroblast/NBN, ependymal cells (Fig. 2-figure supplement 3B, Supplementary table 2). Thus, each datasets had sufficient quality to apply further clustering analysis of merged datasets.

**Figure 2.**
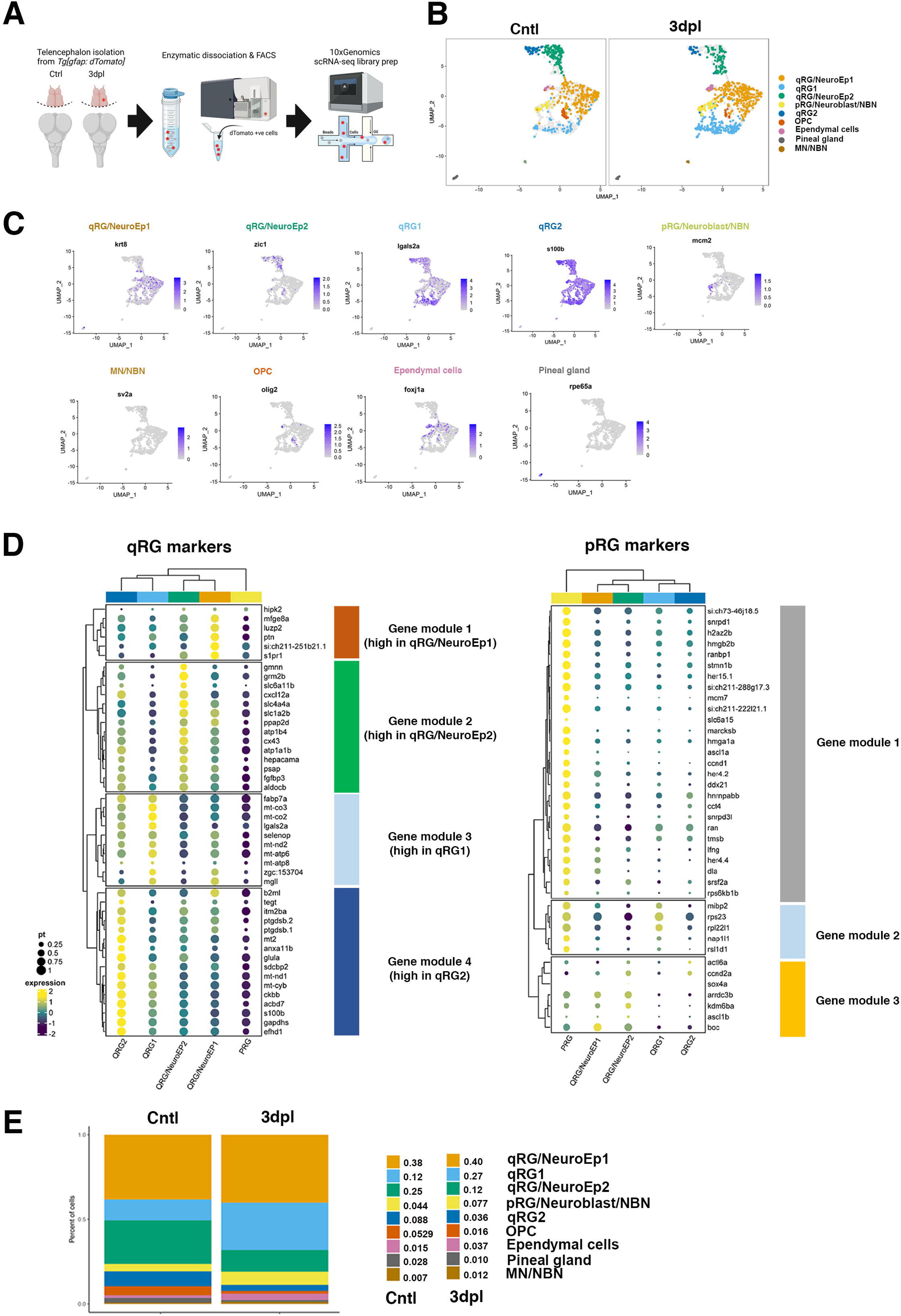
scRNA-seq analysis identifies 4 distinct qRG clusters in 3-dpl and non-injured control telencephalons. (A) Experimental procedures of scRNA-seq analysis of *GFAP:dTomato*^+^ cells in adult telencephalons in 3-dpl and non-injured control conditions. Image was created using Biorender (https://biorender.com). (B) UMAP of scRNA-seq data. Nine clusters are identified: 4 qRG, pRG/Neuroblast/NBN, OPC, Ependymal cells, Pineal gland, MN/NBN. (C) Feature plot of markers defining each of the 9 clusters. (D) Dot plot analysis of 46 qRG (left) and 39 pRG (right) markers in 4 qRG (qRG/NeuroEp1, qRG/NeuroEp2, qRG1, and qRG2) and pRG. qRG and pRG markers are classified into 4 and 3 Gene modules, respectively. (E) Fraction of each cluster in 3-dpl and non-injured control conditions. qRG1 and pRG fractions were increased at 3 dpl, whereas qRG/NeuroEp2 and qRG2 fractions were decreased at 3 dpl. **Figure 2-figure supplement 1.** FACS cell sorting and scRNA-seq data quality check. **Figure2-figure supplement 2.** Cell clusters in the non-injured control dataset. **Figure2-figure supplement 3.** Cell clusters in the 3-dpl dataset. **Figure2-figure supplement 4.** Heatmap of the top 10 markers for clusters in the non-injured control and 3-dpl merged datasets. **Figure2-figure supplement 5.** Gene module analysis of qRG markers. **Figure2-figure supplement 6.** Regional marker analysis of qRGs. **Figure2-figure supplement 7.** Expression analysis of genes responsible for quiescence and stemness of qRGs. **Figure2-figure supplement 8.** Gene module analysis of pRG markers.

Next, we utilized the Harmony merging algorithm and applied clustering analysis to the two merged datasets, non-injured controls and 3-dpl samples (Korsunsky et al., 2019). gfap:dTomato^+^ cells were classified into 9 clusters, corresponding to 4 types of qRGs, pRG/Neuroblast/MBN, MN/NBN, OPC, ependymal cells, and pineal gland (Fig. 2B). Heat map analysis of TOP10 genes differentially expressed in each cluster confirmed clear separation of each cluster (Fig.2-figure supplement 4). We also confirmed how each of the 9 clusters of the merged datasets corresponded to 6 clusters of non-injured control cells (Fig.2-figure supplement 2C) and 7 clusters of 3-dpl cells (Fig.2-figure supplement 3C).

### Four qRGs differentially express a distinct subset of qRG-specific genes

4 qRG clusters express qRG markers, and two of them also express neuroepithelial markers, *krt8* and *zic1*, respectively (Fig. 2C). So, we named these two clusters qRG/NeuroEp1 and qRG/NeuroEp2. In other two qRG clusters, one expressed *lgals2a* and the other expressed *s100β* in TOP10 genes (Fig.2-figure supplement 4), which are named qRG1 and qRG2, respectively (Fig. 2C). To further characterize the 4 qRG clusters, we focused on 47 genes that were reported to be specifically expressed in ccnd1-negative qRG (Lange et al., 2020). Among these 47 genes, only one gene, *drg3a*, was not included for analysis. Interestingly, the other 46 genes were differentially expressed in each of the 4 qRG clusters (Fig. 2D). These data suggest that qRG/NeuroEp1, qRG/NeuroEp2, qRG1 and qRG2 correspond to different states of qRGs.

Interestingly, these 46 qRG markers are separated into 4 gene modules, each of which were highly expressed in qRG/NeuroEp1, qRG/NeuroEp2, qRG1 and qRG2 (Fig. 2-figure supplement 5A). Gene module 1 contains 6 qRG genes: *hipk2*, *mfge8a*, *luzp2*, *ptn*, *si:ch211-251b21.1*, and *s1pr1*, that are prominently expressed in qRG/NeuroEp1 (Fig. 2-figure supplement 5BC). Gene module 2 contains 14 qRG genes: *gmnn*, *grm2b, slc6a11b, cxcl12a, slc4a4a, slc1a2b, ppap2d, atp1b4, cx43, atp1a1b, hepacama, psap, fgfbp3,* and *aldocb*, that are prominently expressed in qRG/NeuroEp2 (Fig. 2-figure supplement 5D). Gene module 3 contains 10 qRG genes: *fabp7a, mt-co3, mt-co2, lgals2a, selenop, mt-nd2, mt-atp6, mt-atp8, zgc:153704,* and *mgll*, that are prominently expressed in qRG1 (Fig. 2-figure supplement 5E). Gene module 4 contains 16 qRG genes: *b2ml, tegt, itm2ba, ptgdsb.2, ptgdsb.1, mt2, anxa11b, glula, sdcbp2, mt-nd1, mt-cyb, ckbb, acbd7, s100b, gapdhs,* and *efhd1*, that are prominently expressed in qRG2 (Fig. 2-figure supplement 5F).

Previous scRNA-seq analysis of zebrafish telencephalon revealed that heterogeneity of qRG clusters mostly depends on telencephalic ventricular domains, where each qRG subtype is positioned (Cosacak et al., 2019). Accordingly, we examined whether these 4 qRG clusters are linked to different telencephalic ventricular domains. We focused on 8 telencephalic ventricular regional markers used in Cosacak et al. (2019): *foxp4* (Vd), *Pou3f1* (Dd, Dm, Dl), *dmrta2* (Dd, Dm), *gsx2* (Dm), *nr2f1b* (Dd), *foxj1a* (Vd), *zic1* (Vv), and *iqgap2* (pre-neuroblasts/neuroblasts), and examined their expression in 4 qRG clusters (Fig. 2-figure supplement 6A). *zic1* was specifically expressed in qRG/NeuroEp2 and qRG2 (Fig. 2-figure supplement 6BC). *gsx2* was prominently expressed in qRG/NeuroEp2 and qRG2, but weakly expressed in a relatively small fraction of qRG/NeuroEp1 (Fig. 2-figure supplement 6BC). *foxj1a* was specifically expressed in qRG/NeuroEp1 (Fig. 2-figure supplement 6BC). *nr2f1b* was expressed in a small fraction of qRG1, whereas *pou3f1, dmrta2*, and *foxp4* were expressed in a small fraction of qRG/NeuroEp1 and pRG/Neuroblast/NBN (Fig. 2-figure supplement 6BC). These data suggest that qRG/NeuroEp1, qRG1 and pRG/Neuroblast/NBN are expressed in the dorsal ventricular zone (Dm, Dd, and Dl) as well as Vd, whereas qRG/NeuroEp2 and qRG2 are expressed in Dm and Vv. These data suggest that qRG/NeuroEp1, qRG1 and pRG/Neuroblast/NBN link each other in terms of ventricular domains, especially Vd, Dm, Dd, and Dl. This is consistent with our observation that RG proliferation is significantly increased in the Dd area after TBI (Fig. 1H).

In zebrafish telencephalon, Notch3 and its downstream target, Hey1, are essential for maintenance of quiescence and stemness of RGs (Than-Trong et al., 2018). *gfap*, *her4.1, her6, her9*, *sox2,* and *nestin* are expressed in RGs (Than-Trong and Bally-Cuif, 2015; Than-Trong et al., 2020). *vimentin* is strongly expressed in RGs of the dorsal ventricular domain and Vd, but weakly expressed in Vv (Ganz et al., 2010). The JNK-Jun pathway mediates stress responses after neuronal injury (Coffey, 2014). Therefore, we examined expression of these genes in 4 qRG and pRG clusters. Interestingly, these genes are classified into 4 modules (Fig. 2-figure supplement 7A). *notch3, hey1, sox2* and *her6* are expressed in qRG/NeuroEp1, qRG/NeuroEp2, and qRG2; however, their expression is low in qRG1 and pRG (Fig. 2-figure supplement 7BC), suggesting that stemness and quiescence start to be lost in qRG1. *vimentin, nestin, her4.1* are expressed very little in qRG/NeuroEp2 and qRG2, whereas they are highly expressed in qRG/NeuroEp1, qRG1, pRG (Fig. 2-figure supplement 7BC), suggesting that regenerative responses start in qRG/NeuroEp1, qRG1 and pRG. Furthermore, a low level of *vimentin* expression is consistent with our regional marker analysis, indicating that qRG/NeuroEp2 and qRG2 have a more ventral subpallium character (Fig. 2-figure supplement 6). Interestingly, *jun* is expressed in qRG/NeuroEp1, qRG/NeuroEp2, pRG, but little expressed in qRG1 and qRG2 (Fig. 2-figure supplement 7BC), suggesting that qRG1 is not exposed to a stress response.

Next, we examined pRG markers in 4 qRG clusters. Here we focused on 47 genes that were reported to be specifically expressed in ccnd1-positive pRGs (Lange et al., 2020). Among these 47 pRG genes, expression of eight genes (*si:ch211-193l2.3, si:ch211-193l2.4, si:ch211-193l2.5, si:ch211-193l2.6, si:ch211156b7.4, prrc2b, tcp1*, and *h3f3a*) was not detected for pRG analysis. Thus, we investigated expression of the remaining 39 pRG-specific genes in 4 qRG clusters and pRG (Fig. 2D). These 39 genes are classified into three groups, Gene modules 1-3, depending on their expression profile in qRG clusters (Fig. 2-figure supplement 8A). Gene module 1 contains 27 genes that are highly expressed in pRG, but little expressed in all qRG clusters (Fig. 2-figure supplement 8BC). Gene module 2 contains 5 genes (*mibp2, rps23, rpl22l2, nap1l1,* and *rsl1d1*) that are highly expressed in both pRG and qRG1 (Fig. 2-figure supplement 8D). Nap1l1 is a nucleosome assembly protein 1-like 1 and promotes cell proliferation of neuronal progenitors and neurogenesis (Qiao et al., 2018). Rsl1d1 is a ribosomal L1 domain-containing 1 that directly interacts with Mdm2 to promote tp53 degradation (Ding et al., 2021). In addition, increased expression of ribosomal proteins such as *rps23* and *rpl22l2* indicates energy metabolism in qRG1. Thus, these genes may promote cell proliferation and survival in qRG1. Gene module 3 contains 7 genes (*actl6, ccnd2a, sox4a, arrdc3b, kdm6ba, ascl1b,* and *boc*) that are highly expressed in pRG, qRG/NeuroEp1, and qRG/NeuroEp2 (Fig. 2-figure supplement 8E). In zebrafish, Kdm6ba is a histone demethylase JMJB3 and is essential for tissue regeneration (Stewart et al., 2009). ascl1 is a master regulator of neuronal regeneration (Ramachandran et al., 2011). So, elevated expression of *kdm6ba* and *ascl1b* may indicate that a regenerative response starts to be activated in qRG/NeuroEp1.

### Each RG fraction is dynamically changed in response to TBI

We compared the fraction of each cluster in non-injured controls and 3-dpl conditions (Fig. 2E). First, the fraction of the pRG/Neuroblast/NBN cluster was increased in 3-dpl conditions (7.7%), compared with non-injured controls (4.4%). Accordingly, the fraction of MN/NBN was also increased in in 3-dpl conditions (1.2%), compared with non-injured controls (0.7%). This is consistent with our observation that a higher number of RGs enter cell proliferation to regenerate neurons in response to TBI at 3 dpl (Fig. 1E). Another drastic increase was detected in the fraction of qRG1 in 3-dpl conditions (27.0%), compared with that of non-injured controls (12.0%) (Fig. 2E). On the other hand, the fraction of qRG/Ep2 was decreased in 3-dpl conditions (12.0%), compared with non-injured controls (25.0%). The fraction of qRG2 was also decreased in 3-dpl conditions (3.6%), compared with non-injured controls (8.8%). Since qRG1 seems to be spatially separated from qRG2 and qRG/NeuroEp2 (Fig. 2-figure supplement 6), the reduction of the qRG2 and qRG/NeuroEp2 fractions may result from the increase of the qRG1 and pRG/Neuroblast/NBN fractions. The qRG/NeuroEP1 fraction was not changed between 3-dpl conditions (40.0%) and non-injured controls (38.0%). In summary, in response to TBI, qRG2 and qRG/Ep2 were decreased, whereas qRG1, pRG/Neuroblast/NBN, MN/NBN were increased.

### qRG1 is a ribosomal protein-enriched RG that increased in response to TBI

Since about 70% of the TOP100 differentially expressed genes of qRG1 are ribosomal proteins (Supplemental Table 3), qRG1 shows a high expression of ribosomal proteins, which is shared with pRG, but not with other qRG populations (Fig. 2-figure supplement 8D). In addition, qRG1 does not express Notch3, but expresses *her4.1*, and *vimentin* (Fig.2-figure supplement 7). Since Notch3 is required to maintain quiescence and stemness of NSCs (Alunni et al., 2013; Than-Trong et al., 2020), these data suggest that qRG1 is in transition from a dormant qRG state. Furthermore, previous scRNA-seq analyses using adult mouse brain revealed that dormant NSCs enter a primed-quiescent state before activation, which involves downregulation of Notch signaling and concomitant upregulation of ribosomal protein expression, leading to elevated protein synthesis (Basak et al., 2018; Dulken et al., 2017; Llorens-Bobadilla et al., 2015). Thus, these primed quiescent NSCs in mice have a similar transcriptomic profile to qRG1 in zebrafish. Since the qRG1 fraction increases in response to TBI, it is expected that protein translation by upregulated ribosomal genesis is elevated in TBI.

To confirm this possibility, we examined whether the mammalian target of rapamycin (mTOR) signaling is activated in telencephalon in response to TBI. mTOR signaling is important for stem cell self-renewal and differentiation in mammalian brain development (Meng et al., 2018; Saxton and Sabatini, 2017). During activation of mTOR signaling, mammalian TOR complex 1 (mTORC1) activates ribosomal protein S6 kinase (S6K) (Magnuson et al., 2012), which subsequently induces phosphorylation of ribosomal protein S6 (rpS6), a component of the S40 ribosome (Ruvinsky and Meyuhas, 2006). Furthermore, rpS6 phosphorylation by S6K is required for regeneration of zebrafish fin (Hirose et al., 2014), heart (Miklas et al., 2022), and retinal pigment epithelium (Lu et al., 2022). Thus, we conducted labeling of zebrafish telencephalon at 3 dpl with anti-phospho-rpS6 antibody. Phosphorylation of rpS6 was increased in the dorsal ventricular zone (Dm, Dd, and Dl) of lesioned hemispheres, compared with non-injured control hemispheres (Fig. 3A). The fraction of p-rpS6^+^ area in GFAP^+^ area is significantly higher in lesioned hemispheres than in intact non-injured hemispheres (Fig. 3B), suggesting that the mTOR-activated population of RGs is increased by TBI. These data suggest that mTOR signaling is activated in response to TBI in zebrafish telencephalon.

**Figure 3.**
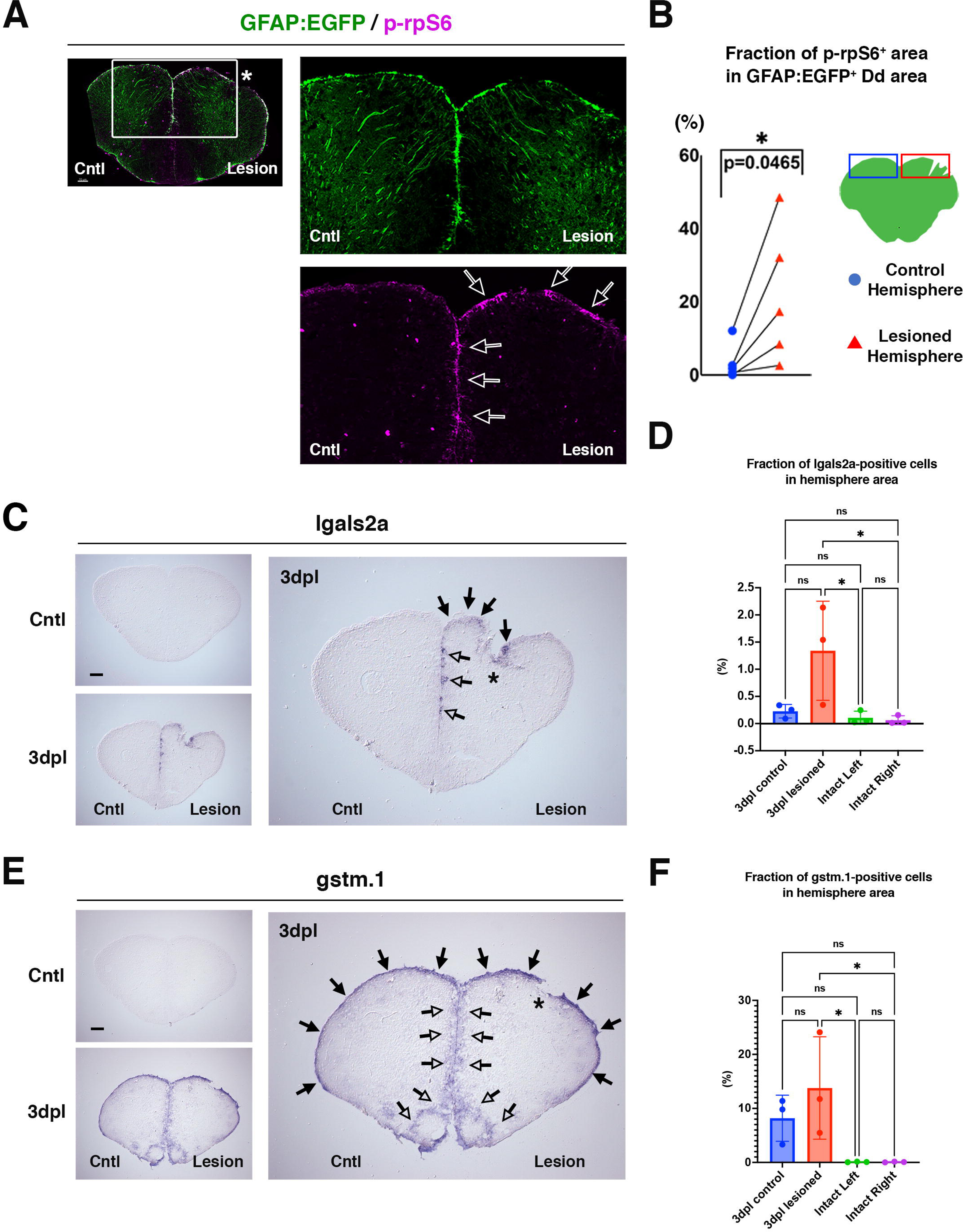
TBI promotes the mTOR signaling pathway and transcription of *lgals2a* and *gstm.1* in the ventricular zone of the pallium. (A) Labeling of 3-dpl *Tg[GFAP:EGFP]* (green) transgenic telencephalon with anti-p-rpS6 antibody (magenta). Right panels show higher magnification images. Arrowheads indicate p-rpS6^+^ RG, which are increased in the injured hemisphere (right). Scale: 70μm. (B) Fraction of p-rpS6^+^ area in the GFAP:EGFP^+^ Dd area in non-injured control and lesioned hemispheres. The fraction of p-rpS6^+^ area is significantly higher in lesioned hemispheres than in non-injured control hemispheres. p*<0.05; paired t-test. (C) *In situ* hybridization of adult zebrafish telencephalon with the *lgals2a* RNA probe in non-injured control and 3-dpl conditions. The right panel shows a higher-magnification image of a 3-dpl sample. An asterisk indicates the stab-induced injury site. Black filled and open arrowheads indicate mRNA expression in Dd and Dm ventricular domains, respectively. Scale: 100 μm. (D) The fraction of *lgals2a*^+^ cells in the hemisphere area. This fraction is markedly higher in 3-dpl lesioned hemispheres. mean±SD. p*<0.05; One way ANOVA Turkey’s multiple comparison test. (E) *In situ* hybridization of adult zebrafish telencephalon with the *gstm.1* RNA probe in non-injured control and 3-dpl conditions. The right panel shows a higher-magnification image of a 3-dpl sample. An asterisk indicates the stab-induced injury site. Black filled and open arrowheads indicate mRNA expression in Dd and Dm/Vd/Vv ventricular domains, respectively. Scale: 100 μm. (F) The fraction of *gstm.1*+ cells in the hemisphere area. mean±SD. The fraction is higher in 3-dpl control and lesioned hemispheres. p*<0.05; One way ANOVA Turkey’s multiple comparison test. **Source Data 1.** Data for Figure 3B. **Source Data 2.** Data for Figure 3DF.

Next, we examined whether qRG1 cells increase in response to TBI in zebrafish telencephalon *in vivo*. One of the qRG markers, *lgals2a*, is strongly expressed in qRG1 and moderately expressed in qRG2, but very low in qRG/Ep1 and qRG/Ep2 (Fig. 2D and Fig.2-figure supplement 5AE). *lgals2a* expression was upregulated in the dorsal ventricular zone (Dd) of lesioned hemispheres, compared with non-injured control hemispheres, in which *lgals2a* expression is very low (Fig. 3CD). It was reported that *gstm.1* is expressed in primed quiescent neural stem cells in mice (Basak et al., 2018). scRNA-seq analysis revealed that *gstm.1* is one of the differentially expressed genes of qRG1 (Supplementary Table 3). *gstm.1* expression was highly upregulated in RGs of both lesioned hemispheres and non-injured hemispheres in TBI-induced brain, compared with non-injured control brains, in which *gstm.1* expression is very low (Fig. 3EF). These data suggest that ribosomal protein enriched qRG1 is increased in the ventricular zone of the dorsal telencephalon in response to TBI.

### Pseudo-time analysis reveals that qRG1 and pRG are generated from qRG/NeuroEp1

qRG1 in zebrafish and primed quiescent NSCs in mice share upregulation of ribosomal proteins and downregulation of Notch signaling (Dulken et al., 2017; Llorens-Bobadilla et al., 2015), suggesting that qRG1 may be in a transient state from dormant qRG toward pRG. To examine this possibility, we conducted pseudo-time analysis using Monocle3 (Trapnell et al., 2014). We determined qRG/NeuroEp1 as the starting point for pseudo-time analysis, because qRG/NeuroEp1 expresses dormant markers of qRG such as *notch3* (Alunni et al., 2013; Than-Trong et al., 2018) (Fig. 2-figure supplement 7AC) and *mfge8a* (Zhou et al., 2018) (Fig. 2-figure supplement 5AC). In addition, two qRG markers highly expressed in qRG/NeuroEp1, *si:ch211-251b21.1* and *s1pr1* (Fig. 2-figure supplement 5AC), are downregulated in *Notch3^-/-^*mutants (Than-Trong et al., 2018), suggesting that Notch3 promotes expression of these qRG markers in qRG/NeuroEp1. Our pseudo-time transcriptome analysis revealed that qRG/NeuroEp1 was split into three branches in non-injured controls, which lead to pRG/Neuroblast/NBN, qRG1, and qRG/NeuroEp2, respectively (Fig. 4A and Fig. 4-figure supplement 1A). On the other hand, in 3-dpl conditions, qRG/NeuroEp1 was split into two branches, which lead to pRG/Neuroblast/NBN and qRG1, respectively, and qRG/NeuroEp2 was not mapped on either pseudo-time branches (Fig. 4A and Fig. 4-figure supplement 1A). Gene expression dynamics on UMAP (Fig. 4-figure supplement 1B) and a Dot-plotting graph (Fig. 4B) confirmed different gene expression dynamics between a branch moving toward qRG1 (Branch1) and a branch moving toward pRG/Neuroblast/NBN (Branch2). These data suggest that qRG1 is not an intermediate state from qRG/NeuroEp1 into pRG/Neuroblast/NBN, but that qRG1 and pRG are produced via two separate lineages from qRG/NeuroEp1. However, Monocle3 UMAP indicates that a small population of pRG/Neuroblast/NBN cells are associated with a Branch1 trajectory path and spatially segregated with a dense population of pRG/Neuroblast/NBN cells mapped on a Branch2 trajectory path, suggesting the possibility that qRG1 cells change into pRGs/Neuroblasts/NBNs (Fig. 4-figure supplement 2A). To clarify this inconsistency, we extracted both Branch1- and Branch2-associated pRG/Neuroblast/NBN cells and examined their gene expression profiles. We found that differentially expressed genes differ completely between Branch1- and Branch2-associated pRG/Neuroblast/NBN cells. Branch1-associated pRG/Neuroblast/NBN cells showed a very similar gene expression profile to qRG1, indicating that Branch1-associated pRG/Neuroblast/NBN cells are qRG1 (Fig. 4-figure supplement 2BC). These data suggest that qRG/NeuroEp1 is a source of both qRG1 and pRG/Neuroblast/NBN, and that these two RG types are differentially generated from qRG/NeuroEp1.

**Figure 4.**
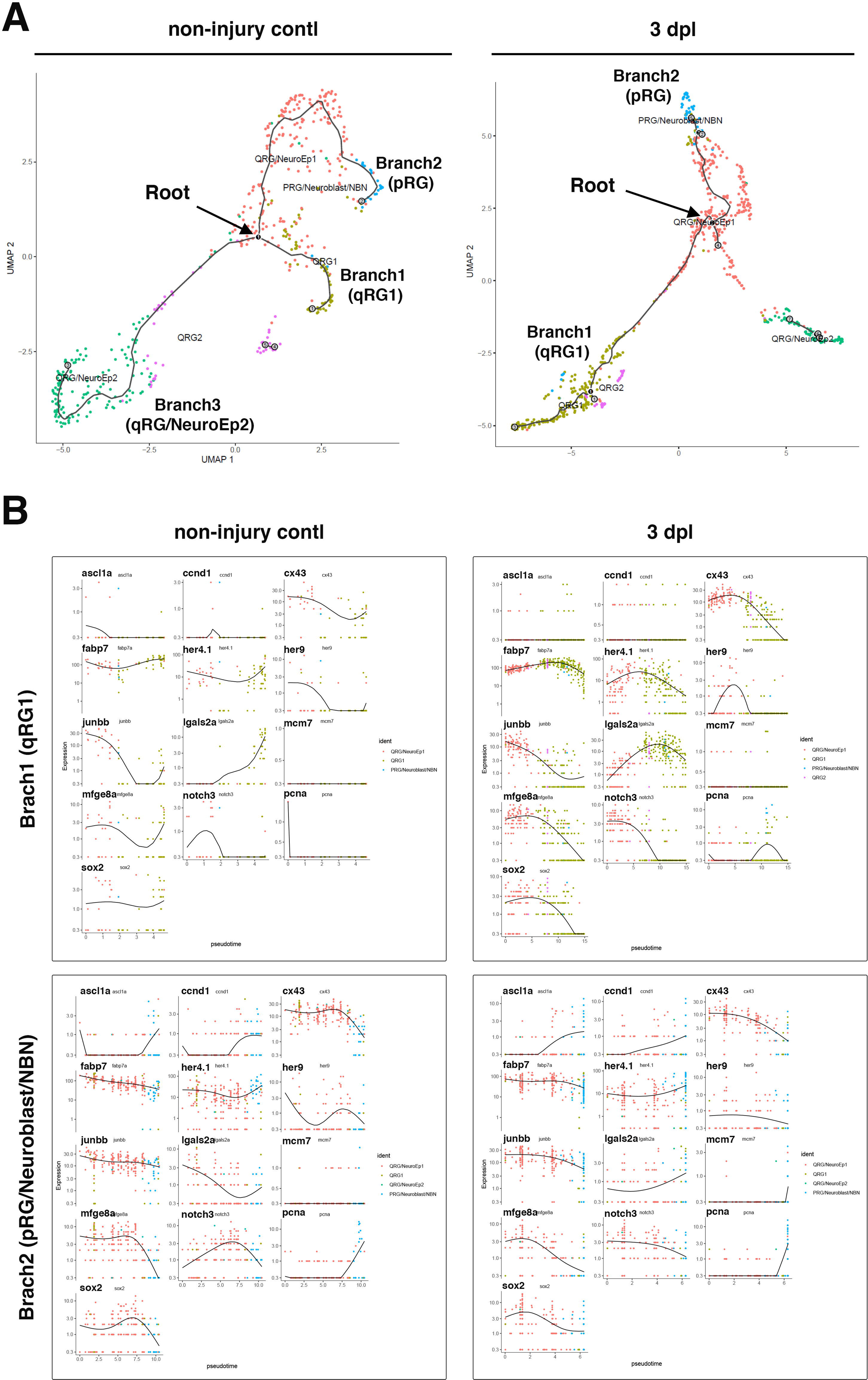
Pseudo-time analysis indicates that qRG/NeuroEp1 generates qRG1 and pRG in separate lineages. (A) Pseudo-time trajectory generated with monocle 3 for non-injured control (left) and 3-dpl conditions (right). qRG/NeuroEp1 is selected as a root for pseudo-time trajectory. Three branches are specified in non-injured control conditions, whereas two branches are specified in 3-dpl conditions. Trajectories toward qRG1 and pRG are observed as Branch1 and Branch2, respectively, in both conditions. (B) Gene expression profile along pseudo-time trajectory of branch1(qRG1) and branch2 (pRG) in non-injured control and 3-dpl conditions. Thirteen genes are selected: *ascl1a*, *ccnd1*, *cx43*, *fabp7*, *her4.1*, *her9*, *junbb*, *lgals2a*, *mcm7*, *mfge8*, *notch3*, *pcna*, and *sox2*. **Figure 4-figure supplement 1.** Pseudo-time analysis using Monocle3 for non-injured control and 3-dpl conditions. **Figure 4-figure supplement 2.** Gene expression analysis of branch1- and branch2-associated pRG/Neuroblasts/NBN cells. **Figure 4-figure supplement 3.** AhR signaling pathway is specifically activated in qRG/NeuorEp2.

Interestingly, the trajectory path from qRG/NeuroEp1 to qRG/NeuroEp2 disappeared in TBI conditions (Fig. 4A). The qRG1 population is increased by TBI, whereas qRG/NeuorEp2 is decreased by it (Fig. 2E). In adult zebrafish pallium, 17% of RGs produce neurons by direct conversion; however, the percentage of RGs undergoing direct conversion was markedly deceased to 3 % after TBI (Barbosa et al., 2015). Recently it was reported that high aryl hydrocarbon receptor (AhR) signaling promotes the direct conversion of a specific subset of RGs into post-mitotic neurons, while low AhR signaling promotes proliferation of RGs (Di Giaimo et al., 2018). After brain damage, AhR signaling is promptly suppressed in 1–5 dpl and recovered to a normal level at 7 dpl in zebrafish, indicating that brain damage transiently inhibits AhR signaling, leading to decrease of the rate of direct conversion. We found that a major mediator of AhR signaling in zebrafish, Ahr2, and its transcriptional target, cytochrome P450 1B1 oxidase (Cyp1b1), are highly expressed in qRG/NeuroEp2 but not in other qRGs (Fig. 4-figure supplement 3). qRG/NeuroEp2 cells are located in Dm and Vv (Fig.2-figure supplement 7). These data raise the possibility that Dm-associated qRG/NeuroEp2 is a major source of direct conversion, which may explain the disappearance of the trajectory path from qRG/NeuroEp1 to qRG/NeuroEp2 after TBI as well as decrease of qRG/NeuorEp2 fraction after TBI. Further *in vivo* analysis will be necessary to clarify this point.

### TBI dynamically modifies regulatory gene expression in the transition from qRG/NeuroEp1 to qRG1 and pRG

To determine how TBI promotes the transition from qRG/NeuroEp1 to qRG1 and pRG, we determined upregulated and downregulated genes in qRG/NeuroEp1 in the injured condition, compared with the non-injured control condition (Fig. 5A). First, we selected genes whose mRNA expression was significantly upregulated in 3-dpl conditions, compared with non-injured control conditions, and applied gene ontogeny (GO) analysis (Fig. 5B). These upregulated genes are classified into several regulatory pathways (Fig. 5-figure supplement 1A), for example, the JAK-STAT pathway (Fig. 5 - figure supplement 1BC), innate immune response (Fig. 5-figure supplement 1D), and ribosomal metabolism including translation, ribosomal biogenesis, ribosome assembly and cell-cycle regulation (Fig. 5-figure supplement 1E).

**Figure 5.**
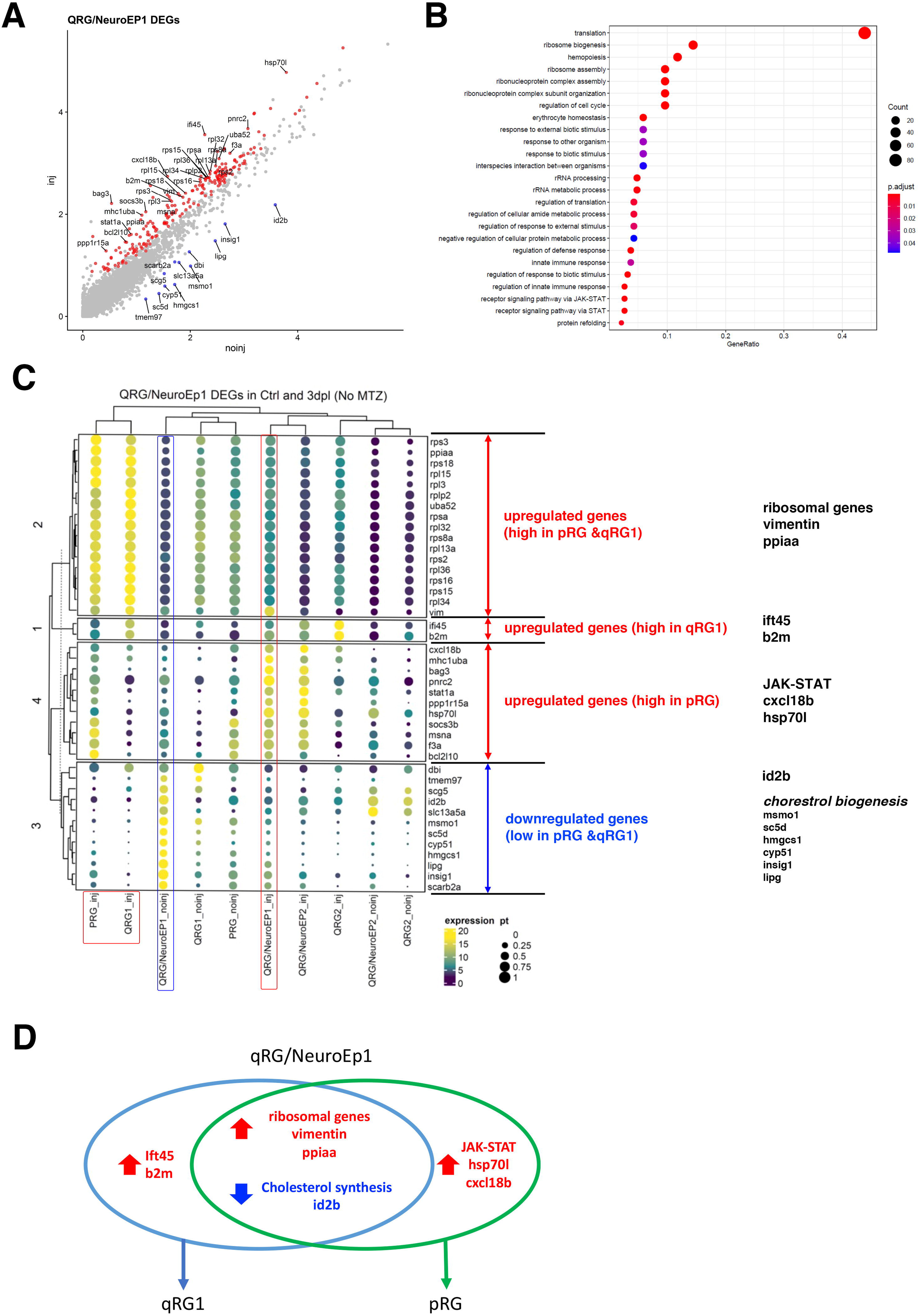
DEG analysis of qRG/NeuroEp1 in response to TBI. (A) Scatter plot visualization of differentially expressed genes (DEGs) in qRG/NeuroEp1 in response to TBI. Upregulated (red) and downregulated (blue) genes are mapped. (B) GO analysis of DEGs of qRG/NeuroEp1 in response to TBI. (C) Dot-plot visualization of up- and downregulated genes of qRG/NeuroEp1 in response to TBI across all RG clusters in non-injured control and 3-dpl conditions. TOP30 upregulated genes (red) and TOP12 downregulated genes (blue) are shown. Upregulated genes are classified into three groups: (1) genes whose expression is high in both qRG1 and pRG at 3 dpl, (2) genes whose expression is high in qRG1 at 3 dpl, and (3) genes whose expression is high in pRG at 3 dpl. All downregulated genes show low expression in both qRG1 and pRG at 3 dpl. Genes nominated by GO analysis are shown in the right-most column. (D) Venn diagram of up- and downregulated genes in qRG/NeuroEp1 linked by the transition to qRG1 and pRG/Neuroblast/NBN. **Figure 5-figure supplement 1.** Signaling network of upregulated genes in qRG/NeuroEp1 in response to TBI. **Figure 5-figure supplement 2.** Gene expression profiles of the TOP30 upregulated genes in qRG/neuroEp1 along a pseudo-time trajectory. **Figure 5-figure supplement 3.** Signaling network of downregulated genes in qRG/NeuroEp1 in response to TBI. **Figure 5-figure supplement 4.** Gene expression profiles of the TOP12 downregulated genes in qRG/neuroEp1 along a pseudo-time trajectory.

Next, among those, the TOP30 upregulated genes were selected. We examined mRNA expression levels of these upregulated genes in each of 4 qRG clusters and the pRG cluster between injured and non-injured conditions (Fig. 5C). Interesting, the TOP30 upregulated genes are classified into three groups. The first group has 17 genes, which are highly expressed in both qRG1 and pRG clusters in the injured condition (Fig. 5C). Most of these genes are ribosomal genes, suggesting that ribosomal biogenesis is enhanced in the transition lineage from qRG/NeuroEp1 to both qRG1 and pRG. In addition, protein folding regulator *ppiaa* and *vimentin* are found in this first group. The second group has 2 genes, *ift45* and *b2m*, whose mRNA expression is higher in qRG1 than in pRG (Fig. 5C), suggesting that these two genes may promote the transition to qRG1. The third group has 11 genes that are highly expressed in pRG, but not in qRG1 (Fig. 5B). The third group contains regulators for the JAK-STAT pathway (*socs3b* and *stat1*), protein folding (*hsp70l*), and innate immune response (*cxcl18b*), suggesting that these regulatory pathways are involved in the transition to pRG. We also confirmed the mRNA expression profile of these three groups of genes along the pseudo-time lineage toward qRG1 (branch1) and pRG (branch2) (Fig. 5-figure supplement 2).

Next, we focused on downregulated genes and selected TOP12 downregulated genes, whose mRNA expression was significantly downregulated in the injured condition. We performed GO enrichment analysis on these downregulated genes (Fig. 5-figure supplement 3A). Interestingly, 5 genes (*msmo1, sc5d, hmgcs1, cyp51*, and *insig1*) are enzymes involved in cholesterol synthesis (Dave and Peeples, 2021) (Fig. 5-figure supplement 3BC), and one gene (*lipg*) is involved in lipid metabolism (Yu et al., 2018) (Fig. 5-figure supplement 3D), suggesting that cholesterol synthesis is suppressed in qRG/NeuroEp1 in the injured condition. In addition, neurogenic inhibitor *id2b* is also suppressed in the injured condition (Fig. 5-figure supplement 3E). We also examined mRNA expression levels of these downregulated genes in each of 4 qRG clusters and pRG between injured and non-injured conditions (Fig. 5C). These 12 downregulated genes form one group, whose mRNA expression is very low in both qRG1 and pRG (Fig. 5C). We also confirmed mRNA expression profiles of these 12 genes along the pseudo-time lineage toward qRG1 (branch1) and pRG (branch2) (Fig. 5-figure supplement 4).

In summary, upregulation of ribosomal metabolism regulators, *ppiaa* and *vimentin* and downregulation of cholesterol biogenesis genes and *id2b* are likely associated with the transition from qRG/NeruoEp1 to both qRG1 and pRG (Fig. 5D). Upregulation of *ift45* and *b2m* may be associated with transition to qRG1, whereas upregulation of JAK-STAT pathway regulators, *cxcl18b*, and *hsp70l* may be associated with the transition to pRG (Fig. 5D).

### Microglia are activated at 1 dpl in response to TBI

In zebrafish, activation of an inflammatory response is required for a regenerative response of RGs after TBI (Kyritsis et al., 2012). First, we examined whether microglia are activated by TBI in zebrafish telencephalon. Microglia show ramified morphologies in a resting state, whereas they become rod-shape or adopt an amoeboid-like morphology in a reactive state. Therefore, we examined microglial morphology by measuring the cell sphericity index, which is defined as the ratio of the surface area of a sphere of the same volume as the given object to the surface area of the object (Fernandez-Arjona et al., 2017). We scanned microglia in adult telencephalon using the transgenic line *Tg[mpeg1.1:EGFP]*, which visualizes microglia (Ranawat and Masai, 2021), and calculated their sphericity indices. Indeed, microglia show a range of sphericity from 0.2 to 0.7 along the axis of cell shape complexity from high to low (Fig. 6-figure supplement 1A). Next, we examined the spatiotemporal profile of the sphericity of microglia in the telencephalon where TBI was induced in one hemisphere. First, we examined the sphericity of microglia located in the Dd area adjacent to the injured site and compared their sphericity distribution with that of microglia located in the equivalent Dd area of the opposite non-injured hemisphere (Fig. 6-figure supplement 1B). The sphericity index for microglia in lesioned areas was shifted to a higher range than that of microglia in control areas at 1 dpl, but not at 3 dpl (Fig. 6-figure supplement 1C), suggesting that microglia are activated at 1 dpl. To examine this more precisely, we focused on microglia with high sphericity, which were defined as more than 0.6 (Fig. 6-figure supplement 1D). The fraction of microglia located in the lesioned hemispheres in both lesioned and non-injured control hemispheres was significantly higher for high microglia sphericity at 1 dpl than in non-injured control conditions, but not at 3 dpl (Fig. 6-figure supplement 1E). Biased distribution of total microglia in the lesioned hemispheres was not observed at both 1 dpl and 3 dpl (Fig. 6-figure supplement 1E). These observations suggest that microglia are activated in response to TBI at 1 dpl.

**Figure 6.**
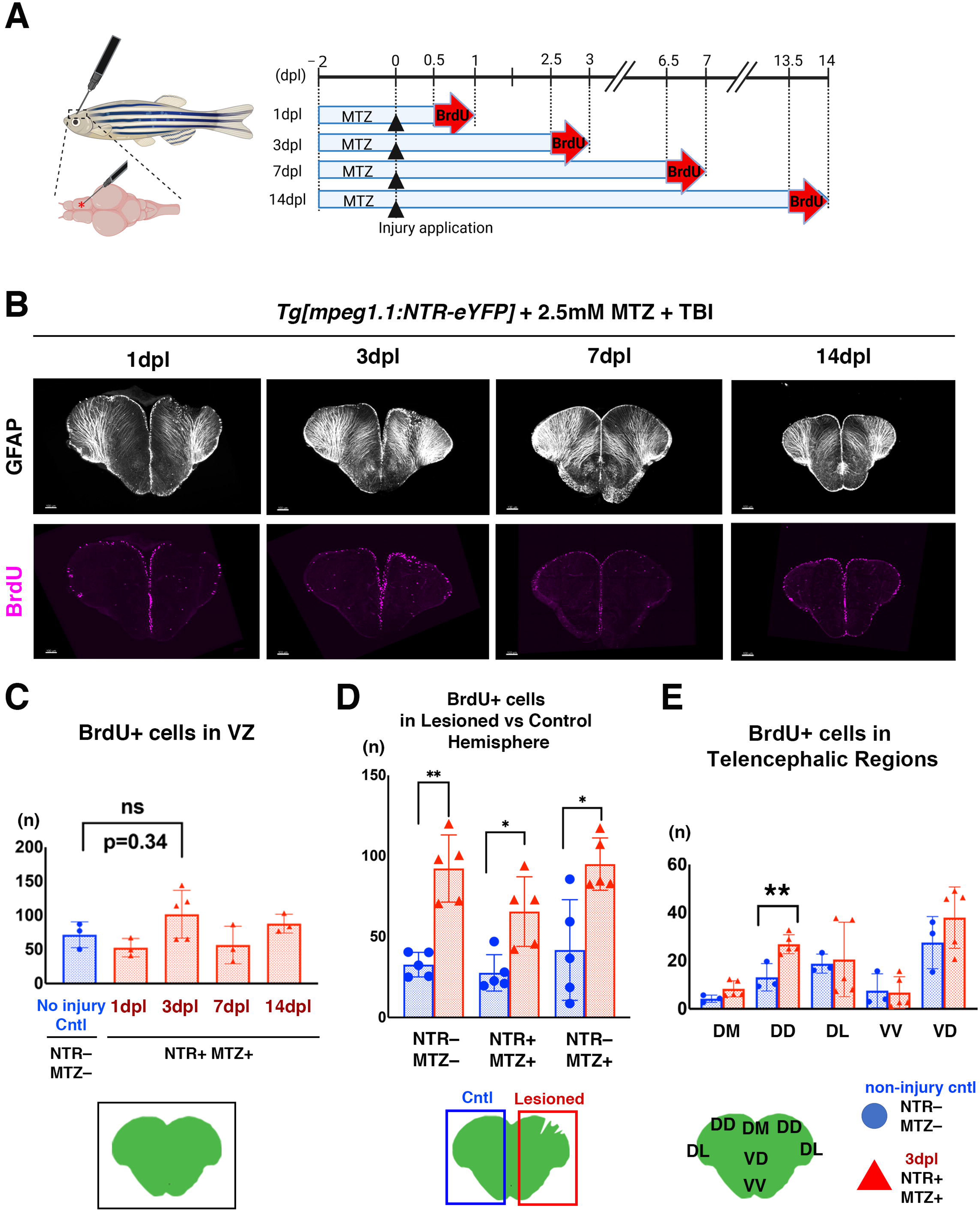
Microglial depletion compromises regenerative responses of RGs after TBI. (A) Experimental design to investigate the temporal profile of regenerative responses of RGs to TBI. Stab injury was introduced through the skull into the right hemisphere of the telencephalon. MTZ was applied to fish for 2 days prior to TBI. After TBI, BrdU was applied for 12 h prior to analysis at 1, 3, 7, and 14 dpl. Image was created using Biorender (https://biorender.com). (B) Double labeling of microglia-depleted telencephalon with anti-GFAP (white) and anti-BrdU (magenta) antibodies at 1, 3, 7, and 14 dpl. Scales: 70μm. (C) The number of BrdU+ cells in the ventricular zones of both hemispheres. The non-injured control (NTR–; MTZ–) is the same sample used for analysis in Fig. 1F. There was no significant difference between non-injured control and 3 dpl brains. mean±SD. One-way ANOVA with Dunnett’s multiple comparisons test. (D) Comparison of the number of BrdU^+^ cells between lesioned and intact hemispheres at 3 dpl in three different conditions: (1) NTR–; MTZ–, (2) NTR+; MTZ+, and (3) NTR–; MTZ+. Microglia-intact control (NTR–; MTZ–) is the same sample used for analysis in Fig. 1G. As in microglia-intact conditions (NTR–; MTZ– and NTR–; MTZ+), the number was significantly higher in lesioned hemispheres than in non-injured control hemispheres for NTR+; MTZ+. However, the BrdU+ cell number was decreased in microglia-depleted conditions (NTR+; MTZ+), compared with microglia-intact conditions (NTR–; MTZ– and NTR–; MTZ+). mean±SD. p*<0.05, p**<0.01; paired t-test. (E) The number of BrdU^+^ cells in different ventricular domains of telencephalon: Dm, Dd, Dl, Vv and Vd domains. Non-injured control (NTR–; MTZ–) was the same sample used for analysis in Fig. 1H. The number was significantly higher in the Dd domain in 3-dpl microglia-depleted brain (NTR+; MTZ+) than in non-injured control microglia-intact brain (NTR–; MTZ–). P**<0.01; unpaired students’ t-test. **Source data 1**. Data for Figure 6CDE **Source data 2.** Data for Figure 6-figure supplement 1B **Source data 3.** Data for Figure 6-figure supplement 1E **Source data 4.** Data for Figure 6-figure supplement 2BC **Source data 5.** Data for Figure 6-figure supplement 3B **Figure 6-figure supplement 1.** Microglia are activated in lesioned hemispheres at 1dpl **Figure 6-figure supplement 2.** NTR-MTZ treatment affects microglial integrity and reduces their number. **Figure 6-figure supplement 3**. Microglial depletion reduces the proliferative response of RGs.

### Genetic elimination of microglia with the NTR-MTZ system

Next, to determine the role of microglia in TBI-mediated neuronal regeneration, we eliminated microglia in adult zebrafish telencephalon. We used *Tg[mpeg1.1:NTR-eYFP]*, which expresses nitroreductase (NTR) under the control of the *mpeg1.1* promoter. NTR-mediated conversion of metronidazole (MTZ) results in cytotoxic reagent formation, which causes death of NTR-expressing cells (Petrie et al., 2014). We applied 2.5 mM MTZ within 2 or 3 days to *Tg[mpeg1.1:NTR-eYFP]* transgenic zebrafish at 4 – 7 mpf and examined whether microglia are eliminated (Fig. 6-figure supplement 2A). The number of microglia was decreased in telencephalon of *Tg[mpeg1.1:NTR-eYFP]* transgenic zebrafish treated with MTZ for 2 days compared with those with no MTZ treatment, although the difference was not significant (p=0.0752) (Fig. 6-figure supplement 2B). On the other hand, the number of microglia was significantly decreased in telencephalon of *Tg[mpeg1.1:NTR-eYFP]* transgenic zebrafish treated with MTZ for 3 days, compared with those with no MTZ treatment (Fig. 6-figure supplement 2B). We also examined microglial sphericity in telencephalon of MTZ-treated *Tg[mpeg1.1:NTR-eYFP]* transgenic zebrafish. Microglial sphericity was significantly increased in *Tg[mpeg1.1:NTR-eYFP]* transgenic zebrafish treated with MTZ for 2 days compared with those with no MTZ treatment (Fig. 6-figure supplement 2C), but not in *Tg[mpeg1.1:NTR-eYFP]* transgenic zebrafish treated with MTZ for 3 days (p=0.0592) (Fig. 6-figure supplement 2C). Although the number of microglia was not significantly decreased 2 days after MTZ treatment, microglia are likely to become unhealthy, suggesting that elimination of microglia starts 2 days after MTZ treatment. Since treatment with MTZ at 2.5 mM for 3 days or at 5 mM for 2 days affected telencephalon morphology (data not shown), we decided to apply MTZ at 2.5 mM within 2 days for all MTZ-related experiments.

### Microglia elimination suppresses the TBI-induced proliferative response of RGs

To examine the role of microglia in the TBI-induced proliferative response of RGs, we applied MTZ at 2.5 mM for 2 days to *Tg[mpeg1.1:NTR-eYFP]* transgenic zebrafish at 4 – 7 mpf, and then introduced TBI. We applied BrdU for 12 h prior to dissection of their telencephalons, which were fixed with PFA at 1, 3, 7, and 14 dpl (Fig. 6A). Fixed telencephalons were sliced at 100 μm with a vibratome. Slices were labeled with anti-GFAP and anti-BrdU antibodies (Fig. 6B). First, we evaluated TBI-induced RG proliferation in MTZ-treated transgenic telencephalons. Previously, our observations indicated that the number of BrdU^+^ RGs increased significantly at 3 dpl telencephalons compared with non-injured controls (Fig. 1E and 1F). However, in MTZ-treated injured telencephalons of *Tg[mpeg1.1:NTR-eYFP]* transgenic fish, the number of BrdU^+^ RGs increased slightly at 3 dpl, but not significantly compared to non-injured controls of MTZ-untreated, non-transgenic telencephalons (Fig. 6C). Next, we compared the number of BrdU^+^ RGs between lesioned and non-injured hemispheres at 3 dpl. Previously, our observations indicated that the number of BrdU^+^ RGs is significantly higher in lesioned hemispheres than in non-injured hemispheres in telencephalons (Fig. 1G). On the other hand, the number of BrdU^+^ RGs in lesioned hemispheres of MTZ-treated *Tg[mpeg1.1:NTR-eYFP]* transgenic telencephalons was decreased compared with that of lesioned hemispheres in MTZ-untreated, non-transgenic telencephalons or in MTZ-treated, non-transgenic telencephalons (Fig. 6D); however, the number was still significantly higher than in a non-injured hemispheres of MTZ-treated *Tg[mpeg1.1:NTR-eYFP]* transgenic telencephalons (Fig. 6D). Previously, our observations indicated that RGs in the Dd area more efficiently responded to TBI and incorporated BrdU (Fig. 1H). A similar significant increase in the number of BrdU+RGs was observed in the Dd area of MTZ-treated *Tg[mpeg1.1:NTR-eYFP]* transgenic telencephalons compared with non-injured controls of MTZ-untreated, non-transgenic telencephalons (Fig. 6E). However, the number of BrdU+ RGs in the Dd area was significantly decreased in MTZ-treated *Tg[mpeg1.1:NTR-eYFP]* transgenic telencephalons, compared with that of MTZ-treated, non*-*transgenic telencephalons (Fig. 6-figure supplement 3). These observations suggest that microglial elimination affects the proliferative response of RGs in response to TBI, although a local response was still retained to some degree.

### The fractions of qRG1 and pRG decrease in response to TBI in microglia-depleted telencephalons

We applied scRNA-seq analysis to zebrafish MTZ-treated *Tg[mpeg1.1:NTR-eYFP; gfap:dTomato]* transgenic telencephalons with TBI treatment. Similar to scRNA-seq analysis of TBI-induced and non-injured control telencephalons (Fig. 2A), stab injuries were introduced in telencephalons of 4 mpf adult zebrafish. At 3 dpl, telencephalons were homogenized to prepare cell suspensions, which were subjected to flow-cytometry-assisted cell sorting to collect the *gfap:dTomato*^+^ RG fraction (Fig. 2-figure supplement 1BC). We used a 10x genomics system to prepare scRNA-seq libraries. After read sequencing and mapping to the zebrafish genomic database GRCz11, we evaluated nUMI, nGene, and mitochondrial genes for two datasets, non-injured and 3-dpl MTZ-treated *Tg[mpeg1.1:NTR-eYFP]* brains, to ensure that all our data were comparable (Fig. 2-figure supplement 1E). We analyzed 992 cells from 3-dpl MTZ-treated *Tg[mpeg1.1:NTR-eYFP]* brains following the thresholding for unique gene counts and mitochondrial gene expression and obtained scRNA-seq data on RGs of 3-dpl MTZ-treated *Tg[mpeg1.1:NTR-eYFP]* brains, which we called “3dpl+MTZ brains” or “3dpl microglia-depleted brain” hereafter (supplementary table 4). We conducted unbiased clustering of the dataset from 3-dpl+MTZ sample, using the Seurat package (Butler et al., 2018). In 3-dpl+MTZ sample, cells were classified into 11 clusters (Fig. 7-figure supplement 1A), which correspond to 4 types of qRGs, pRG/Neuroblast/NBN, 3 types of MN/NBN, ependymal cells, pineal gland, microglia (MG)/macrophage (Fig. 7-figure supplement 1B, and Supplementary Table 4), confirming sufficient quality to apply further clustering analysis of merged datasets. Then, we used the Harmony merging algorithm to integrate two datasets: scRNA-seq data of RGs between 3-dpl and 3dpl+MTZ brains, the former of which is shown in Fig. 2. After clustering analysis of merged scRNA-seq data, we identified 11 clusters, including 6 qRG populations, pRGs/Neuroblasts/NBNs, MNs/NBNs, ependymal cells, pineal gland cells, microglia/macrophages (Fig. 7A, and Supplementary Table 5). We determined the TOP10 differentially expressed genes for each cluster (Fig. 7-figure supplement 2) and found that 6 qRG clusters contain qRG/NeuroEp1, qRG/NeuroEp2, qRG1 and RG2, which are identified by RNA-seq analysis on merged data of 3-dpl and non-injured controls (Fig. 2), and two new RG clusters, namely, qRG3 and qRG4 (Fig. 7-figure supplement 2). Interestingly, in their TOP10 genes, qRG3 and qRG4 express *cxcl18b* and *id2b*, respectively, which are upregulated and downregulated in qRG/NeuroEp1 under 3-dpl conditions (Fig. 5C, Fig. 5-figure supplement 1A, and Fig. 5-figure supplement 3A). To clarify the origin of qRG3 and qRG4, we mapped 3-dpl populations of qRG3 and qRG4 to UMAP of 3-dpl scRNA-seq data (Fig. 7-figure supplement 3AB). We found that qRG3 corresponds to a part of qRG/NeuroEP1 in 3-dpl UMAP, whereas qRG4 corresponds to a part of qRG/NeuroEP1 and OPC in 3-dpl UMAP (Fig. 7-figure supplement 3C). Thus, four qRG populations, qRG/NeuroEP1, qRG/NeuroEP2, qRG1 and qRG2, remain to be clustered in merged data of 3-dpl and 3dpl+MTZ brain, although small fractions of qRG/NeuroEP1 form two new clusters, qRG3 and qRG4. We also confirmed how each of the 11 clusters of the merged datasets corresponded to 11 clusters of 3 dpl+MTZ cells (Fig. 7-figure supplement 1D).

**Figure 7.**
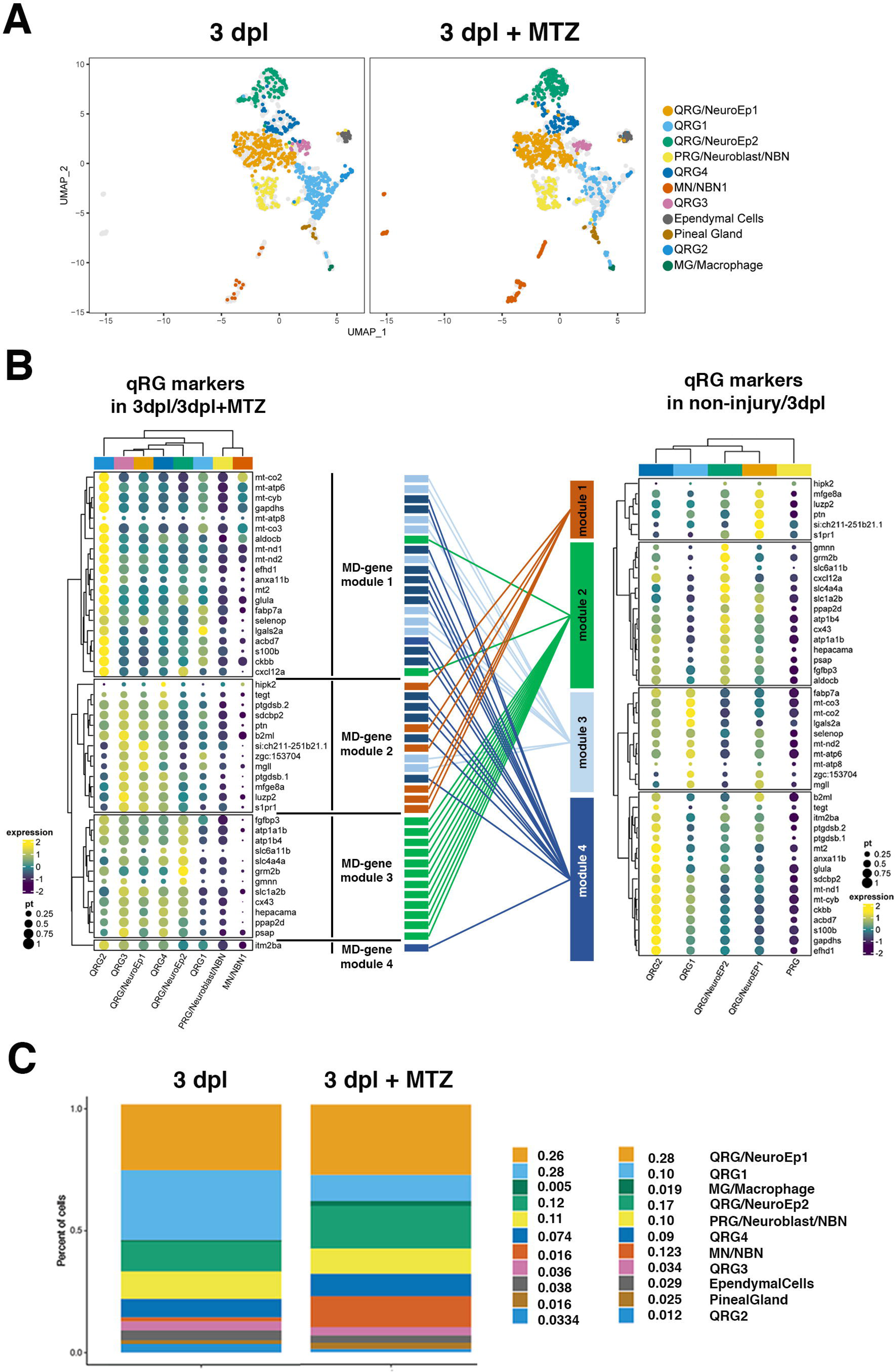
scRNA-seq analysis reveals that qRG1 is decreased in 3-dpl microglia-depleted telencephalon. (A) UMAP of scRNA-seq data in the 3-dpl and 3 dpl+MTZ merged dataset. Twelve clusters are identified: 6 qRG, pRG/Neuroblast/NBN, Ependymal cells, Pineal gland, MN/NBN, Microglia (MG)/Macrophage. (B) Dot-plot analysis of qRG marker expression in 6 qRG clusters, pRG/Neuroblast/NBN, and MN/NBN of the 3-dpl and 3 dpl+MTZ merged dataset. qRG marker genes are classified into 4 new gene modules, called DM-gene modules 1-4. qRG marker genes of “Gene module 3” in the non-injured control and 3-dpl merged dataset are rearranged into two MD-gene modules 1 and 2. Furthermore, qRG marker genes of “Gene module 1” in the non-injured control and 3-dpl merged dataset are also merged with “Gene module 4” to form MD-gene module 2. (C) Fractions of each cluster in the 3-dpl and 3 dpl+MTZ merged dataset. qRG1 is decreased in the 3-dpl+MTZ condition, whereas qRG/NeuroEp2 and MN/NBN are increased. **Figure 7-figure supplement 1.** Cell clusters in the 3 dpl+MTZ dataset. **Figure 7-figure supplement 2.** Heatmap of the top 10 markers for clusters identified in the 3-dpl and 3 dpl+MTZ merged dataset. **Figure 7-figure supplement 3.** Clarifying origins of new qRG3 and qRG4 clusters in the 3-dpl and 3dpl+MTZ merged dataset. **Figure 7-figure supplement 4.** Dot-plot analysis of pRG marker expression in clusters of the 3-dpl and 3 dpl+MTZ merged dataset.

Next, we examined 46 qRG markers in 6 qRG clusters in merged data of 3-dpl and 3dpl+MTZ brain (Fig. 7B). 46 qRG markers were split into four gene modules, “Microglia-depleted (MD)-gene modules1–4”. MD-gene module 1 contains 20 genes that are highly expressed in qRG2. MD-gene module 1 contains 10 qRG markers of “Gene module 4” and 8 qRG markers of “Gene module 3” in non-injured controls and the 3-dpl merged dataset (Fig. 7B). “Gene module 3” genes are highly expressed in qRG1; however, “Gene module 3”-derived genes except *lgals2a* of MD-gene module 1 seem to be downregulated in qRG1. DM-gene module 2 contains 13 genes that are highly expressed in qRG/NeuroEp1, qRG3, and qRG4. MD-gene module 2 contains all 6 “Gene module 1” genes in non-injured controls and the 3-dpl merged dataset (Fig. 7B). Since qRG3 and qRG4 are thought to be derived from qRG/NeuroEp1, it is reasonable that DM-gene module 2 genes are highly expressed in qRG3 and qRG4. DM-gene module 3 contains 12 genes that are highly expressed in qRG/NeuroEp1, qRG/NeruoEp2, qRG3 and qRG4. All 12 genes of DM-gene module 3 are “Gene module 2” genes in non-injured controls and the 3-dpl merged dataset (Fig. 7B), suggesting that DM-gene module 3 is equivalent to “Gene module 2”. DM-gene module 4 contains only one gene, *itm2ba*, which is highly expressed in all 6 qRG populations. *itm2ba* belongs to “Gene module 4” of non-injured controls and the 3-dpl merged dataset (Fig. 7). Thus, “Gene module 3” genes, which are highly expressed in qRG1, are split into MD-gene modules 1–2, whereas “Gene module 1” genes, which are highly expressed in qRG/NeuroEp1, are merged with “Gene module 4” genes to form DM-gene module 2. Thus, microglia elimination rearranges the state of qRG/NeuroEp1 and qRG1.

Next, we examined pRG markers. pRG markers were split into 4 gene modules, DM-modules 1–4 (Fig. 7-figure supplement 4). DM-gene module 1 contains 11 genes that are highly expressed in pRG/Neuroblast/NMN and MN/NBN, but show low expression in all 6 qRG clusters. DM-gene module 2 contains 6 genes that are highly expressed in pRG/Neuroblast/NMN and MN/NBN, as well as qRG1 and qRG2. DM-gene module 3 contains 16 genes that are highly expressed only in pRG/Neuroblast/NMN. Most genes in DM-gene modules 1–3 overlapped with pRG markers of “Gene module 1” in non-injured controls and the 3-dpl merged dataset (Fig. 7-figure supplement 4). DM-gene module 4 contains 6 genes that are highly expressed in pRG/Neuroblast/NMN as well as qRG/NeuroEp1, qRG/NeuroEp2, qRG3 and qRG4. Five of these 6 genes overlap with “Gene module 3” genes in merged data of non-injured and 3-dpl brains (Fig. 7-figure supplement 4). Thus, “Gene module 2” genes, which are highly expressed in qRG1, are split into DM-gene modules 1–3.

Next, we compared the fraction of each cluster between 3-dpl and 3dpl+MTZ conditions (Fig. 7C). Interestingly, the fraction of qRG1 was markedly decreased in 3dpl+MTZ conditions (10.0%), compared with 3-dpl (28.0%). Thus, qRG1 increases in response to TBI in a microglia-dependent manner. In addition, the fraction of qRG/NeuroEP2 and MN/NBN clusters were increased in 3dpl+MTZ conditions, compared with 3-dpl.

To identify what kind of signaling pathway is a potential target of microglia during the TBI-mediated transition from qRG/NeuroEp1 into either qRG1 or pRG, we determined upregulated and downregulated genes in qRG/NeuroEp1 of 3dpl+MTZ brain relative to 3-dpl brain (Fig. 8A). GO analysis revealed that genes related to ribosomal metabolism (*rps* and *rpl* genes), chaperone cofactor-mediated protein folding (*hsp70l, hsp70.1, hsp70.4, hsp90aa1.2*), response to cytokine and JAK-STAT pathway (*cxcl12a, stat2, socs3a*), Jun (*jun, junba, junbb, jund*), Hes (*her4.3, her4.4, her6*) are downregulated in 3-dpl+MTZ brain, compared with 3-dpl brain (Fig. 8B, Fig. 8-figure supplement 1A). Downregulation of mRNA expression in qRG/NeuroEp1 was confirmed on UMAP of 3-dpl and 3dpl+MTZ for the JAK-STAT pathway (Fig. 8-figure supplement 1BC), for chaperone cofactor-mediated protein folding (Fig. 8-figure supplement 1D), for ribosomal metabolism (Fig. 8-figure supplement 1E), for Hes (Fig. 8-figure supplement 1F), and for Jun (Fig. 8-figure supplement 1G). Furthermore, GO analysis revealed that four genes related to wound response and sensory epithelial regeneration, *hbegfa, mdka, hmgb1a*, and *lgals2a* are upregulated in qRG/NeuroEp1 (Fig. 8C, Fig. 8-figure supplement 2A). Upregulation of mRNA expression in qRG/NeuroEp1 was confirmed in UMAP of 3-dpl and 3 dpl+MTZ (Fig. 8-figure supplement 2BC) and dot-plot histogram (Fig. 8B).

**Figure 8.**
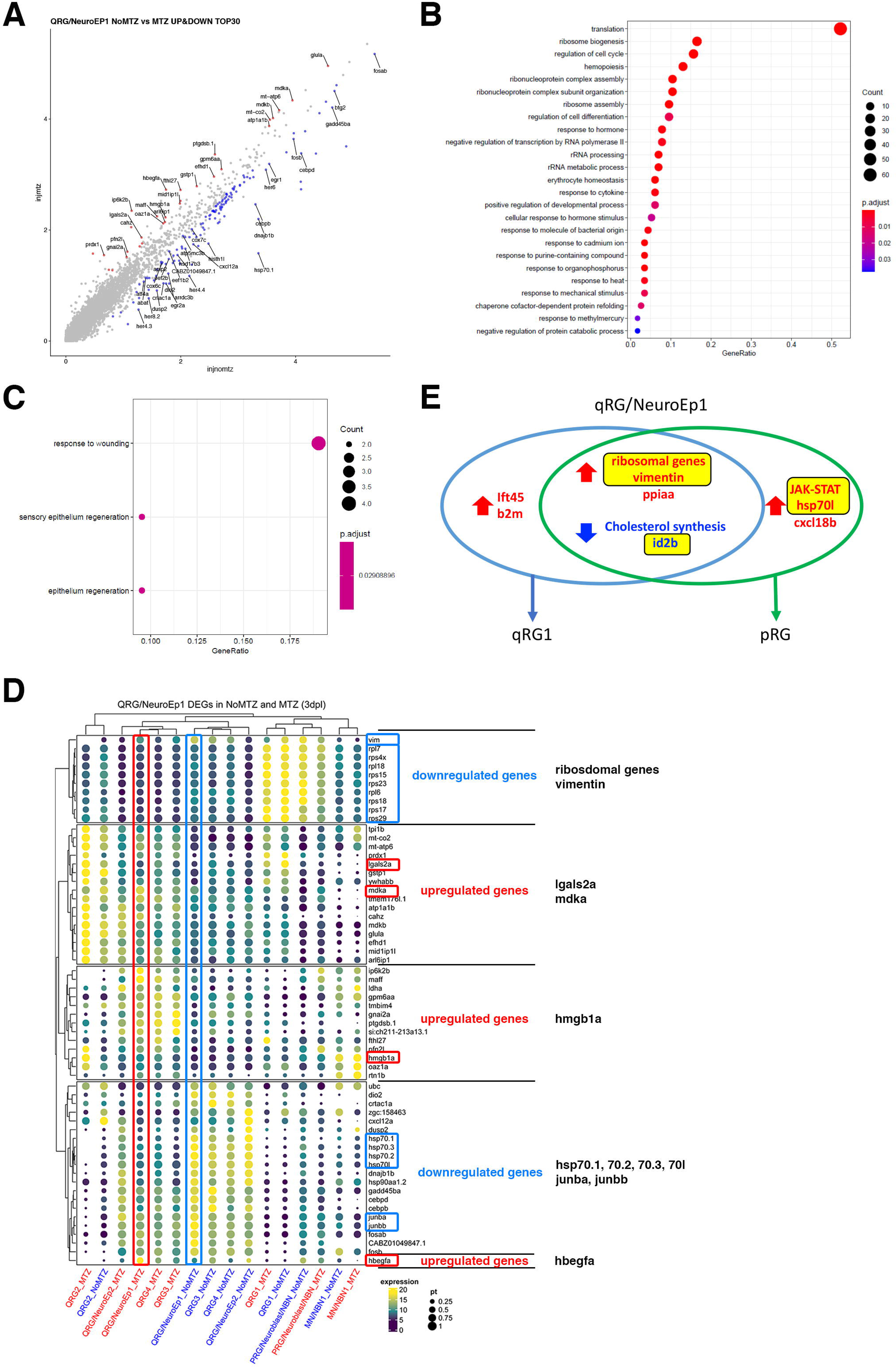
DEG analysis of qRG/NeuroEp1 in response to TBI. (A) Scatter plot visualization of differentially expressed genes (DEGs) in qRG/NeuroEp1 at 3 dpl under microglia-depleted conditions versus microglia-intact conditions. Upregulated (red) and downregulated (blue) genes are mapped. (B) GO analysis of downregulated genes in qRG/NeurEp1 in response to microglial depletion. (C) GO analysis of upregulated genes in qRG/NeurEp1 in response to microglial depletion. (D) Dot-plot visualization of DEGs of qRG/NeuroEp1 across all RG clusters at 3 dpl under microglia-depleted and -intact conditions. Gene expression columns of qRG/NeuroEp1 under the 3-dpl and 3 dpl+MTZ conditions are indicated by blue and red squares, respectively. The TOP30 upregulated genes and TOP30 downregulated genes are shown. Genes nominated by GO analysis are shown in the right-most column: 4 upregulated genes: *lgasl2a*, *mdka*, *hmgb1a*, and *hbegfa*, and downregulated genes: ribosomal genes, *vimentin*, *hsp70.1*, *hsp 70.2*, *hsp70.3*, *hsp 70l*, *junba*, and *junbb*. (E) Venn diagram of up- and downregulated genes in qRG/NeuroEp1 linked to the transition to qRG1 and pRG/Neuroblast/NBN. Potential targets of microglia are indicated by yellow squares. **Figure 8-figure supplement 1.** qRG/NeuroEp1 downregulated genes in response to microglial depletion. **Figure 8-figure supplement 2.** qRG/NeuroEp1 upregulated genes in response to microglial depletion.

We also confirm downregulation of these genes in dot-plot histogram (Fig. 8D). These data suggest that microglia normally promote mRNA upregulation of genes related to ribosomal genesis, *vimentin*, the JAK-STAT pathway, and *hsp70l* in response to TBI (Fig. 8E). Upregulation of *cxcl18b, ppiaa, ift45,* and *b2m* mRNA and downregulation of genes related to cholesterol biogenesis are induced by TBI, independent of or upstream of microglia-mediated expansion of qRG1 and pRG. This may be consistent with the observation that a new EP1-derived cluster, qRG3, expresses *cxcl18b* (Fig. 7-figure supplement 2). In addition, a new EP1-derived cluster, qRG4, expresses *id2b* (Fig. 7-figure supplement 2), it is likely that microglia elimination affects *id2b* downregulation in qRG/NeuroEp1 or its derived qRG4 in response to TBI (Fig. 8E). It is possible that microglia act on these clusters to change their states into qRG1 and pRG during the TBI-mediated regenerative response of qRG.

## Discussion

In this study, we analyzed scRNA of RGs in adult zebrafish telencephalon in response to TBI. In the merged data set of non-injured control and 3-dpl conditions, we identified 4 qRG clusters, qRG/NeuroEp1, qRG/NeuroEp2, qRG1 and qRG2. Previous scRNA analysis of RGs in zebrafish adult telencephalon revealed 47 qRG-specific genes (Lange et al., 2020). Interestingly, our 4 identified qRG clusters express different combinations of these qRG-specific genes, indicating heterogeneity of qRGs in zebrafish. Our regional marker analysis revealed that these qRG clusters are segregated into two groups, which are localized in dorsal and ventral telencephalic domains. *vimentin* and *zic1* are highly expressed in qRG of dorsal (pallium) and ventral (subpallium) domains of telencephalon, respectively. qRG1 and qRG/NeuroEp1 are likely to be pallial qRGs, because of *zic1*-negative and *vimentin*-positive clusters. On the other hand, qRG2 and qRG/NeuroEp2, seem to be subpallial qRGs, because of *zic1*-positive and *vimentin*-negative clusters. Thus, it is likely that qRG1 and qRG/NeuroEp1 are spatially segregated to qRG2 and qRG/NeuroEp2.

Among 4 qRG clusters, qRG/NeuroEp1 seems to be a dormant qRG, because qRG/NeuroEp1 prominently expresses *Notch3* and *mfge8*, which are required for stemness and quiescence, respectively (Than-Trong et al., 2018; Zhou et al., 2018). qRG1 is a ribosomal gene-enriched cluster and markedly increased in cell number in response to TBI. Consistently, qRGs expressing a mTOR marker, p-rpS6, are significantly increased in dorsal pallium of injured hemispheres, compared with non-injured control hemispheres. Pseudo-time analysis revealed that qRG/NeuroEp1 produces qRG1 and pRG via independent lineages. This is in contrast to primed quiescent NSCs in the V/SVZ of adult mouse brain, which shows a similarly ribosomal gene-enriched population and represents the intermediate state from dormant NSCs to active proliferative NSCs (Dulken et al., 2017; Llorens-Bobadilla et al., 2015).

What is the physiological role of qRG1 in the regenerative response of RGs after TBI? One of differentially expressed genes in qRG1 is *lgals2*, which encodes Galectin-2. Galectins are β-galactose-binding animal lectins (Nio-Kobayashi, 2017). Galectin-1 is expressed in V/SVZ astrocytes of adult mouse brain and promotes proliferation of NSCs in a carbohydrate-binding manner (Sakaguchi et al., 2006). Galectin-1 also promotes proliferation of neural progenitor cells in the SGZ of the dentate gyrus of adult mouse brain (Kajitani et al., 2009). Galectin-1 and 3 are required for astrocyte reactivity after adult mouse brain injury (Sirko et al., 2015). In animal models of experimental autoimmune encephalomyelitis (EAE), Galectin-1 is secreted from astrocytes, binds to M1 microglia, and negatively regulates M1 microglia-mediated neurotoxicity, leading to promotion of M2 microglia-mediated neuroprotection (Starossom et al., 2012). Thus, Galectins regulate astrocyte or NSC behaviors through modulation of the immune system. In zebrafish retina, Galectin-2 expression is rapidly induced in retinal NSCs called Müller glia after light-induced damage, and its knockdown affected the rod regenerative response (Craig et al., 2010), suggesting that upregulation of *lgals2* expression is the first regenerative response of retinal NSCs in zebrafish. Indeed, our data showed that *lgals2*-expressing cells are induced only in the ventricular region near the injury site of the brain, and that the number of *lgals2*-expressing cells correlates with the size of injury (Fig. 3C). A possible scenario is that damaged qRG/NeuroEp1 cells exit the dormant state and become qRG1, which subsequently expresses Galectin-2. Galectin-2 secreted from qRG1 may act on microglia to migrate into the damaged area for clearance of dead and damaged cells. Galectin-2 may promote surrounding qRG/NeuroEp1 cells to differentiate into pRGs in concert with microglia. If this is the case, qRG1 may function as a sensor to detect damage levels and promote the transition from dormant qRGs to pRGs. It will be interesting to examine whether the TBI-induced regenerative response of qRGs is affected in adult zebrafish *lgals2* knockdown telencephalon.

Another important question is what kinds of regulatory factors are involved in production of qRG1 and pRG by qRG/NeuroEp1. We examined up- and down-regulated genes in qRG/NeuroEp1 at 3 dpl versus non-injured controls. GO analysis revealed that four signaling pathways, ribosomal metabolism, JAK/STAT pathway, cytokine signaling including *cxcl18b*, and chaperonin-mediated protein folding, including *hsp70l* and *ppiaa*, are upregulated in 3-dpl qRG/NeuroEp1. In addition, vimentin, *ift45* and *b2m* are upregulated. Interestingly, these upregulated genes are classified into three groups. The first group (ribosomal genes, *vimentin, ppiaa*) is highly expressed in both pRGs and qRG1 after injury. The second group (*ift45, b2m*) is highly expressed in qRG1, but not pRG, and the third group (JAK-STAT, *cxcl18l, hsp70l*) is highly expressed in pRG, but not qRG1 (Fig. 5C). It is possible that these three group genes differentially generate both pRG and qRG1, qRG1 or pRG, respectively (Fig. 5D). Furthermore, three signaling pathways: cholesterol biogenesis, a lipid metabolism factor lipg, and a neurogenic inhibitor, *id2b*, are downregulated in 3-dpl qRG/NeuroEp1. Almost all these downregulated genes show low expression in both qRG1 and pRG at 3 dpl, suggesting that their downregulation promotes the transition from qRG/NeuroEp1 to both qRG1 and pRG (Fig. 5D). At present, it is unknown how ribosomal metabolic genes are upregulated in qRG/NeuroEp1 by TBI. It was reported that *mfge8* is highly expressed in quiescent NSCs in the SGZ of adult mouse dentate gyrus, and that Mfge8 promotes NSC quiescence by suppressing the mTOR pathway in an autocrine manner (Zhou et al., 2018). Since mfge8 is highly expressed in qRG/NeuroEp1, but downregulated along the lineage toward qRG1 and pRG (Fig. 4B), downregulation of *mfge8* may activate the AKT/mTOR pathway to increase ribosomal genesis in both qRG1 and pRG.

Another interesting finding is the downregulation of cholesterol synthesis genes. ER membrane-localized cholesterol sensor protein (*insig1*) and four components of the cholesterol synthesis pathway (*hmgcs1, cyp51, msmo1, sc5d*) are downregulated in qRG/NeuroEp1 after injury. Furthermore, endothelial lipase (*lipg*), which promotes lipid biosynthesis through cleavage of high-density-lipoprotein (HDL) (Yu et al., 2018), is downregulated in qRG/NeuroEp1 after injury. Brain cholesterol metabolism is tightly regulated through cooperation between neurons and different glial-cell types (Dai et al., 2021; Dave and Peeples, 2021). Since peripheral cholesterol cannot enter the brain across the blood-brain-barrier, cholesterol in adult brain is synthesized *de novo* in glial cells, primarily astrocytes. Cholesterol is loaded onto apolipoprotein E (ApoE) in astrocytes and secreted through ATP binding cassette (ABC) transporters. Neurons and oligodendrocytes take up lipidated ApoE particles via receptor-mediated endocytosis, so they depend on astrocytes for cholesterol. However, in chronic demyelinated lesions, top-downregulated genes in astrocytes are involved in cholesterol synthesis (Itoh et al., 2018), suggesting that reduction of cholesterol biogenesis is an initial response of astrocytes after brain damage. At present, its physiological significance is unknown; however, it was recently reported that the stem cell marker Prominin1 promotes axon regeneration by downregulating cholesterol synthesis via Smad signaling (Lee et al., 2020). Thus, cholesterol biogenesis was implicated in axonal regeneration. It will be interesting to investigate how cholesterol regulates regenerative responses of RGs after brain injury.

Previous studies revealed that acute inflammation is required for the regenerative response of RGs to TBI in zebrafish (Kyritsis et al., 2012). We confirmed that microglial depletion compromises the regenerative response of RGs to TBI. Next, scRNA-seq analysis was applied to 3-dpl zebrafish telencephalon under microglia-depleted conditions. In the merged dataset of GFAP:dTomato^+^ cells in 3-dpl zebrafish telencephalon with and without microglia, 11 clusters were identified. Among them, qRGs are classified into 6 clusters. qRG/NeuroEp1, qRG/NeuroEp2, qRG1, qRG2, qRG3 and qRG4. qRG3 and qRG4 are a part of qRG/NeuroEp1 in 3-dpl conditions, suggesting that these clusters are derived from qRG/NeuroEp1. Interestingly, the fraction of qRG1 is markedly decreased in 3-dpl microglia-depleted conditions, compared with 3-dpl microglia-intact conditions, suggesting that expansion of qRG1 by brain injury depends on microglia. To determine what kinds of signaling networks are targeted by microglia during brain damage, upregulated and downregulated genes in qRG/NeuroEp1 were compared between 3-dpl microglia-depleted conditions and microglia-intact conditions. GO analysis revealed that downregulated genes pertain to ribosomal metabolism, chaperone-mediated protein folding, cytokine and JAK-STAT signaling, jun, and her4/6. These data suggest that upregulation of ribosomal metabolism, chaperone-mediated protein folding, cytokine and JAK-STAT signaling in qRG/NeuroEp1 after TBI depend on microglia. Only four genes in qRG/NeuroEp1 are upregulated: *lgals2a, mdka, hmgb1a,* and *hbegfa*. Although the qRG1 fraction is markedly decreased, the qRG1-specific marker *lgals2a* is highly expressed in qRG/NeuroEp1. So, TBI may induce *lgals2a* transcription even in microglia-depleted conditions; however, *lgals2a*-expressing cells fail to differentiate into qRG1, because ribosomal genes and vimentin fail to be upregulated in qRG/NeuroEp1 after TBI. *cxcl18b* expression in qRG/NeuroEp1 was not downregulated in 3-dpl microglia-depleted conditions and *cxcl18b* is highly expressed in qRG3, which is likely to derive from qRG/NeuroEp1. These observations suggest that *cxcl18b* upregulation of 3-dpl qRG/NeuroEp1 occurs prior to microglial action. Similarly, expression of cholesterol biogenesis genes was not upregulated in qRG/NeuroEp1 under microglia-depleted conditions, suggesting that downregulation of cholesterol biogenesis gene expression occurs prior to microglial action. Taken together, these data illustrate that modulation of gene expression for cholesterol biogenesis, *ppiaa, cxcl18b, id2b, ift45*, and *b2m* occurs prior to the interaction between microglia and qRG/NeuroEp1, whereas expression of genes for ribosomal metabolism, vimentin, the JAK/STAT pathway, *hsp70l*, *junba/bb*, and *her4/6* are modulated by the interaction between microglia and qRG/NeuroEp1 (Fig. 8E). In the future, further studies are necessary to identify the mechanism underlying the interaction between microglia and dormant RGs during TBI.

## Materials and Methods

### Fish Strains

Zebrafish (*Danio rerio*) were maintained using standard procedures (Westerfield, 2000). Adult zebrafish at 3-7 months old were used for all experiments. Female and male fish were randomly selected for analysis. *okinawa wild-type* (*oki*) was used for the wild-type strain. *Tg[GFAP:GFP]*^mi2001^ (Bernardos and Raymond, 2006) and *Tg[GFAP:dTomato]*^nns17Tg^ (Satou et al., 2012) were used to visualize qRG. *Tg[mpeg1.1:EGFP]*^oki053^ (Ranawat and Masai, 2021) was used to visualize microglia. *Tg[mpeg1:NTR-EYFP]*^w202^ (Petrie et al., 2014) was used for depletion of microglia.

### Ethics statement

All zebrafish experiments were performed in accordance with the Animal Care and Use Program of Okinawa Institute of Science and Technology Graduate University (OIST), Japan, which is based on the Guide for the Care and Use of Laboratory Animals by the National Research Council of the National Academies. The OIST animal care facility has been accredited by the Association for Assessment and Accreditation of Laboratory Animal Care (AAALAC International). All experimental protocols were approved by the OIST Institutional Animal Care and Use Committee. The OIST Institutional Animal Care and Use Committee approved all experimental protocols (Protocol no. 2019-267, 2019-268, 2019-269, 2019-270).

### Stab lesion assay

Following anesthesia of adult zebrafish with Tricaine/MS222 (200μg/ml), a needle with a length of 2 mm and an outer diameter of 300 μm was vertically inserted through the skull into the medial region of the right hemisphere of the telencephalon. After lesioning, fish were maintained in separate tanks until the times for analysis and were sacrificed with ice water following Tricaine-mediated anesthesia. Heads were separated from the bodies at the level of the gills, and fixed with 4% PFA for 2 h at room temperature or 6 h at 4°C. Afterward, brains were dissected from skulls and fixed again with 4% PFA overnight at 4°C. After injuring the fish, when mild hemorrhage was noticed, they were permitted to recover in fresh water, which was daily changed. Uninjured intact zebrafish were used for controls.

### Histology and antibody labeling

Using a vibratome (Dosaka, EM Co. Ltd., Linear Slicer PRO7), 100-μm frontal sections were prepared along the antero-posterior axis of the adult telencephalon. These vibratome sections were used for immunolabeling according to a published protocol (Schmidt et al., 2014). The following primary antibodies were used at an appropriate dilution rate: anti-BrdU (Abcam, ab6326, 1:400), anti-HuC/D (ThermoFisher, A21271, 1:500), rabbit anti-GFP (ThermoFisher, A11122, 1:200), anti-GFAP (zrf1) (ZIRC, 1:200), anti-phosphorylated rpS6 (Cell signaling, 2211,1:500). Anti-mouse, anti-rabbit, and anti-rat antibodies conjugated with Alexafluor dyes (Alexa 546, and Alexa 633) were used as secondary antibodies (ThermoFisher, 1:500). Immunolabeled sections were mounted on glass slides in PBS buffer with 70% glycerol for confocal LSM scanning. Vibratome brain slices were scanned with A1R confocal LSM (Nikon). 3D images covering a large area were captured using a 20x/0.75 objective lens with tiling and stitching functions. Sample preparation and immunolabeling of cryo-sections were carried out according to standard protocols (Masai et al., 2003). Cryo-sectioned samples were scanned with confocal LSM, A1R (Nikon).

### *In situ* hybridization

cDNA fragments of *gstm.1* and *lgals2a* were amplified by RT-PCR using mixed cDNA synthesized from the heads of 1-4 dpf embryos as a template and the following primers:

*gstm.1* forward, 5’-ATGGCAATGAAATTGGCTTATTG-3’

*gstm.1* reverse, 5’-TCACTCCTTCTTGTTTCCCCA-3’

*lgals2a* forward, 5’-ATGGCCGGTGTGCTTATACAGA-3’

*lgals2a* reverse, 5’-CTATTTAATTTCAACCCCTTGGAT-3’

Amplified fragments were subcloned into the PCRII-TOPO vector (Invitrogen). Antisense and sense RNA probes were synthesized from the linearized plasmid template by *in vitro* transcription using a DIG RNA Labeling Kit (Roche). *In situ* hybridization was performed on cryo-sections of telencephalon as previously described (Masai et al., 2003) with a minor modification. Hybridization was performed overnight at 60°C using *lgals2a*, *gstm.1* antisense RNA probes at 0.52 ng/μL and 0.31 ng/μL, respectively.

### BrdU incorporation, detection, and analysis

Adult fish were incubated in 10 mM BrdU-containing water for 12 h and sacrificed to dissect telencephalons and prepare vibratome- or cryo-sections for further analysis. To increase efficiency of anti-BrdU antibody labeling, vibratome- and cryo-sections were pretreated with 2N HCl for 1 h at room temperature to expose BrdU antigen from nuclear packed chromatin. BrdU-incorporated brain slices were labeled with anti-BrdU and anti-GFAP (zrf1) antibodies and scanned with confocal LSM A1R (Nikon) with large-scan combined tiling and stitching functions. GFAP and BrdU double-positive nuclei in the ventricular zone of the dorsal pallium were visualized by surface rendering using Imaris software (Bitplane) and used for statistical analysis using GraphPad Prism (ver.9).

### Sphericity analysis of microglia

Vibratome brain slices of zebrafish transgenic line *Tg[mpeg1.1:EGFP]* were scanned with confocal LSM A1R (Nikon). After confocal scanning, target telencephalic areas were analyzed using Imaris software (Bitplane) to prepare surface rendering objects for each of *mpeg1.1:EGFP*+ microglia. For sphericity analysis, the sphericity index was defined as the ratio of the surface area of a sphere of the same volume as a given object to the surface area of the object (Fernandez-Arjona et al., 2017). The sphericity index was calculated using surface area and volume data for each object. Statistical analysis was performed using GraphPad Prism (ver.9).

### Surface-Rendering Process

To identify proliferating cells localized in the ventricular zone, the niche of RG, GFAP^+^ surface was created, followed by creating BrdU^+^ surfaces. Depending on the threshold used for creating GFAP and BrdU surfaces, a colocalization channel was created. The colocalized surface, which is identical to the BrdU^+^ surface, but is not located in the parenchyma, was counted as proliferating cells in the ventricular zone. If necessary, threshold adjustment was carried out to decrease the noise level during surface-creation. To evaluate the sphericity of microglia, due to their complicated shapes and their close association with the lesioned site, a manual threshold was specified across all sections and samples for surface rendering.

### Microglia depletion by NTR-MTZ system

To eliminate microglia from adult zebrafish brains, we used a transgenic line *Tg[mpeg1:NTR-eYFP]* that expresses NTR under the control of the *mpeg1.1* promoter (Petrie et al., 2014). Primer information for genotyping the transgene *Tg[mpeg1:NTR-eYFP]* are provided in Key Resource Table. Fish were maintained in water containing 2.5 mM MTZ from 2 days prior to injury until harvesting of brains. To evaluate levels of microglia elimination, the transgenic line *Tg[mpeg1.1:NTR-eYFP]* was combined with another transgenic line *Tg[mpeg1.1:EGFP]* (Ranawat and Masai, 2021), which labels microglia or microglial precursor cells more efficiently. This double transgenic line was used for sphericity analysis (Fig. 6-figure supplement 1) and for counting microglial numbers (Fig. 6-figure supplement 2).

### Fraction of p-rpS6-positive area in the GFAP-positive Dd ventricular zone

Cryosections of zebrafish adult telencephalons carrying the transgene *Tg[GFAP:EGFP]* were labeled with anti-p-rpS6 antibody and scanned with confocal LSM. In the scanning images, the ROI corresponding to the Dd ventricular domain of injured hemispheres or non-injured control hemispheres was selected. Using Image-J (ver.1.44, NIH), GFAP:EGFP^+^ areas were outlined by changing the threshold adjustment to remove background noise, and then the fraction of the GFAP:EGFP^+^ area in the ROI was determined. Similarly, the fraction of the p-rpS6^+^ area in the same ROI was determined. Using the area of the ROI, we calculated the area of p-rpS6^+^ and GFAP:EGFP^+^ regions. The fraction of p-rpS6^+^ in the GFAP:EGFP^+^ area in the ROI of each brain section was calculated. Statistical significance was determined using paired t-tests with GraphPad Prism (ver. 9).

### Fraction of *lgals2a-* and *gstm.1-positive* cells in the total hemisphere area

Cryosections of zebrafish adult telencephalon were labeled with *lgasl2a* or *gstm.1* RNA probe. RNA signals were outlined using Image-J (ver.9) by changing the threshold adjustment to remove background noise. The fraction of the mRNA-expressing area in the total area of the injured or non-injured hemisphere was calculated. Statistical significance was determined using one-way ANOVA with Tukey’s multiple comparison test, using GraphPad Prism (ver. 9).

### Fluorescence-Activated Cell Sorting of RGs

Telencephalons were dissected in ice-cold Hank’s Balanced Salt Solution (HBSS) (0.137 M NaCl, 5.4 mM KCl, 0.25 mM Na_2_HPO_4_, 0.1% glucose, 0.44 mM KH_2_PO_4_, 1.3 mM CaCl_2_, 1.0 mM MgSO_4_, 4.2 mM NaHCO_3_) from non-lesioned or 3-dpl *Tg[gfap:dTomato]* fish. For the microglia-ablated system, MTZ-treated *Tg[gfap:dTomato; mpeg1.1:NTR-eYFP]* fish were used. Seven telencephalons per experimental condition were pooled and processed. Isolated telencephalons were chemically dissociated for 30 - 40 min at room temperature in HBSS containing papain (20 U/mL) and DNase (2000U/mL) with gentle trituration every 10 min. Cell suspensions were filtered with a 40-μm cell strainer (Falcon). Then 10% FBS in L-15 medium was added to stop the enzymatic reaction. Dissociated cells were washed with 10% FBS/PBS solution, and the final pellet was resuspended in 10% FBS/PBS for cell sorting. Before cell sorting, cells were again passed through a 40-μm cell strainer, and Sytox blue dead cell stain (Invitrogen) was added to label dead cells. FACS sorting was performed using a BD FACSAria II, and 9,000 – 20,000 dTomato^+^ cells were collected in a 10% FBS/PBS precoated tube. Collected cells were concentrated by centrifugation at 300 x *g* for 5 min and resuspended in 50-60 μL of 10%FBS/PBS.

### scRNA-seq library preparation

Collected dTomato^+^ cells were loaded onto a 10X Chromium System using 10X v3 chemistry. Single-cell RNAseq libraries were prepared with 12 cycles of cDNA amplification and 15 cycles of indexing PCR, and were sequenced with NovaSeq6000 (Illumia).

### Computational analysis

Sequencing data were analyzed using 10x Genomics Cell Ranger 3.1.0 (Zheng et al., 2017) to perform alignment, filtering, barcode counting, and UMI counting with default settings and the zebrafish reference genome (GRCz11). Filtered output matrices were used for clustering and downstream analysis in the Seurat v4.0 R package (Hao et al., 2021), where cells with a unique feature count between 200 and 2500 and a fraction of mitochondrial transcripts less than 5% and genes expressed in at least 3 cells were used for analysis. Each dataset was normalized using the SCTransform function with regressing out of mitochondrial gene expression as a possible source of variation. Control and 3-dpl datasets were merged using the Harmony algorithm (Korsunsky et al., 2019), followed by principal component analysis. Top 20 principal components were used for cluster generation with resolution = 0.5. UMAP was used to visualize resulting clusters in low-dimensional space. To identify each cluster’s differentially expressed genes, the Findmarkers function with the Wilcoxon rank sum test was used. Differentially expressing genes (DEGs) between controls and 3-dpl datasets in a cluster were determined using the Findmarker function with DESeq2 test. To identify upregulated and downregulated biological processes between control and 3-dpl datasets, gene ontology (GO) enrichment analysis was performed with the list of significantly upregulated and downregulated genes (adjusted *p* value < 0.05) using the clusterProfiler R package (Yu et al., 2012). Trajectory inference was performed using the Monocle3 R package (Qiu et al., 2017). OPC, Ependymal Cells, Pineal Gland, and MN/NBN clusters were dropped from control and 3-dpl merged objects were analyzed with Seurat. Then the object was split into control and 3-dpl. After individual Seurat objects were converted to monocle3 objects with the as.cell_data_set function, preprocessing using num_dim=10, dimensionality reduction, and UMAP projection was performed. To learn the trajectory graph and to order the cells in pseudotime, learn_graph and order_cells functions were used with default parameters.

### Statistical Analysis

In this study, we used brain samples as a biological replicate and brain slices as a technical replicate. As for sample size determination, a minimum of 3 biological replicates were used in all the experiments. When experiments were designed, no power analysis was utilized. Statistical analysis was conducted using GraphPad Prism 9.4.0. For analyzing proliferative responses in 3-dpl injury conditions (Fig. 1) and 3-dpl+MTZ conditions (Fig. 6), the same non-injured control sample was used for the comparison. In the case of analyzing hemisphere-dependent responses, only sections with clear evidence of lesion sites were included in calculations from each biological replicate. The number of samples used per experiment is represented in dot-pot histogram and Source Data files. Tests used to calculate statistical significance are mentioned in figure legends. All data are presented as means ± SDs. A p-value of 0.05 was considered statistically significant. No data point was removed based on variability. In histograms, p*< 0.05, p**< 0.01, p***< 0.001, p****< 0.0001, or ns (not significant) is indicated.

### Data availability

Raw scRNA-seq datasets and filtered matrices generated by Cell Ranger 3.1.0 have been deposited in the NCBI’s Gene Expression Omnibus (GEO) and accessible through GEO Series accession number GSE209705.

## Acknowledgements

We thank Dr. Yuko Nishiwaki, Dr. Tetsuya Harakuni, and Mr. Yutaka Kojima for supporting experiments. We thank Dr. Randall T. Moon and Dr. Jeanot Muster for providing the *Tg[mpeg1.1:NTR-eYFP]* line. We thank the OIST Research Support Division: the Sequencing Section (SQC) for assistance in sequencing, the Instrumental Analysis Section (IAS) for assistance and support in FACS sorting, and the Scientific Imaging Section (SIG) for assistance and support in imaging experiments.

## Legends of Supplementary Figures

**Figure 2-figure supplement 1. FACS cell sorting and scRNA-seq data quality check.**

(A) Setting of gate parameters for sorting dTomato^+^ cells for non-injured control and 3-dpl samples in microglia-intact conditions (NTR–/MTZ–).

(B) Setting of gate parameters for sorting dTomato^+^ cells for non-injured control and 3-dpl samples in microglia-depleted conditions (NTR+/MTZ+).

(C) Percentage of target cell population for four experimental conditions after FACS sorting.

(D, E) scRNA data quality check for non-injured and 3-dpl samples in microglia-intact conditions (NTR–/MTZ–) (D) and microglia-depleted conditions (NTR+/MTZ+) (E). VLN plot for the number of genes detected per cell (nFeature_RNA), the number of UMIs per cell (nCount_RNA) and mitochondrial gene percentages per cell (percent.mt).

**Figure2-figure supplement 2. Cell clusters in the non-injured control dataset.**

(A) UMAP of the non-injured control dataset. Six clusters are specified.

(B) Heatmap of the top 10 differentially expressed genes for each of 6 clusters.

(C) Mapping of 6 clusters of non-injured control samples on UMAP of the merged dataset for non-injured controls and 3-dpl samples. Clusters 0–5 correspond to qRG/NeuroEp1, qRG/NeuroEp2, qRG1+pRG/Neuroblast/NBN+MN/NBN, Ependymal cells+OPC, qRG2, Pineal gland, respectively.

**Figure2-figure supplement 3. Cell clusters in the 3-dpl dataset.**

(A) UMAP of the 3-dpl control dataset. Seven clusters are specified.

(B) Heatmap of the top 10 differentially expressed genes for each of 7 clusters.

(C) Mapping of 7 clusters of 3-dpl samples on UMAP of the merged dataset for non-injured controls and 3-dpl samples. Clusters 0–6 correspond to qRG/NeuroEp1, qRG1+qRG2, qRG/NeuroEp1+OPC, qRG/NeuroEp2, qRG1, pRG/Neuroblast/NBN+MN/NBN, Ependymal cells, respectively.

**Figure2-figure supplement 4. Heatmap of the top 10 markers for clusters in the non-injured control and 3-dpl merged dataset.**

The non-injured control and 3-dpl merged dataset showed 9 clusters. The Top 10 differentially expressed genes are shown in each cluster. In accordance with differentially expressed genes, we classified 9 clusters to 4 qRG (qRG/NeuroEp1, qRG/NeuroEp2, qRG1, qRG2), pRG/Neuroblast/NMN, MN/NBN, OPC, Ependymal cells, Pineal gland.

**Figure2-figure supplement 5. Gene module analysis of qRG markers.**

(A) Dot-plot of qRG marker expression of 4 qRG and pRG. 46 qRG markers were classified into 4 gene modules: Gene modules 1-4 are highly expressed in qRG/NeuroEp1, qRG/NeuroEp2, qRG1 and qRG2, respectively.

(B) UMAP of the non-injured control and 3-dpl merged dataset.

(C) Feature plot analysis of Gene Module1 highly expressed in the qRG/NeuroEp1 cluster.

(D) Feature plot analysis of Gene Module2 highly expressed in the qRG/NeuroEp2 cluster.

(E) Feature plot analysis of Gene Module3 highly expressed in the qRG1 cluster.

(F) Feature plot analysis of Gene Module3 highly expressed in the qRG2 cluster.

**Figure2-figure supplement 6. Regional marker analysis of qRGs.**

(A) Dot-plot analysis of regional markers. *zic1* (Vv) was specifically expressed in qRG/NeuroEp2 and qRG2.

(B) UMAP of the non-injured control and 3-dpl merged dataset.

(C) Feature plot analysis of regional marker genes. *zic1* (Vv) was specifically expressed in qRG/NeuroEp2 and qRG2.

**Figure2-figure supplement 7. Expression analysis of genes responsible for quiescence and stemness of qRGs.**

(A) Dot-plot analysis of genes related to quiescence and stemness.

(B) UMAP of the non-injured control and 3-dpl merged dataset.

(C) Feature plot analysis of genes related to quiescence and stemness.

**Figure2-figure supplement 8. Gene module analysis of pRG markers.**

(A) Dot-plot analysis of pRG markers. pRG markers were classified into 3 gene modules.

(B) UMAP of the non-injured control and 3-dpl merged dataset.

(C) Feature plot analysis of Gene Module 1.

(D) Feature plot analysis of Gene Module 2, whose expression is high in qRG1.

(E) Feature plot analysis of Gene Module 3, whose expression is high in qRG/NeuroEp1/2.

**Figure 4-figure supplement 1. Pseudo-time analysis using Monocle3 for non-injured control and 3-dpl conditions.**

(A) Upper: UMAP with cell trajectory generated using Monocle 3 for non-injured control (left) and 3-dpl (right) conditions. Lower: pseudo-time transition of cell trajectory.

(B) Gene expression analysis along pseudo-time trajectory.

**Figure 4-figure supplement 2. Gene expression analysis of branch1- and branch2-associated pRG/Neuroblast/NBN cells.**

(A) Left: Cell trajectory generated with Monocle 3 in 3-dpl conditions. pRG/Neuroblast/NBN cells are indicated in light blue. Right: UMAP of extracted pRG/Neuroblast/NBN cells associated with Branch 1 and Branch 2. Branch 1-associated qRG/Neuroblast/NBN were separated from Branch 2-associated qRG/Neuroblast/NBN.

(B) Dot-plot of differentially expressed genes (DEGs) in Branch1- and Branch2-associated pRG/Neuroblast/NBN cells. Branch 1-associated pRG/Neuroblast/NBN cells show a very similar expression profile to qRG1.

(C) Violin plot of DEGs in Branch 1- and Branch 2-associated pRG/Neuroblast/NBN cells. qRG1-specific RG markers are shown in red squares, and highly expressed Branch 1-associated pRG/Neuroblast/NBN cells.

**Figure 4-figure supplement 3. AhR signaling pathway is specifically activated in qRG/NeuorEp2.**

(A) UMAP of the non-injured control and 3-dpl merged dataset.

(B) Feature plot analysis of AhR signaling pathway genes, arh2 and cyp1b1.

(C) Violin plot of AhR signaling pathway genes, arh2 and cyp1b1.

(D) Dot-plot of expression of AhR target genes in qRGs and pRG.

**Figure 5-figure supplement 1. Signaling network of upregulated genes in qRG/NeuroEp1 in response to TBI.**

(A) Classification of upregulated genes in regulatory pathways; ribosomal metabolism, cytokine signaling and the JAK/STAT pathway.

(B) UMAP of the non-injured control and 3-dpl merged dataset.

(C) Feature plot analysis of upregulated genes related to JAK-STAT signaling.

(D) Feature plot analysis of upregulated genes related to cytokine signaling.

(E) Feature plot analysis of upregulated genes related to ribosomal metabolism.

**Figure 5-figure supplement 2. Gene expression profiles of the top 30 upregulated genes in qRG/neuroEp1 along a pseudo-time trajectory.**

Top: The top 30 upregulated genes whose expression is high in both qRG1 and pRG. Middle: The top 30 upregulated genes whose expression is high in qRG1.

Bottom: The top 30 upregulated genes whose expression is high in pRG.

qRG1 lineage (Branch 1) and pRG lineage (Branch 2) at 3 dpl are shown at the left and right columns, respectively.

**Figure 5-figure supplement 3. Signaling network of downregulated genes in qRG/NeuroEp1 in response to TBI.**

(A) Classification of downregulated genes in regulatory pathways. Inhibition of neurogenesis, lipid metabolism, and cholesterol synthesis.

(B) UMAP of the non-injured control and 3-dpl merged dataset.

(C) Feature plot analysis of downregulated genes related to cholesterol metabolism.

(D) Feature plot analysis of downregulated genes related to lipid metabolism.

(E) Feature plot analysis of downregulated *id2b* expression indicating inhibition of neurogenesis.

**Figure 5-figure supplement 4. Gene expression profiles of the top 12 downregulated genes in qRG/neuroEp1 along a pseudo-time trajectory.**

(A) The top 12 downregulated genes in the 3-dpl qRG1 lineage (Branch 1) using dot-plotting graph along a pseudo-time trajectory path.

(B) The top 12 downregulated genes in the 3-dpl pRG lineage (Branch 2) using a dot-plotting graph along a pseudo-time trajectory path.

**Figure 6-figure supplement 1. Microglia are activated in lesioned hemispheres at 1 dpl.**

(A) Microglial morphology with different sphericity indices. Scales: 10 μm.

(B) Visualization of mpeg1.1:EGFP^+^ microglia in non-injured control, 1 dpl, and 3 dpl telencephalon. The Dd domain in non-injured control and lesioned hemispheres are indicated by blue and red squares, respectively. In the case of non-injured control conditions, left and right hemispheres were used instead of non-injured controls and lesioned hemispheres of 1- and 3-dpl brain. Scales: 100 μm.

(C) Sphericity distribution of microglia in the Dd domain of non-injured control and lesioned hemispheres at 1 dpl and 3 dpl.

(D) Higher magnification of blue and red squares shown in panel (B). Microglia with high and low sphericity are shown in yellow and red, respectively. Scales: 20 μm.

(E) The percentage of the number of microglia located in the lesioned hemispheres relative to the number of microglia located in both lesioned and non-injured control hemispheres. Histogram for microglia with high sphericity (index > 0.6) and histogram for all microglia are shown to the left and right, respectively. Only high-sphericity microglia at 1dpl are significantly higher than in non-injured conditions. p*<0.05: One-way ANOVA, Dunnett’s multiple comparison test.

**Figure 6-figure supplement 2. NTR-MTZ treatment affects microglial integrity and reduces their number.**

(A) Visualization of mpeg1.1:EGFP^+^ microglia in *Tg[mpeg1.1:NTR-eYFP]* transgenic telencephalon with no MTZ treatment, MTZ treatment for 2 days, and MTZ treatment for 3 days. No microglial cells were detected in MTZ treatment for 3 days. Scales: 100 μm.

(B) Number of microglia per section of no MTZ, 2d MTZ, and 3d MTZ treatments. The number is significantly lower in 3d MTZ treatment than in no MTZ treatment. p***<0.005: One way ANOVA, Bonferroni multiple comparison test.

(C) Sphericity index distribution of microglia with no MTZ, 2d MTZ, and 3d MTZ treatments. The sphericity index is significantly higher in 2d MTZ treatment than in no MTZ treatment; however, there is no significant difference between 3d MTZ treatment and no MTZ treatment. p***<0.005: One way ANOVA, Bonferroni multiple comparison test.

**Figure 6-figure supplement 3. Microglial depletion reduces the proliferative response of RGs.**

(A) Double labeling of 3-dpl microglia-depleted (NTR+; MTZ+) and microglia-intact (NTR–; MTZ+) telencephalon with anti-GFAP and anti-BrdU antibodies. Lower panels indicate higher-magnification images of dotted squares shown in the upper panels. Scales: 70 μm (upper panels) and 30 μm (lower higher magnification image panels).

(B) The number of BrdU^+^; GFAP^+^ cells, which correspond to pRGs, in the Dd domain of the ventricular zone. The number is significantly decreased in 3-dpl microglia-depleted telencephalon, compared with 3-dpl microglia-intact telencephalon. p*<0.05: Unpaired t test.

**Figure7-figure supplement 1. Cell clusters in the 3 dpl+MTZ dataset.**

(A) UMAP of the 3 dpl+MTZ dataset. Eleven clusters are specified.

(B) Heatmap of the top 10 differentially expressed genes for each of 11 clusters.

(C) UMAP of the merged dataset for 3-dpl and 3 dpl+MTZ samples.

(D) Mapping of 11 clusters of 3 dpl+MTZ samples on UMAP of the merged dataset for 3-dpl and 3 dpl+MTZ samples. Clusters 0–11 correspond to qRG/NeuroEp1+qRG3, qRG1+qRG2, pRG/Neuroblast/NBN, qRG4, qRG1+qRG2, 3 types of MN/NBN, Pineal gland, Ependymal cells, and Microglia (MG)/macrophage, respectively.

**Figure 7-figure supplement 2. Heatmap of the top 10 markers for clusters identified in the 3-dpl and 3 dpl+MTZ merged dataset.**

Eleven clusters are classified into qRG/NeuroEp1, qRG/NeuroEp2, qRG1, qRG2, pRG/Neuroblast/NBN, MN/NBN, Ependymal cells, Pineal gland, Microglia (MG)/Macrophage. In addition, two new qRG clusters, qRG3 and qRG4, are specified. Genes specific to each cluster are shown in the left column.

**Figure 7-figure supplement 3. Clarifying origins of new qRG3 and qRG4 clusters in the 3-dpl and 3dpl+MTZ merged dataset.**

(A) Split UMAP of the 3-dpl and 3 dpl+MTZ merged dataset.

(B) UMAP of the non-injured control and 3-dpl merged dataset.

(C) Reference mapping cell clusters identified in the 3-dpl and 3 dpl+MTZ merged dataset on UMAP of the non-injured control and 3-dpl merged dataset. qRG3 is derived from qRG/NeuroEp1, whereas qRG4 is derived from qRG/NeuroEp1 and OPC.

**Figure 7-figure supplement 4. Dot-plot analysis of pRG marker expression in clusters of the 3-dpl and 3 dpl+MTZ merged dataset.**

Dot-plot analysis of pRG marker expression in 6 qRG clusters, pRG/Neuroblast/NBN, and MN/NBN of the 3-dpl and 3 dpl+MTZ merged dataset. pRG marker genes are classified into new 4 gene modules, called DM-gene modules 1-4. pRG markers of “Gene module 1” in the non-injured control and 3-dpl merged dataset are rearranged into three MD-gene modules 1 – 3. Most pRG markers of “Gene module 3” in the non-injured control and 3-dpl merged dataset are overlapped with MD-gene modules 4.

**Figure 8-figure supplement 1. qRG/NeuroEp1 downregulated genes in response to microglial depletion.**

(A) Classification of downregulated genes in regulatory pathways: ribosomal metabolism, chaperonin-mediated protein folding, cytokine and JAK/STAT pathway, jun, her4/her6.

(B) Split UMAP of the 3-dpl and 3dpl+MTZ merged dataset.

(C) Feature plot analysis of downregulated genes related to the JAK-STAT and cytokine pathways.

(D) Feature plot analysis of downregulated genes related to chaperone-mediated protein folding.

(E) Feature plot analysis of downregulated genes related to ribosomal metabolism.

(F) Feature plot analysis of downregulated genes related to her4/her6.

(G) Feature plot analysis of downregulated genes related to jun.

**Figure 8-figure supplement 2. qRG/NeuroEp1 upregulated genes in response to microglial depletion.**

(A) Classification of upregulated genes in the regulatory pathway: wound response and epithelium regeneration.

(B) Split UMAP of the 3-dpl and 3dpl+MTZ merged dataset.

(C) Feature plot analysis of upregulated genes related to wound response and epithelium regeneration.

## List of supplementary materials

### Source Data files

**Figure 1-Source Data 1.** Data for Figure 1FGH.

**Figure 3-Source Data 1.** Data for Figure 3B.

**Figure 3- Source Data 2.** Data for Figure 3DF.

**Figure 6- Source Data 1**. Data for Figure 6CDE

**Figure 6- Source Data 2.** Data for Figure 6-figure supplement 1B

**Figure 6- Source Data 3.** Data for Figure 6-figure supplement 1E

**Figure 6- Source Data 4.** Data for Figure 6-figure supplement 2BC

**Figure 6- Source Data 5.** Data for Figure 6-figure supplement 3B

### Supplementary figures

**Figure 2-figure supplement 1.** FACS cell sorting and scRNA-seq data quality check.

**Figure2-figure supplement 2.** Cell clusters in the non-injured control dataset.

**Figure2-figure supplement 3.** Cell clusters in the 3-dpl dataset.

**Figure2-figure supplement 4.** Heatmap of the top 10 markers for clusters in the non-injured control and 3-dpl merged datasets.

**Figure2-figure supplement 5.** Gene module analysis of qRG markers.

**Figure2-figure supplement 6.** Regional marker analysis of qRGs.

**Figure2-figure supplement 7.** Expression analysis of genes responsible for quiescence and stemness of qRGs.

**Figure2-figure supplement 8.** Gene module analysis of pRG markers.

**Figure 4-figure supplement 1.** Pseudo-time analysis using Monocle3 for non-injured control and 3-dpl conditions.

**Figure 4-figure supplement 2.** Gene expression analysis of branch1- and branch2-associated pRG/Neuroblasts/NBN cells.

**Figure 4-figure supplement 3.** AhR signaling pathway is specifically activated in qRG/NeuorEp2.

**Figure 5-figure supplement 1.** Signaling network of upregulated genes in qRG/NeuroEp1 in response to TBI.

**Figure 5-figure supplement 2.** Gene expression profiles of the TOP30 upregulated genes in qRG/neuroEp1 along a pseudo-time trajectory.

**Figure 5-figure supplement 3.** Signaling network of downregulated genes in qRG/NeuroEp1 in response to TBI.

**Figure 5-figure supplement 4.** Gene expression profiles of the TOP12 downregulated genes in qRG/neuroEp1 along a pseudo-time trajectory.

**Figure 6-figure supplement 1.** Microglia are activated in lesioned hemispheres at 1dpl

**Figure 6-figure supplement 2.** NTR-MTZ treatment affects microglial integrity and reduces their number.

**Figure 6-figure supplement 3**. Microglial depletion reduces the proliferative response of RGs.

**Figure 7-figure supplement 1.** Cell clusters in the 3 dpl+MTZ dataset.

**Figure 7-figure supplement 2.** Heatmap of the top 10 markers for clusters identified in the 3-dpl and 3 dpl+MTZ merged dataset.

**Figure 7-figure supplement 3.** Clarifying origins of new qRG3 and qRG4 clusters in the 3-dpl and 3dpl+MTZ merged dataset.

**Figure 7-figure supplement 4.** Dot-plot analysis of pRG marker expression in clusters of the 3-dpl and 3 dpl+MTZ merged dataset.

**Figure 8-figure supplement 1.** qRG/NeuroEp1 downregulated genes in response to microglial depletion.

**Figure 8-figure supplement 2.** qRG/NeuroEp1 upregulated genes in response to microglial depletion.

### Supplementary Tables

**Supplementary Table 1.** Differentially expressed genes for each cluster in the non-injured dataset.

**Supplementary Table 2.** Differentially expressed genes for each cluster in the 3-dpl dataset.

**Supplementary Table 3.** Differentially expressed genes for each cluster in the non-injured control and 3-dpl merged dataset.

**Supplementary Table 4.** Differentially expressed genes for each cluster in the 3dpf+MTZ dataset.

**Supplementary Table 5.** Differentially expressed genes for each cluster in the 3-dpl and 3dpl+MTZ merged data set.

## Appendix

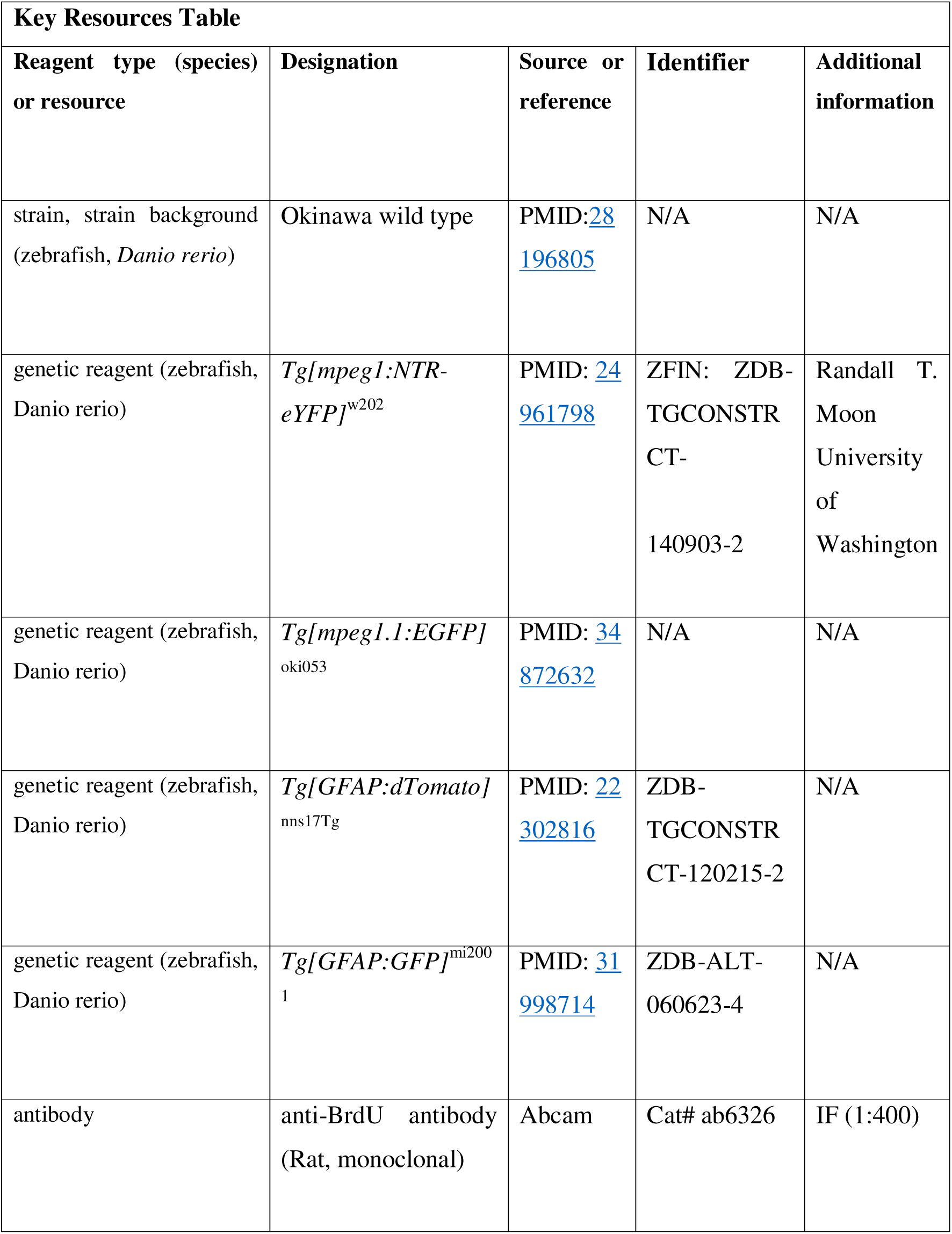
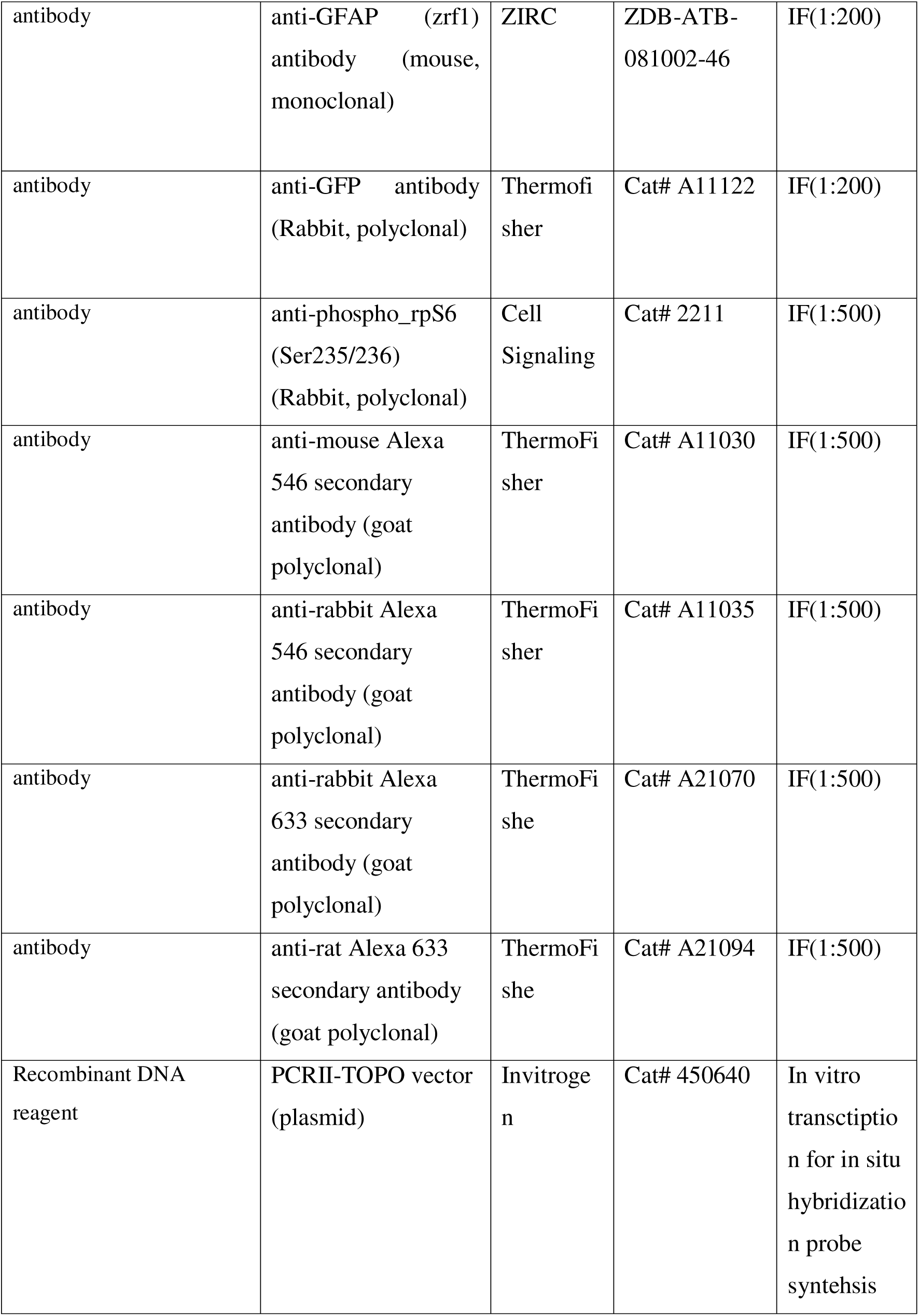
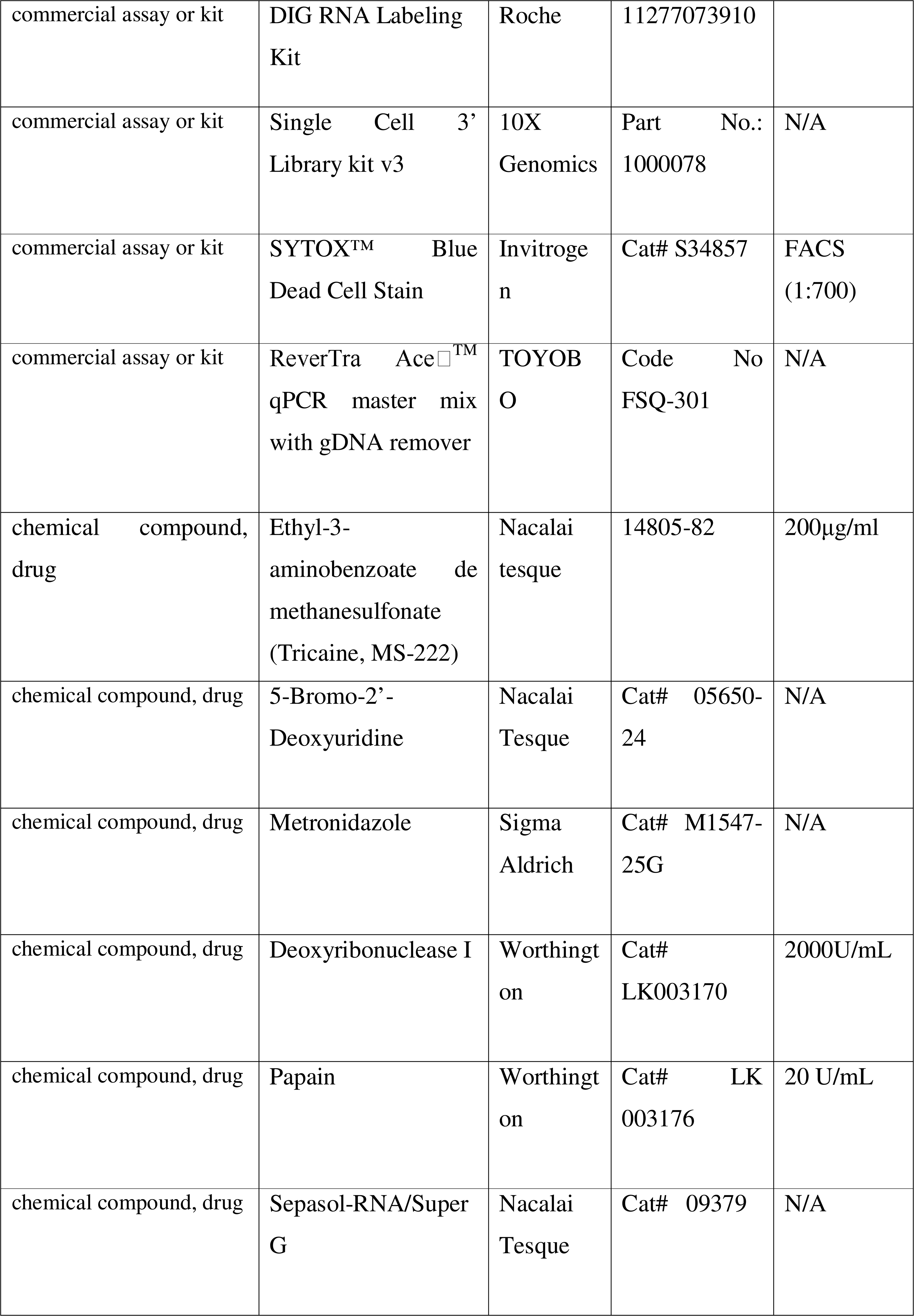
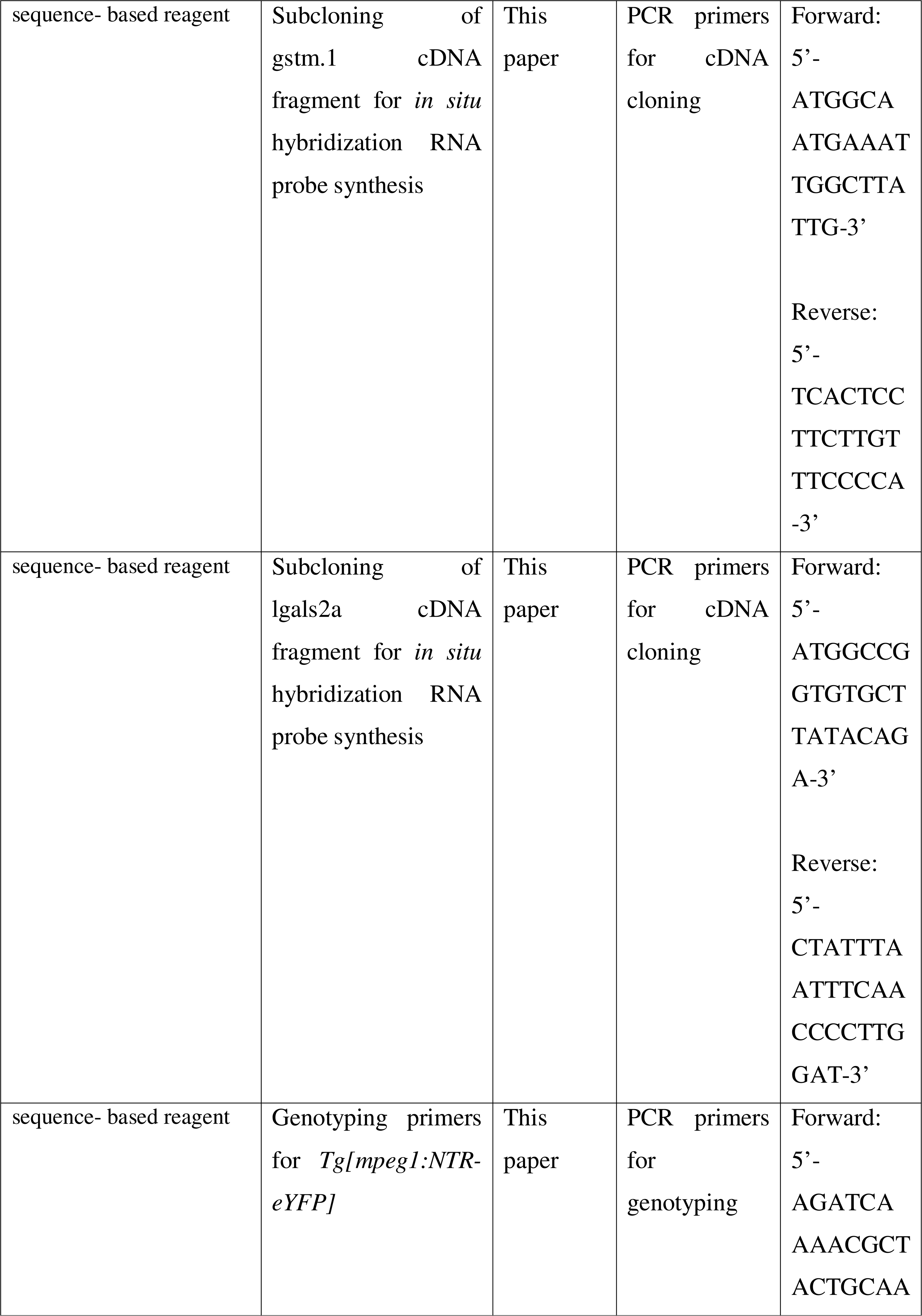
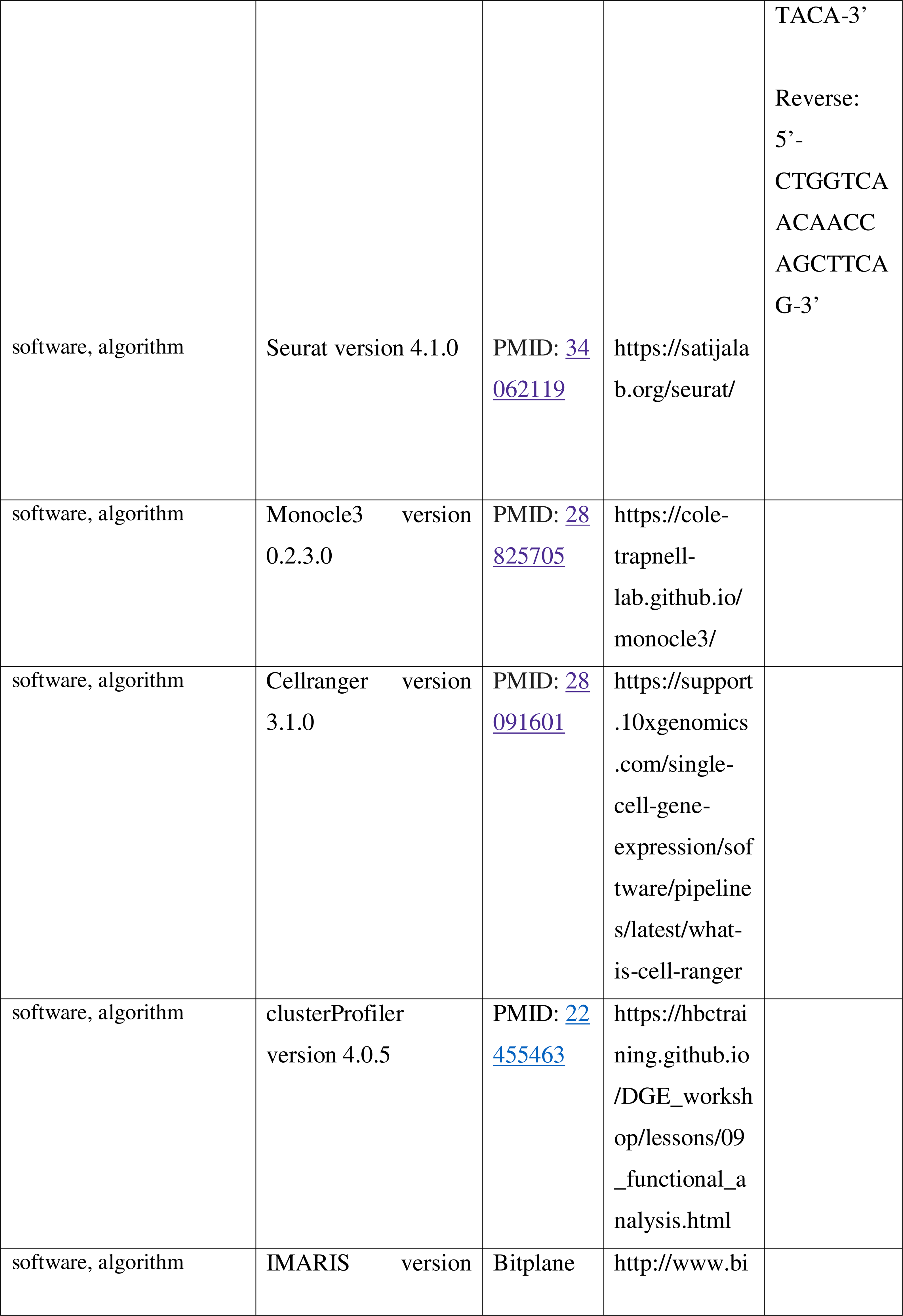
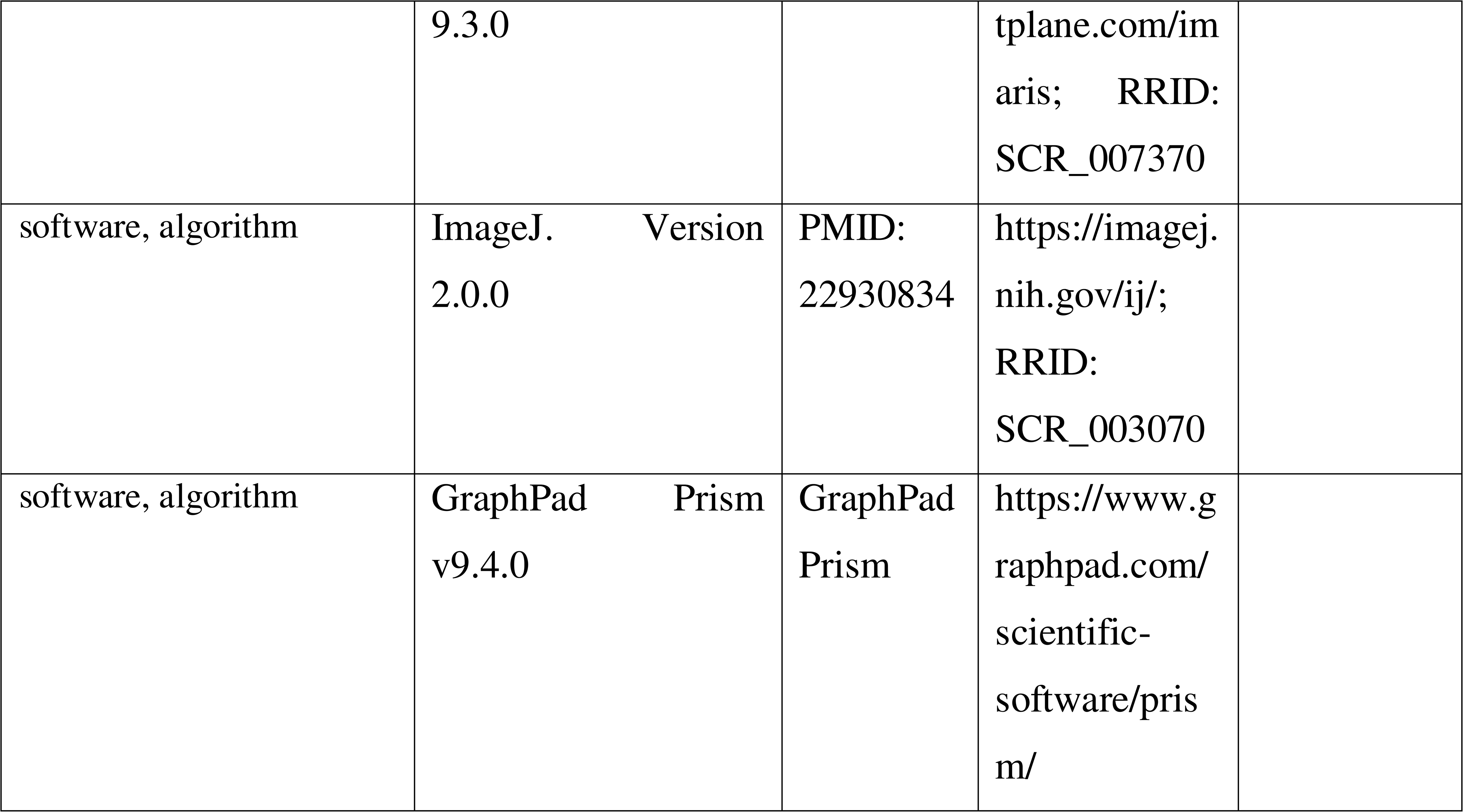

