## Supplementary figures for "Single-cell transcriptome analysis reveals heterogeneity and a dynamic regenerative response of quiescent radial glia in adult zebrafish brain"

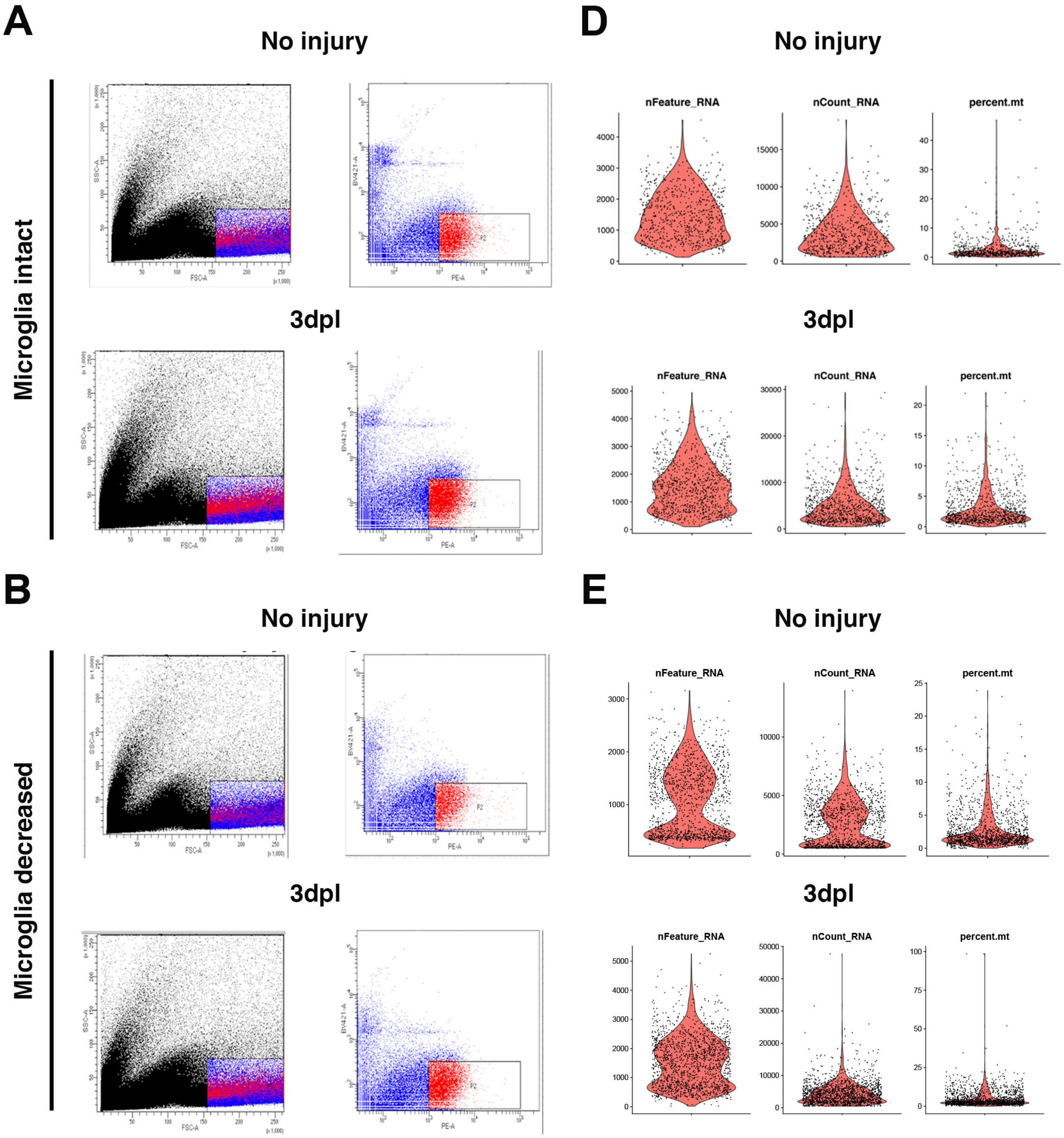


C

| Injury/Microglia Status / Cells Detected per 1000000 Event | No Injury/ Microglia Intact | 3dpl/ Microglia Intact | No Injury/ Microglia Decreased | 3dpl/ Microglia Decreased |
|---|---|---|---|---|
| P2(Target Population) | 7106 | 11322 | 4993 | 8746 |
| P1(Dead Cell Population) | 266081 | 187244 | 298785 | 231610 |
| P2/P1*100 | 2.67% | 6.04% | 1.67% | 3.77% |

Figure2-figure supplement 1

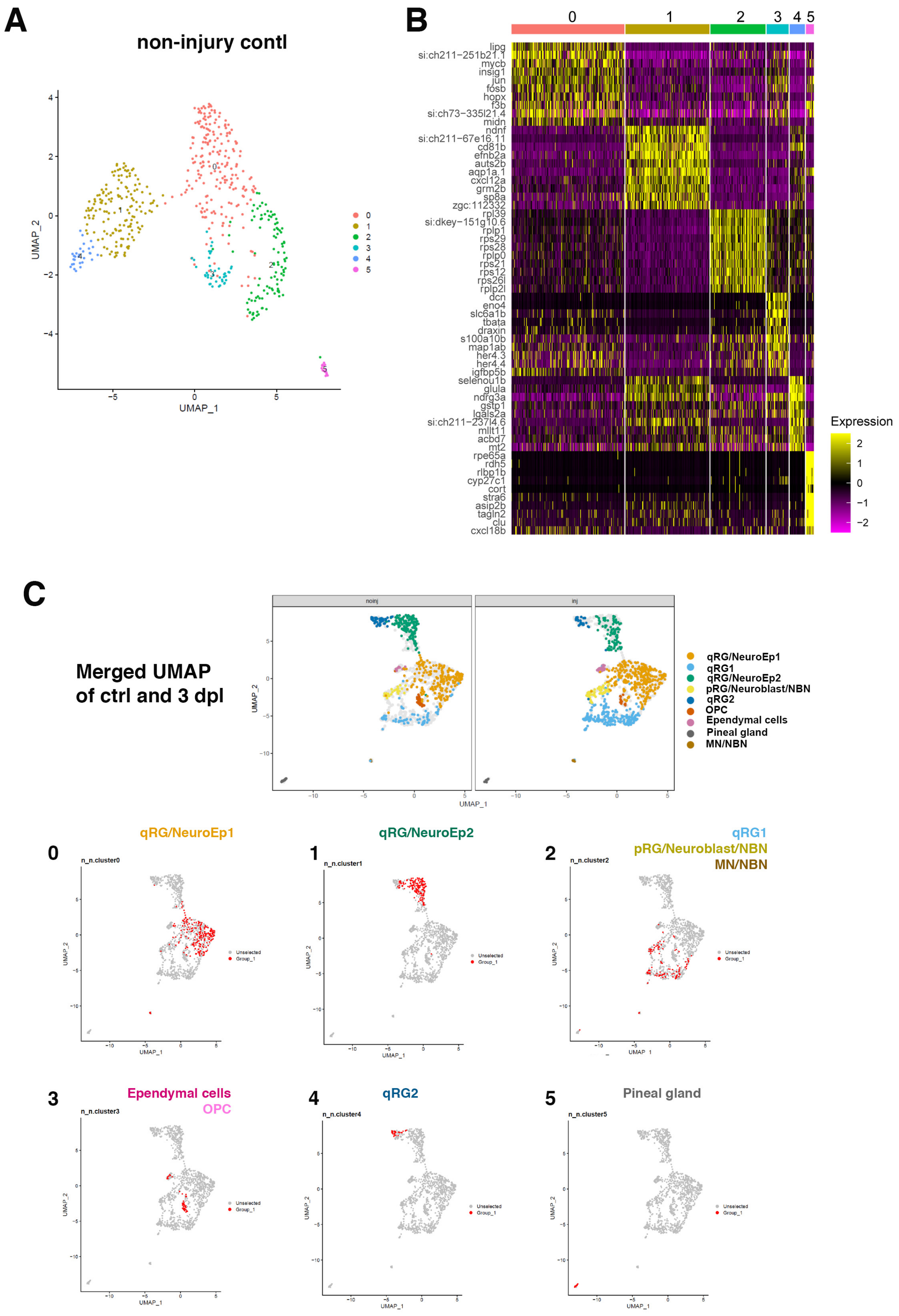


### Figure 2-figure supplement 2

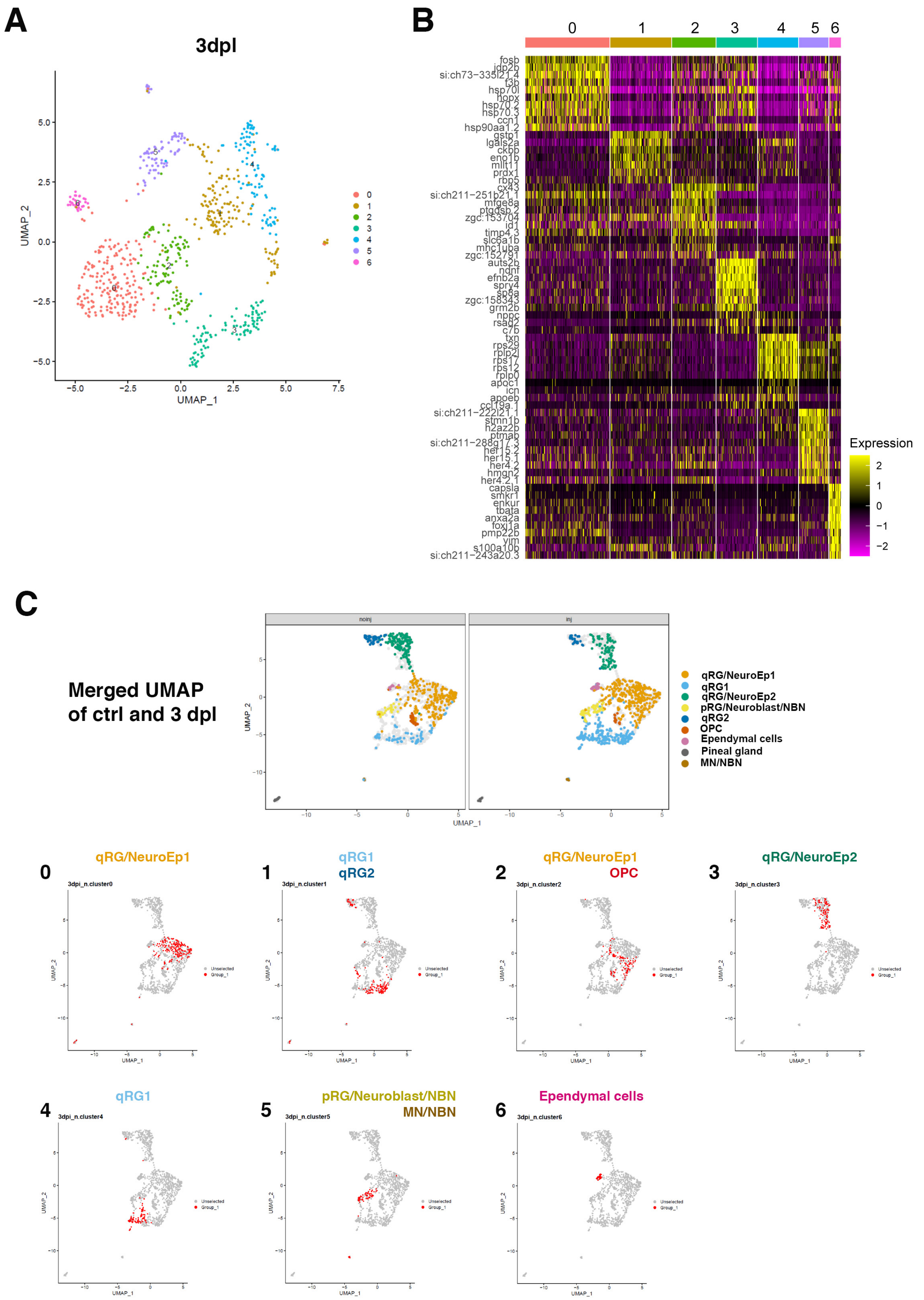


### Figure 2-figure supplement 3

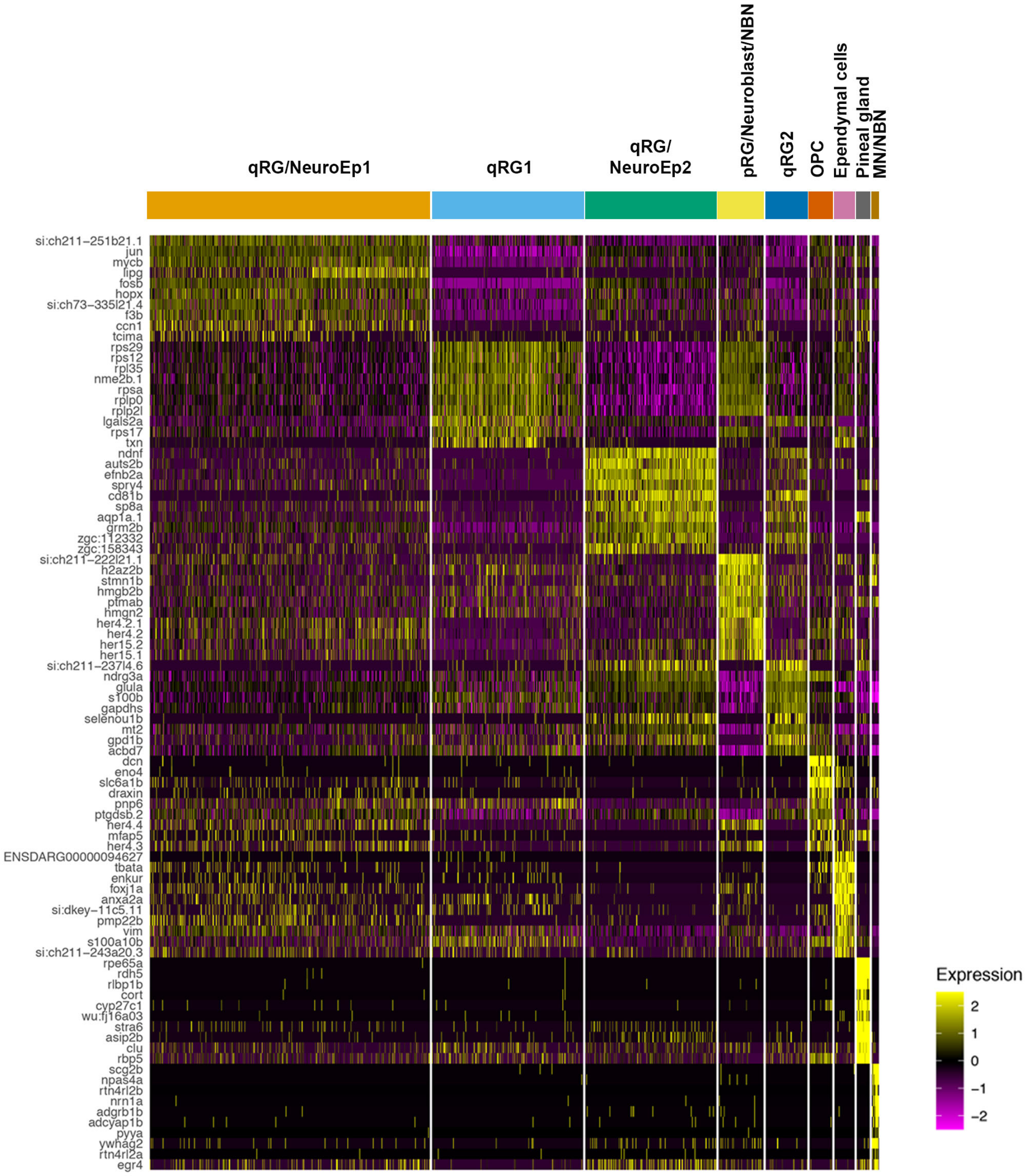


**Figure2-figure supplement 4**

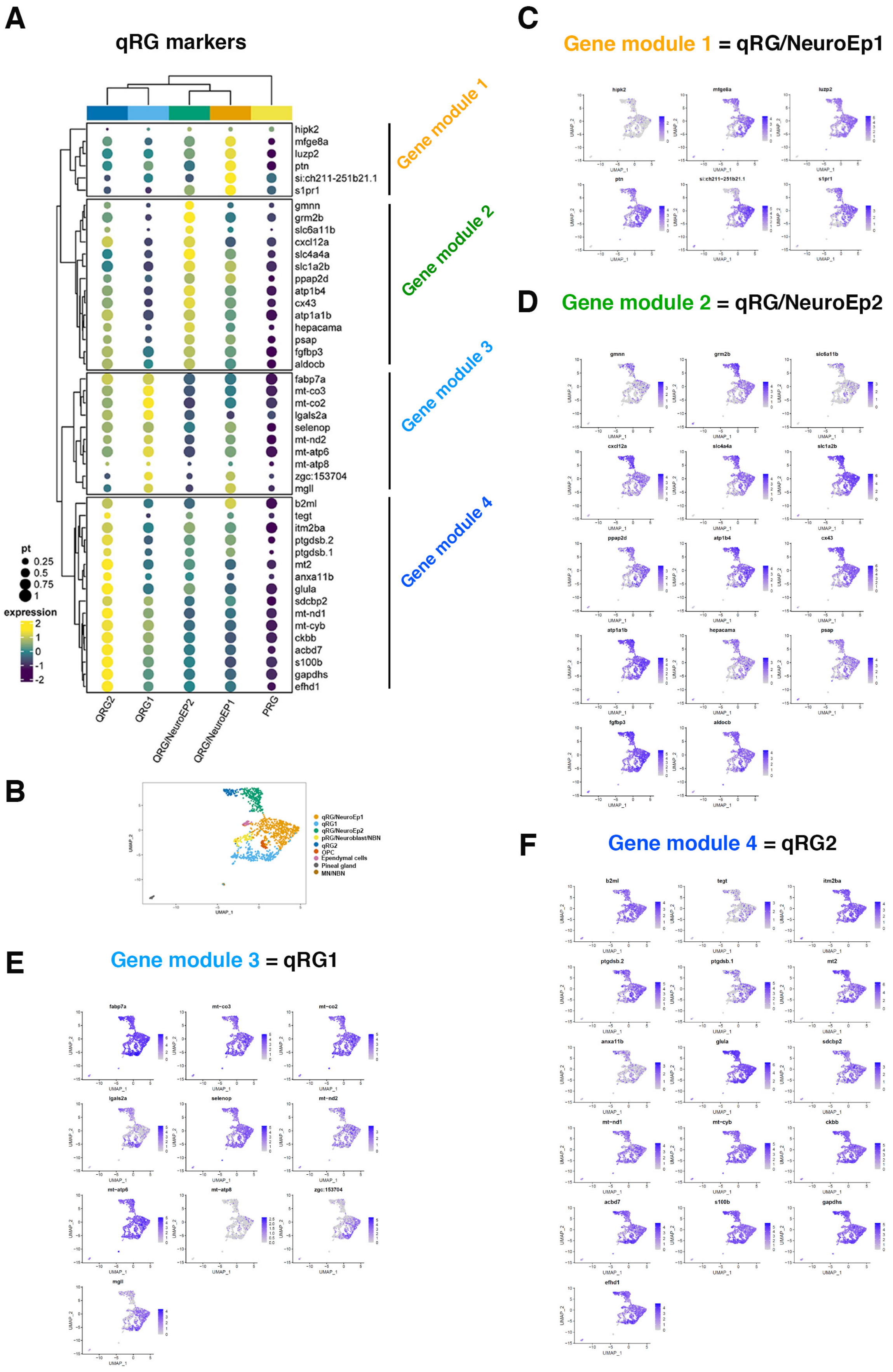


Figure2-figure supplement 5

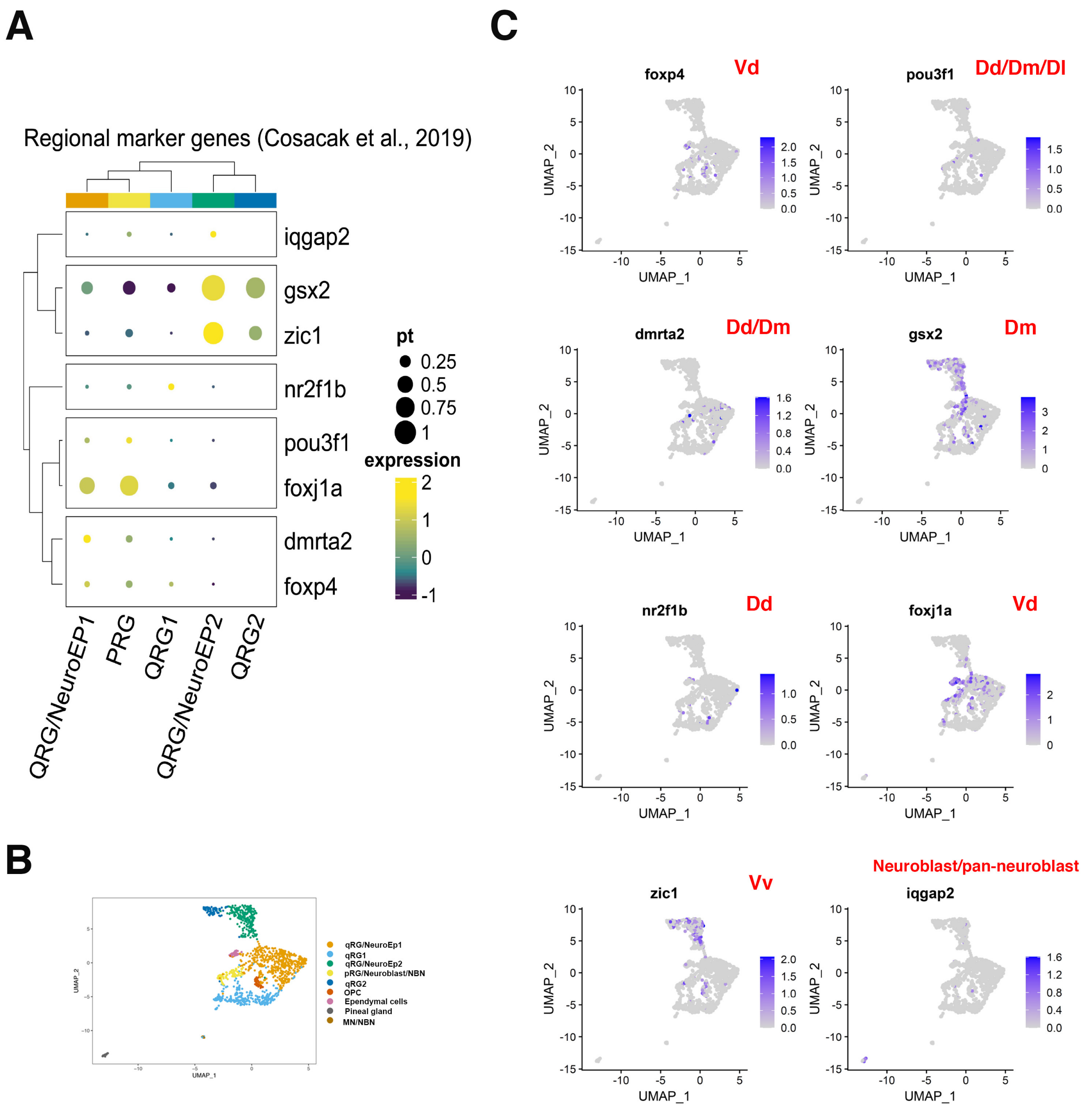


**Fig. 2- figure supplement 6**

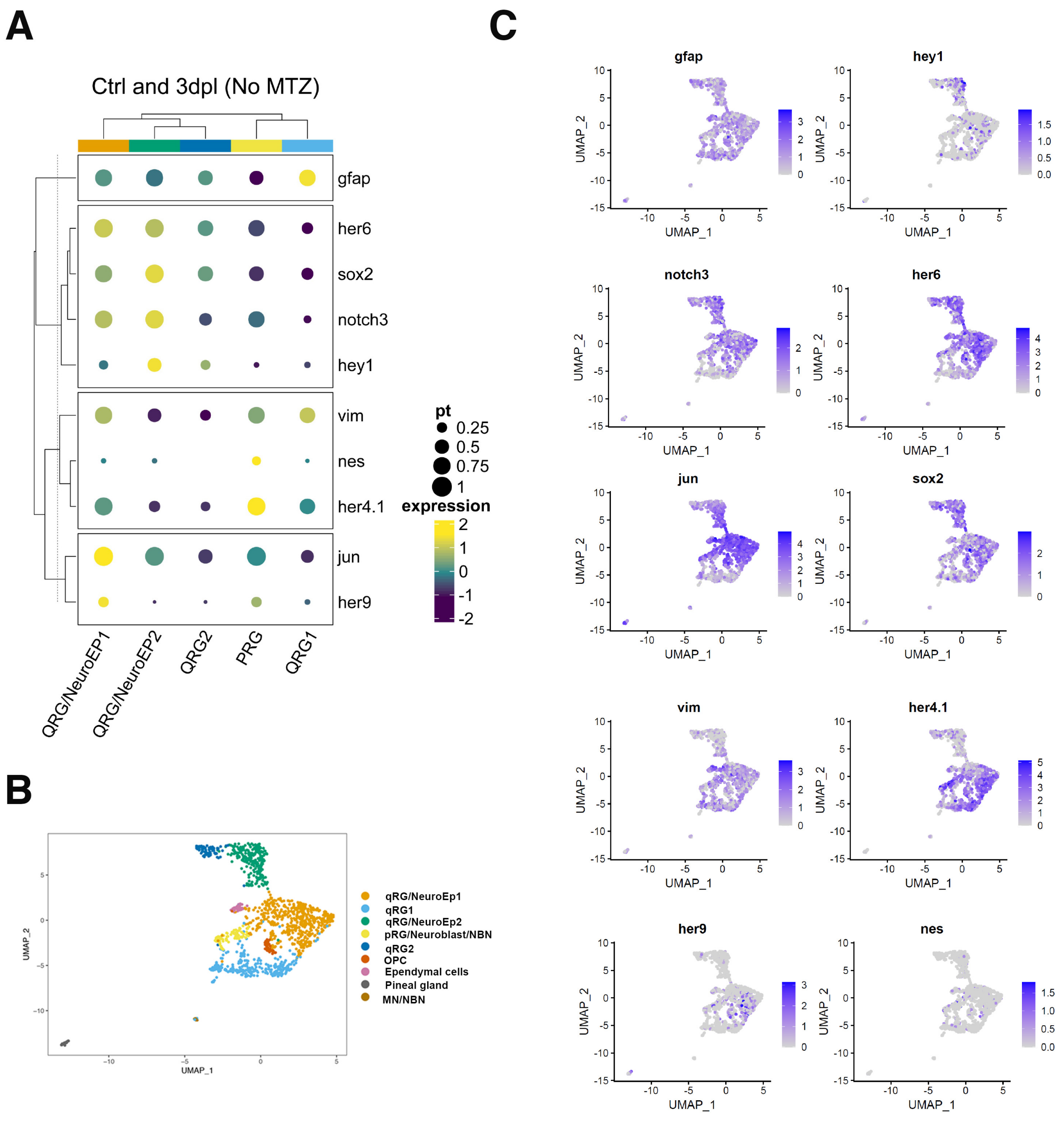


**Fig. 2- figure supplement 7**

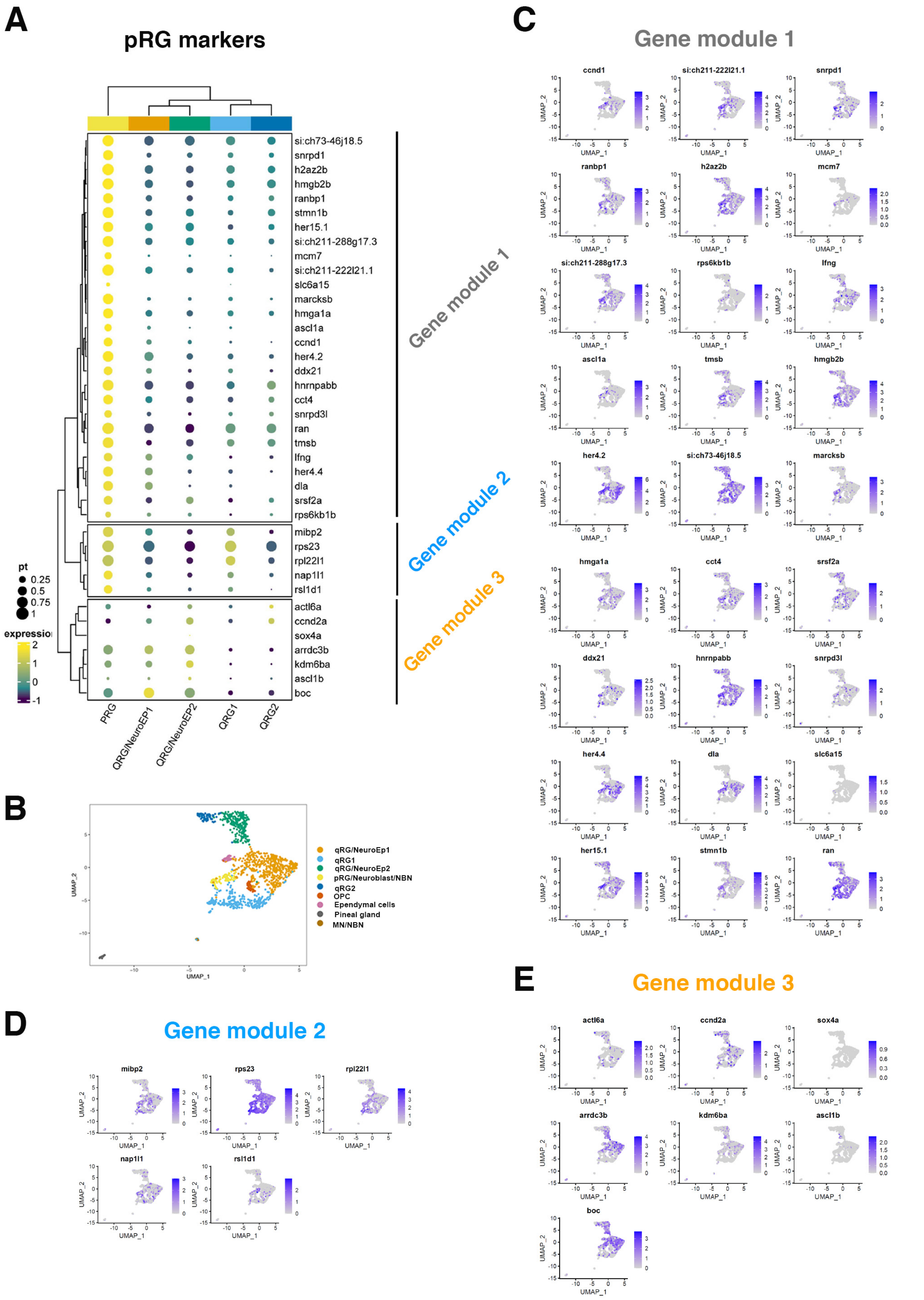


### Figure2-figure supplement 8

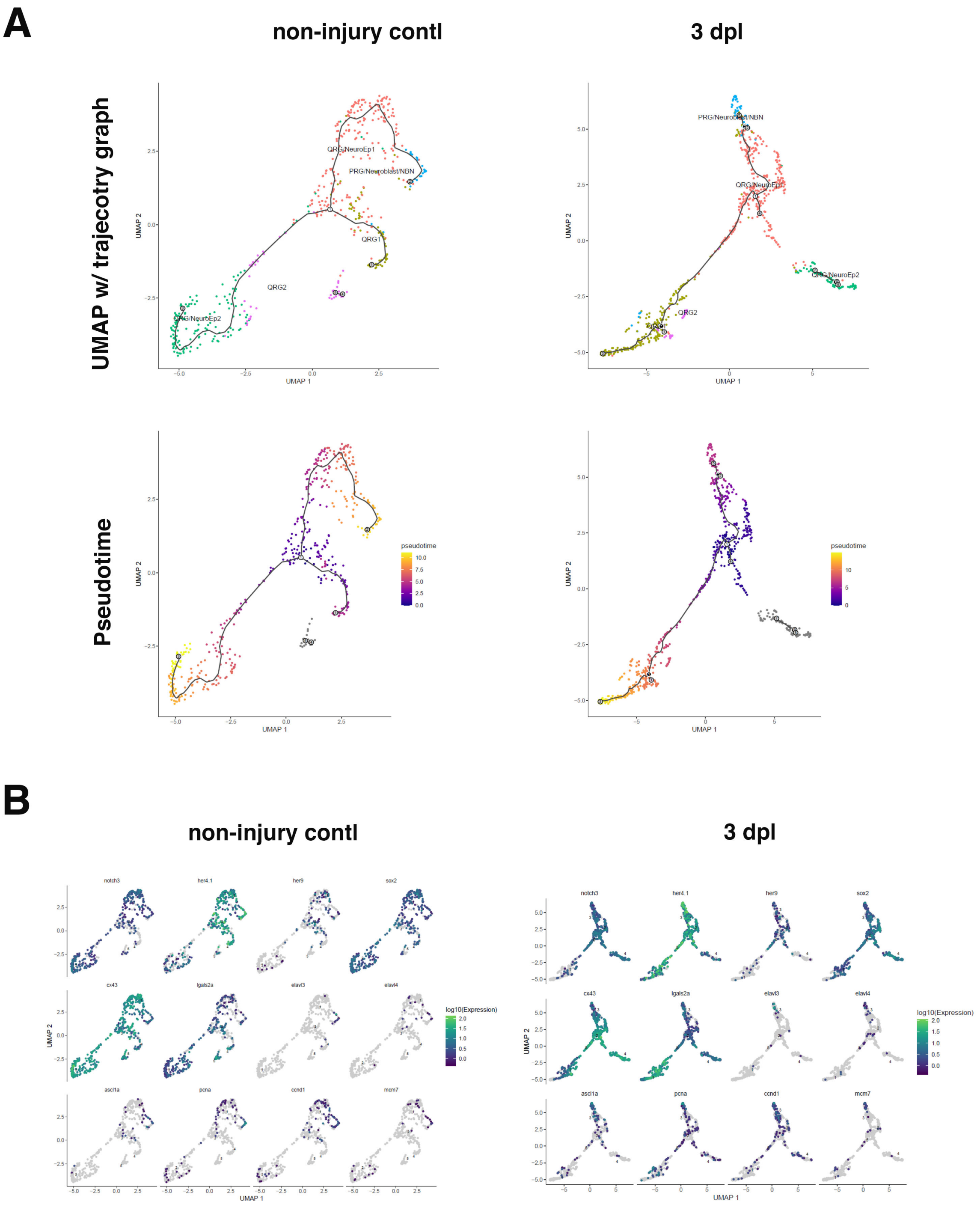


**Fig. 4-figure supplement 1**

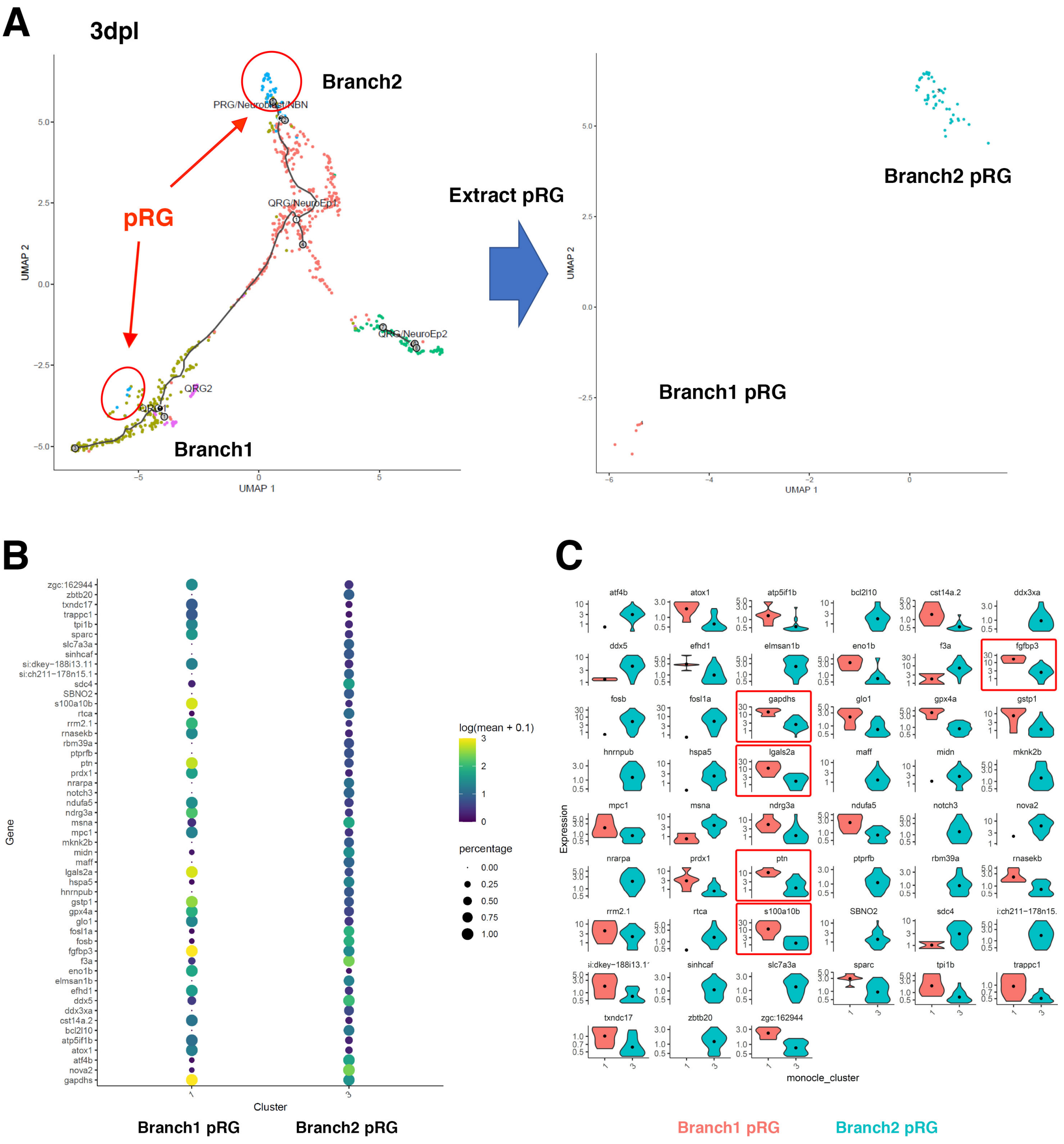


**Fig. 4-figure supplement 2**

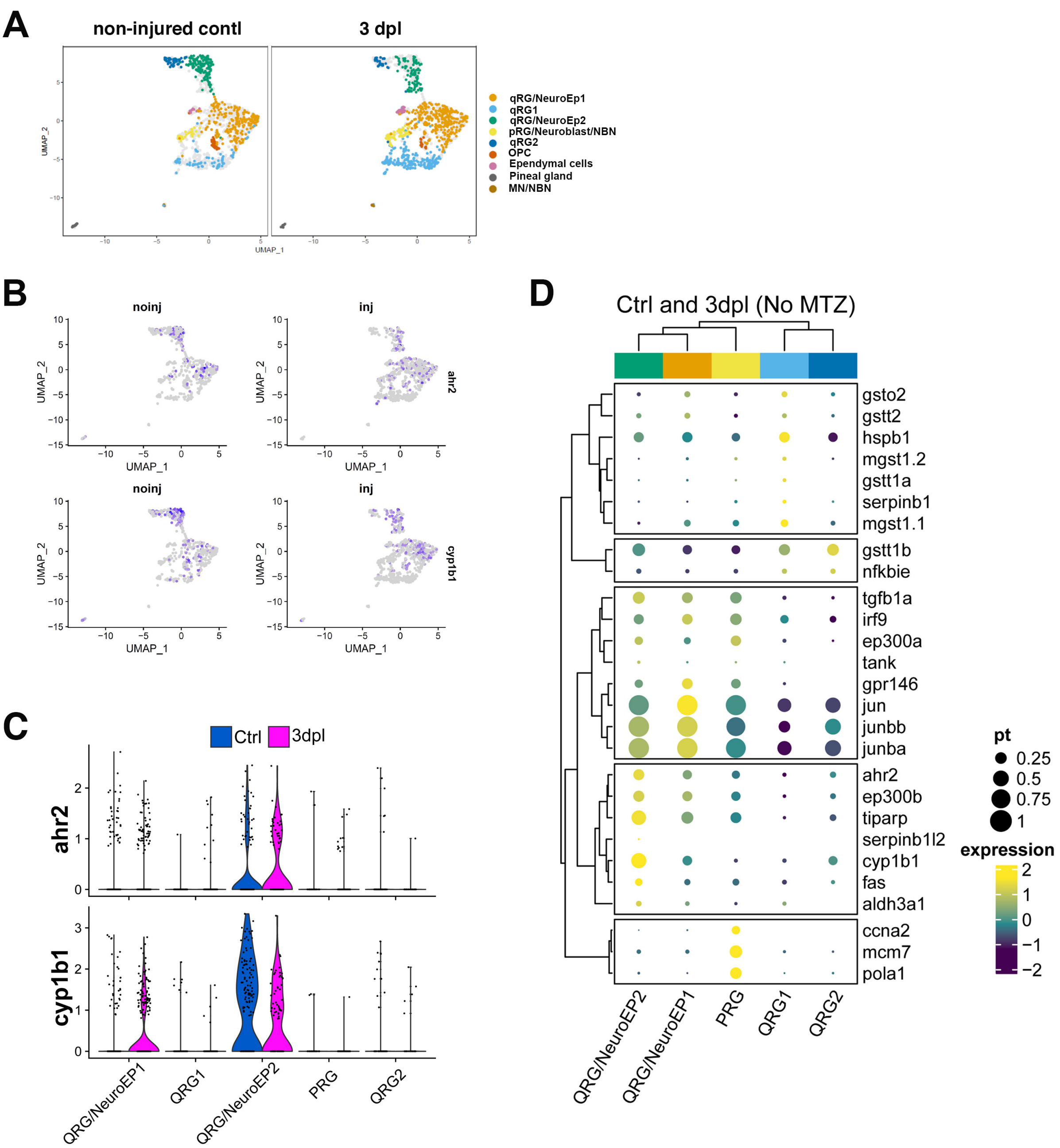


**Figure 4-figure supplement 3**

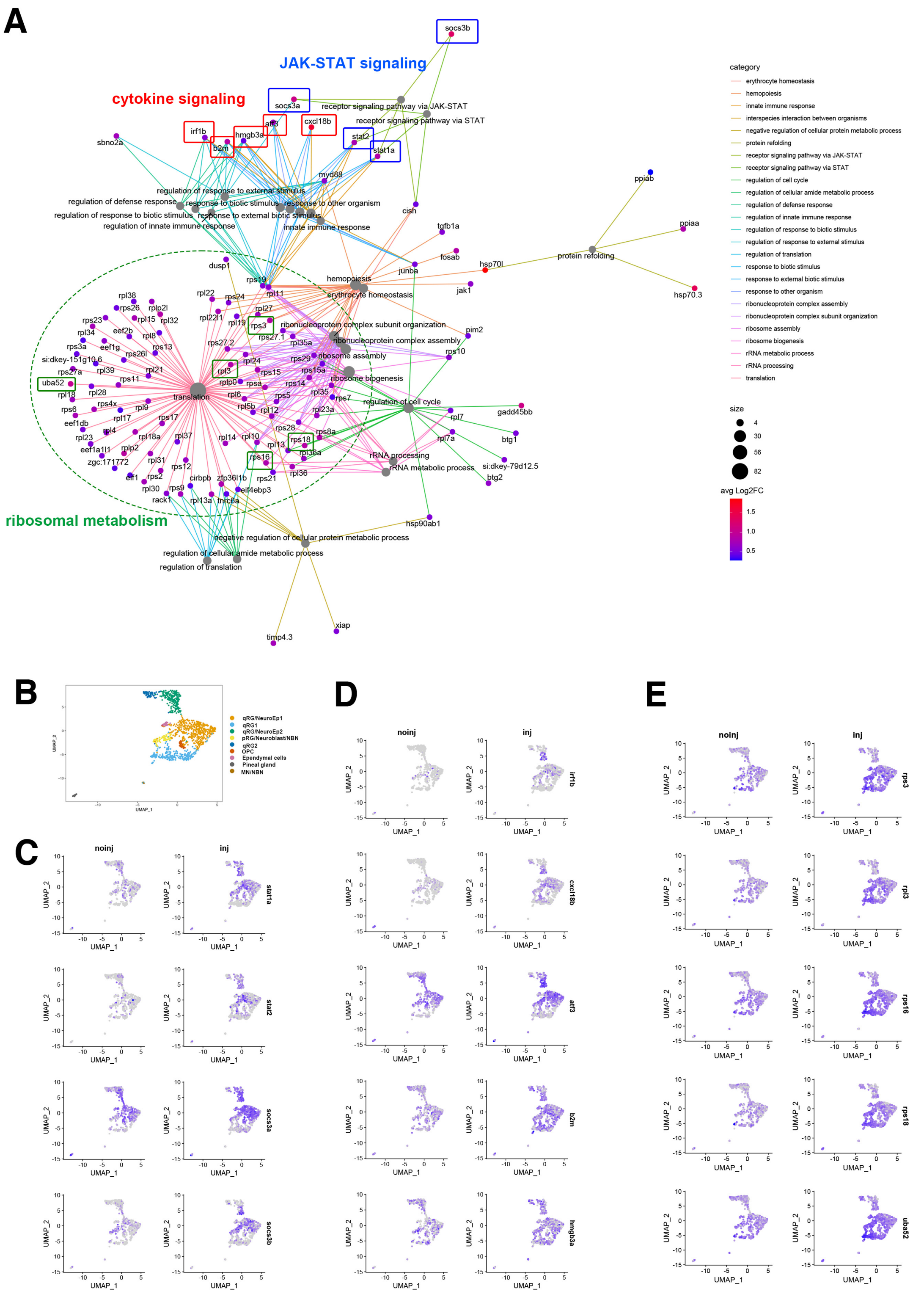


### Fig. 5-figure supplement 1

### Top30 up-regulated genes in pRG/NeuroEp1 (injury condition)

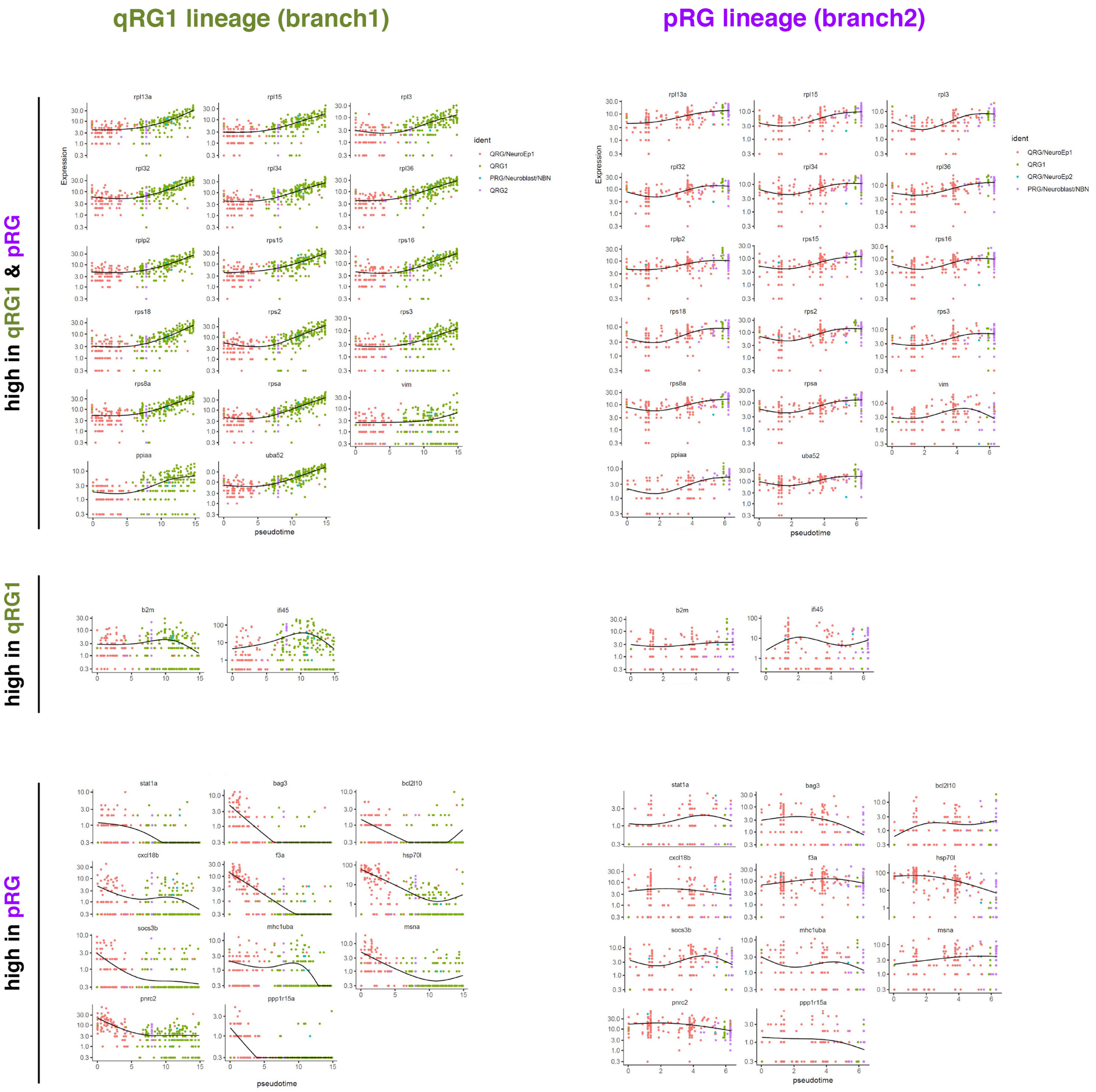


### Figure 5-figure supplement 2

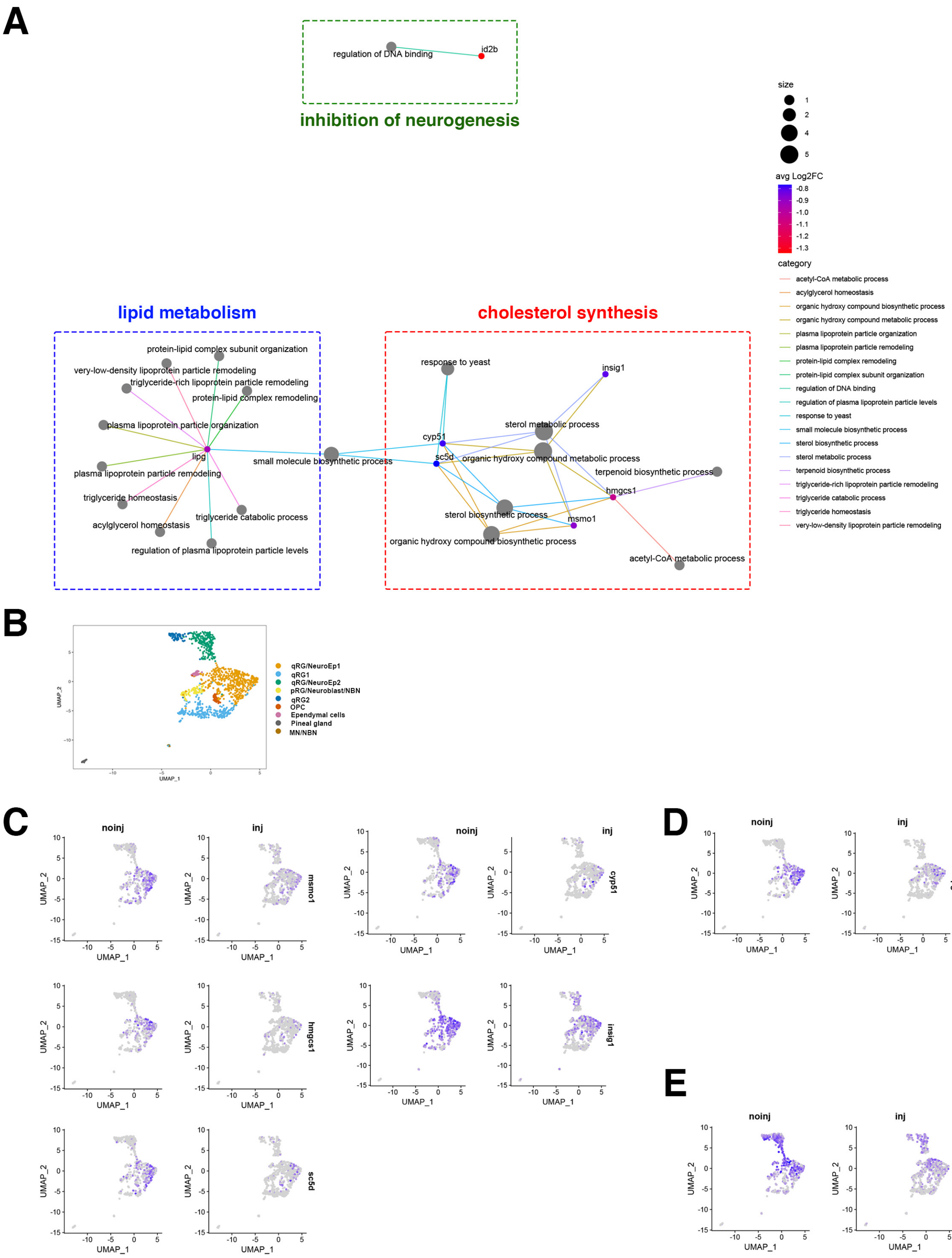


**Fig. 5-figure supplement 3**

**A** **Top12 down-regulated genes in pRG/NeuroEp1 (injury condition)**

**qRG1 lineage (branch1)**

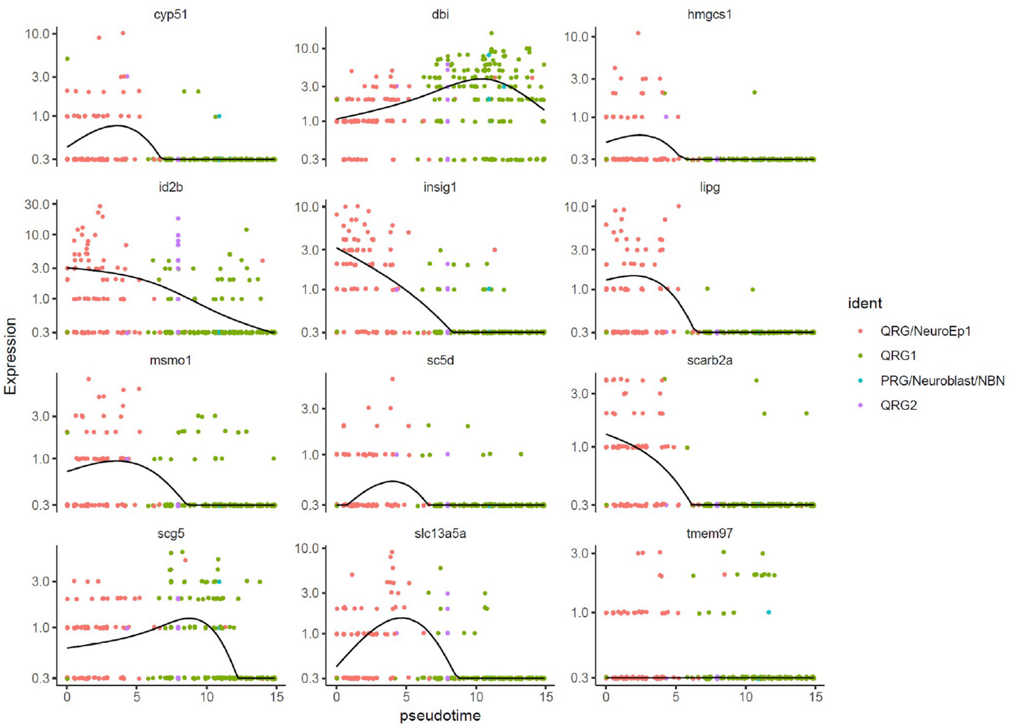


**B**

**pRG lineage (branch2)**

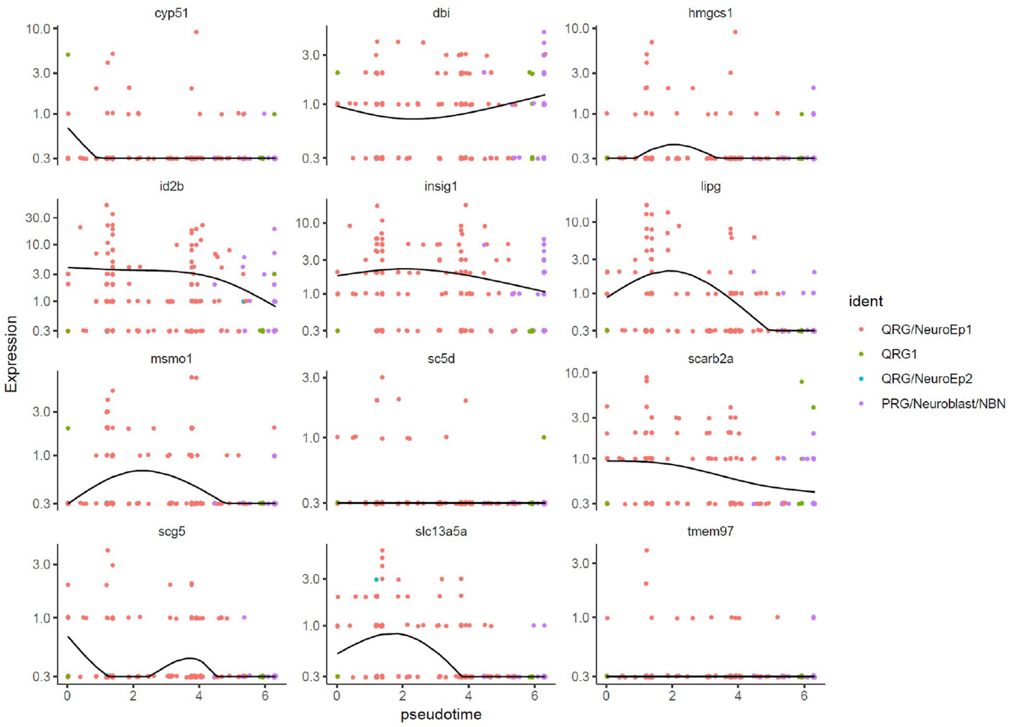


### Figure 5-figure supplement 4

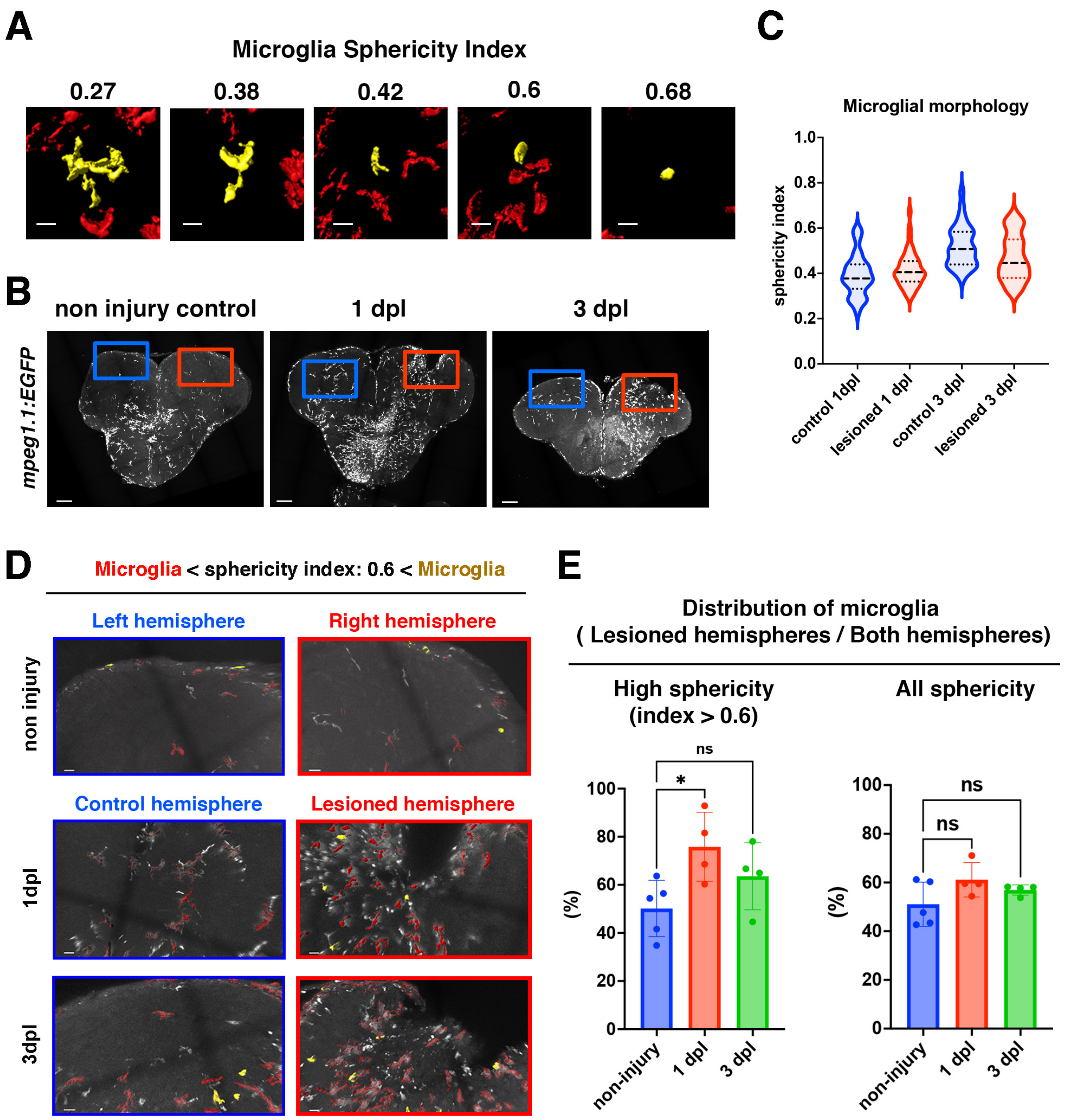


**Fig. 6-figure supplement 1**

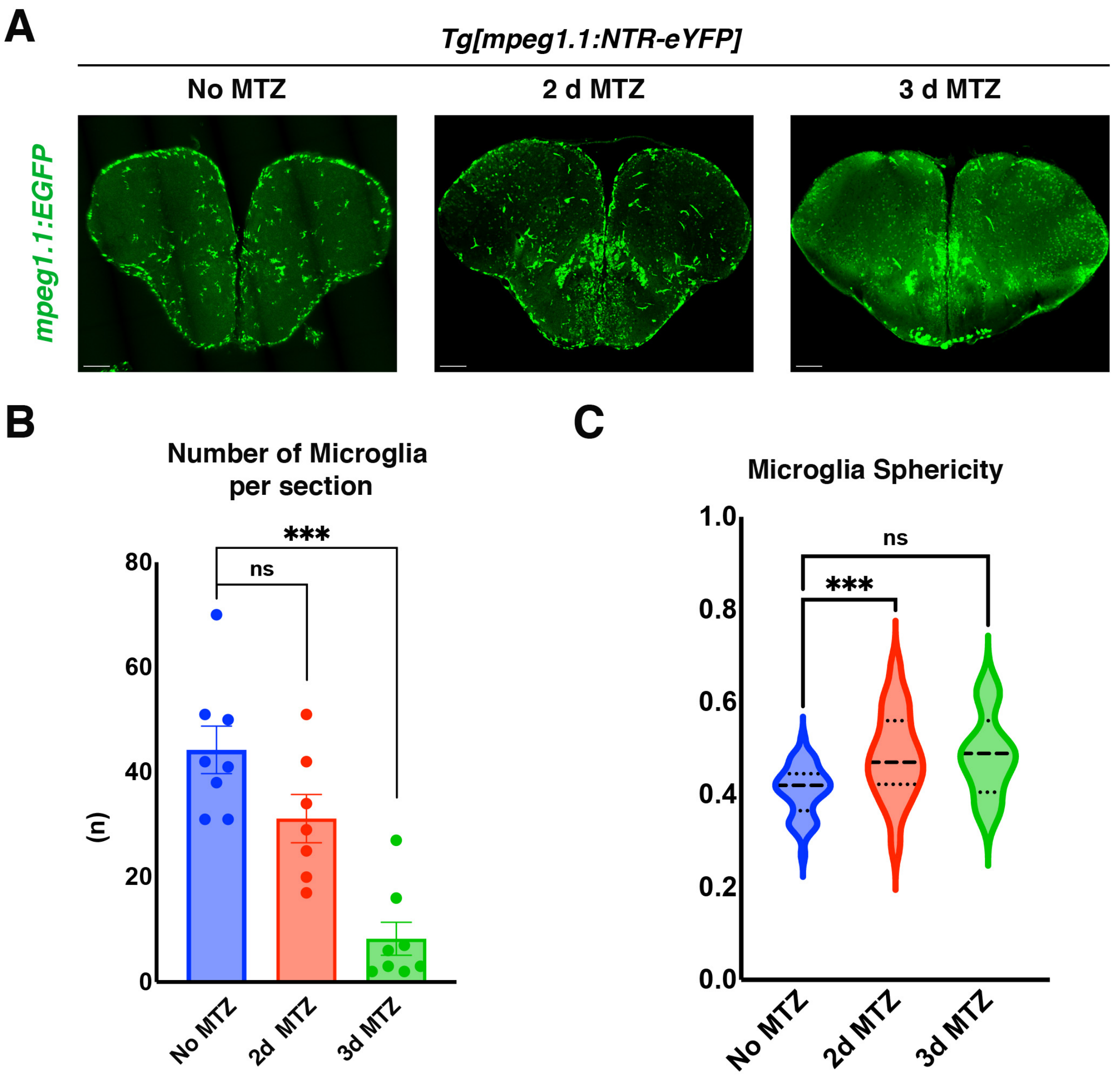


Fig. 6-figure supplement 2

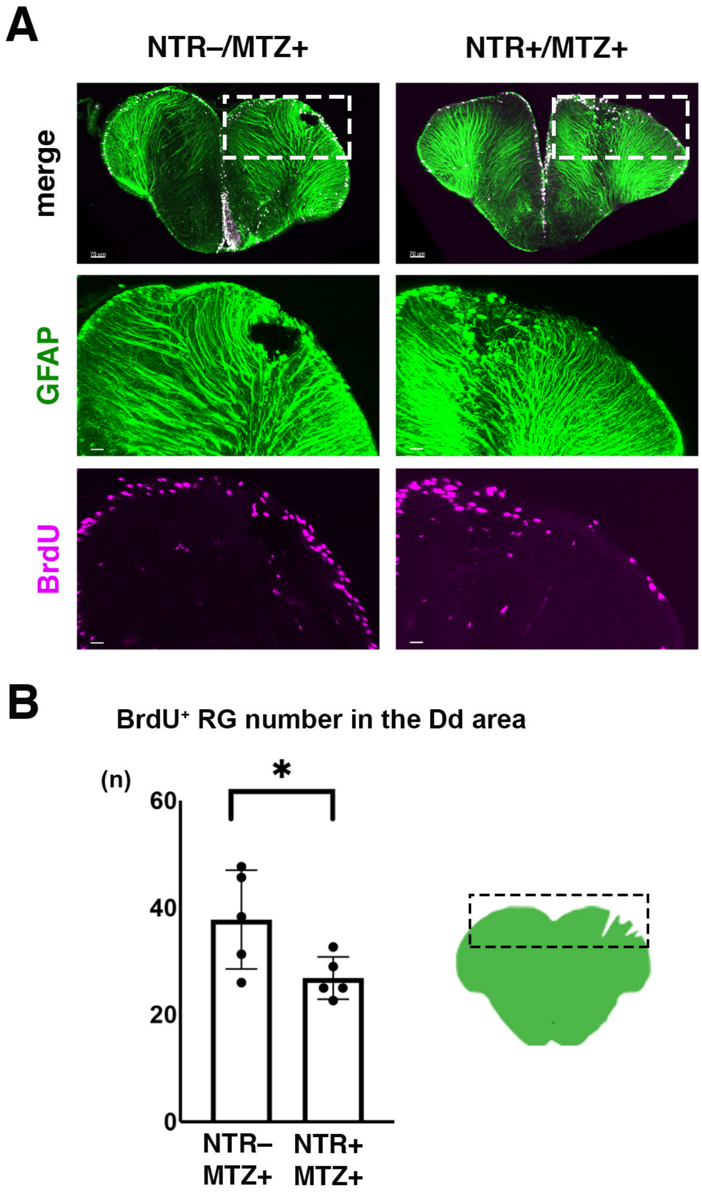


Fig. 6-figure supplement 3

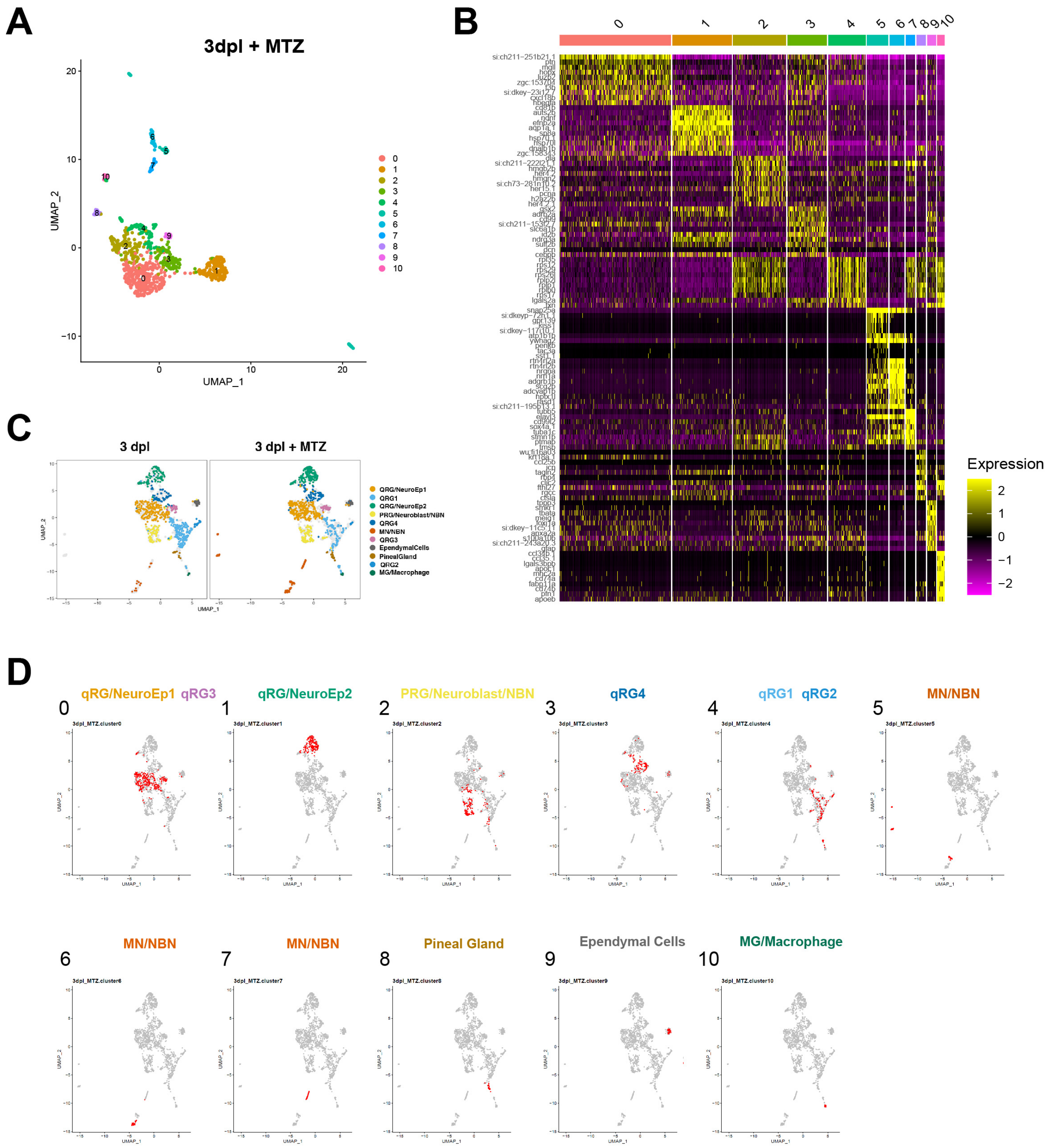


**Fig. 7-Figure supplement 1**

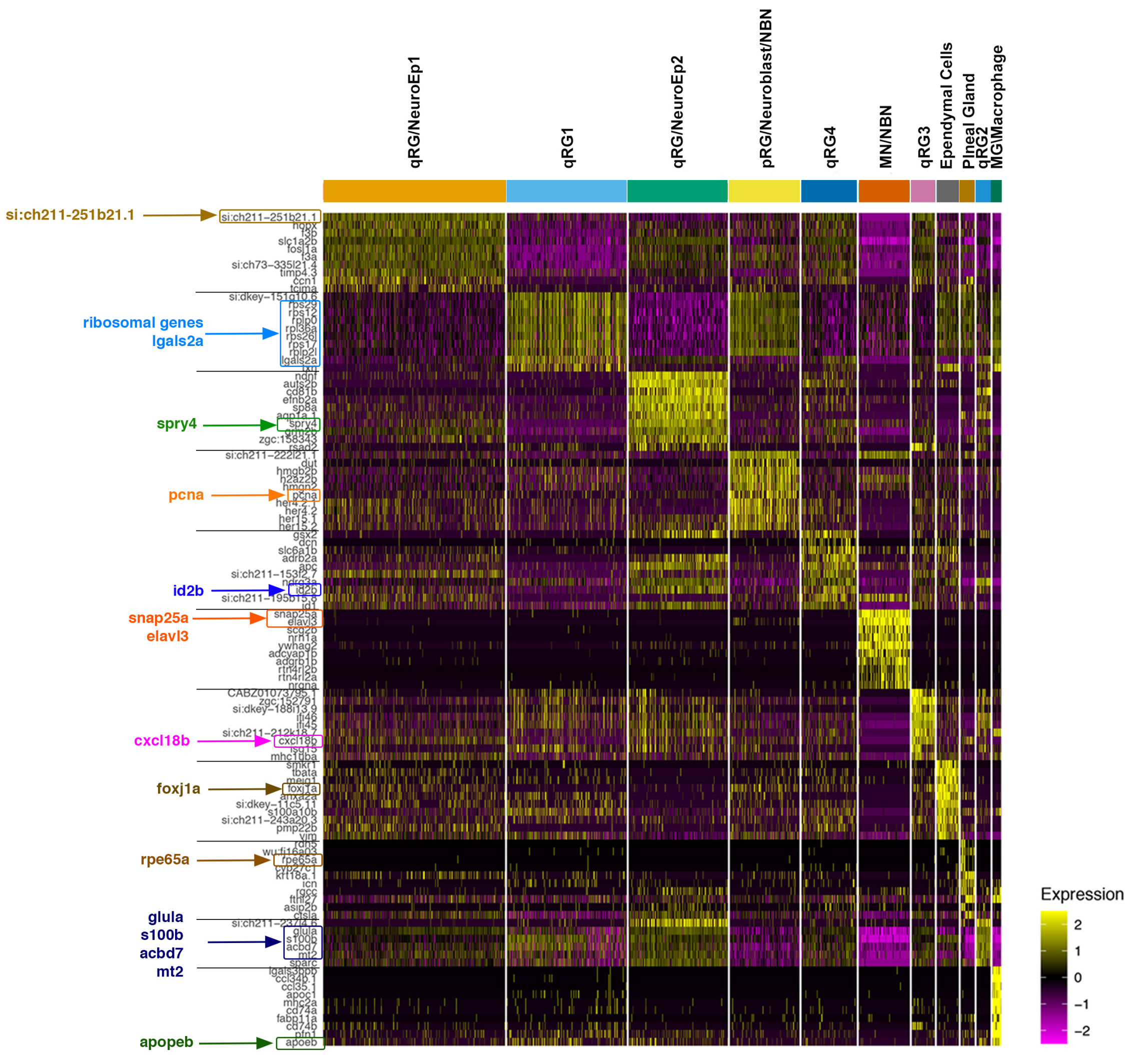


### Fig. 7-figure supplement 2

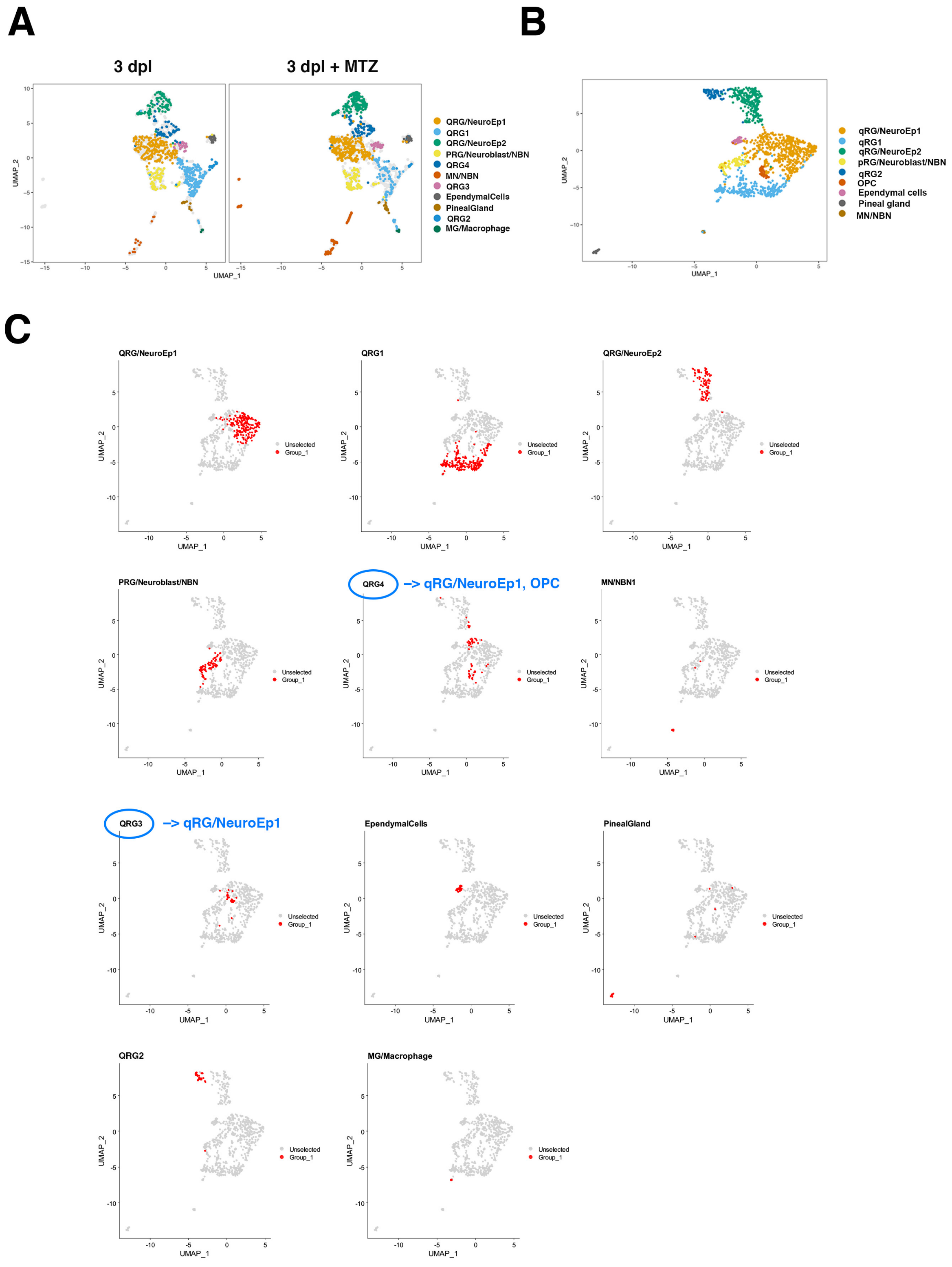


### Fig. 7-figure supplement 3

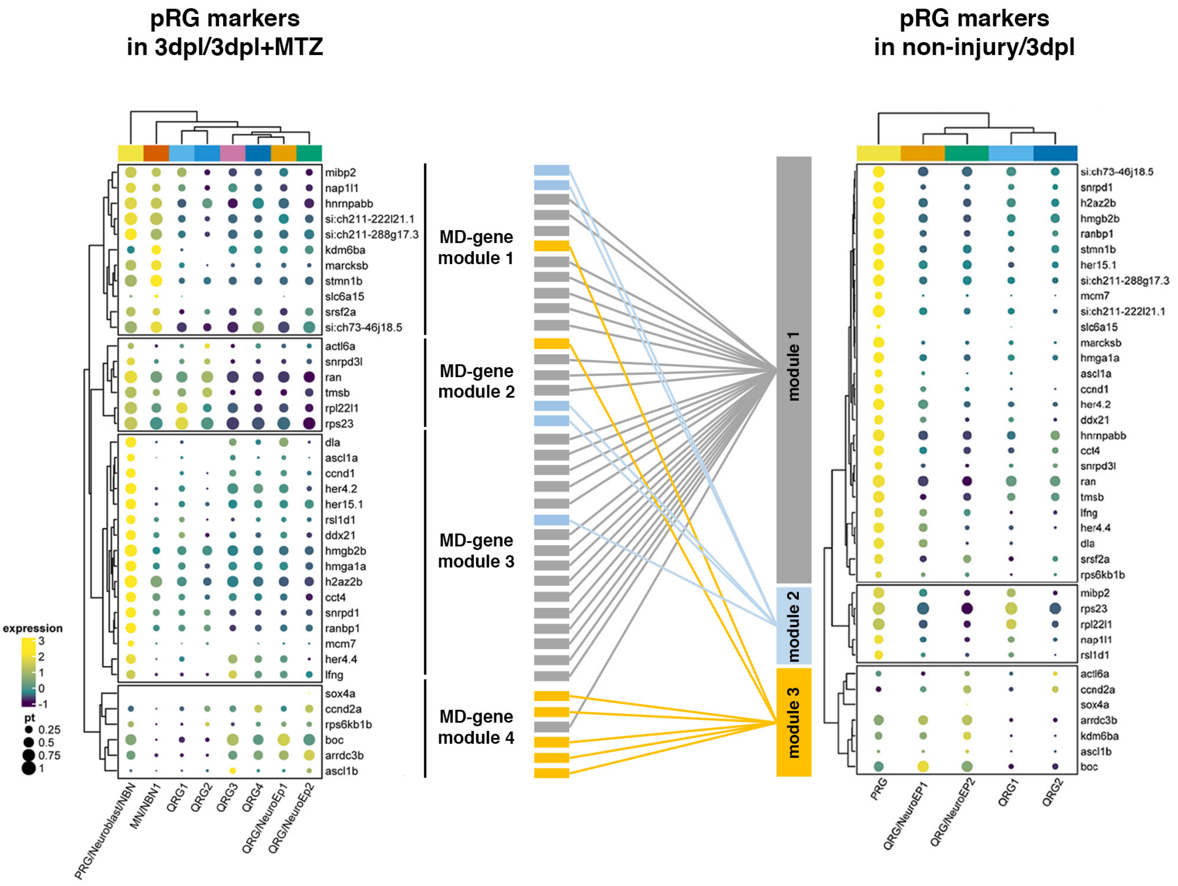


### Figure 7-figure supplement 4

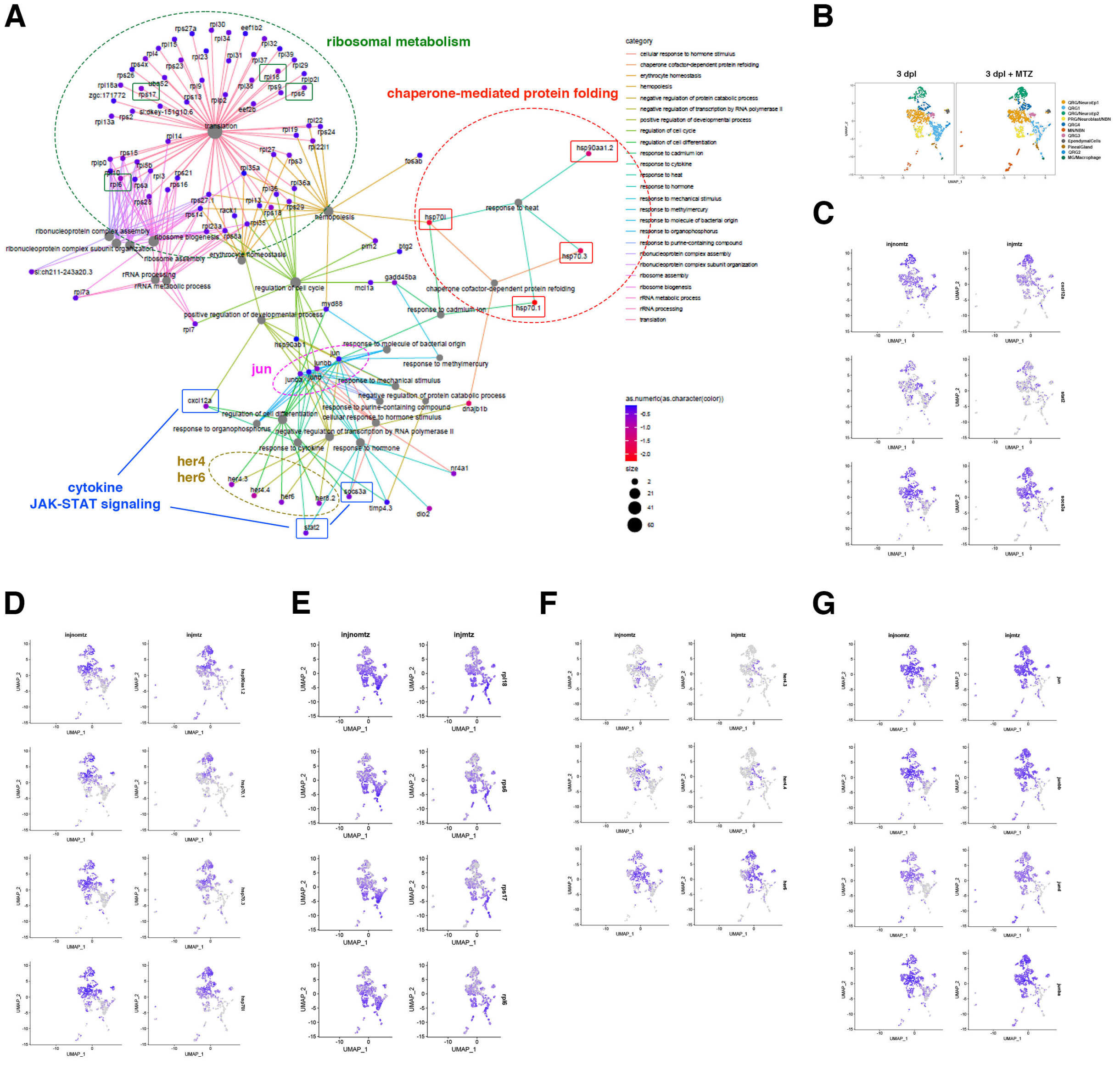


**Fig. 8-figure supplement 1**

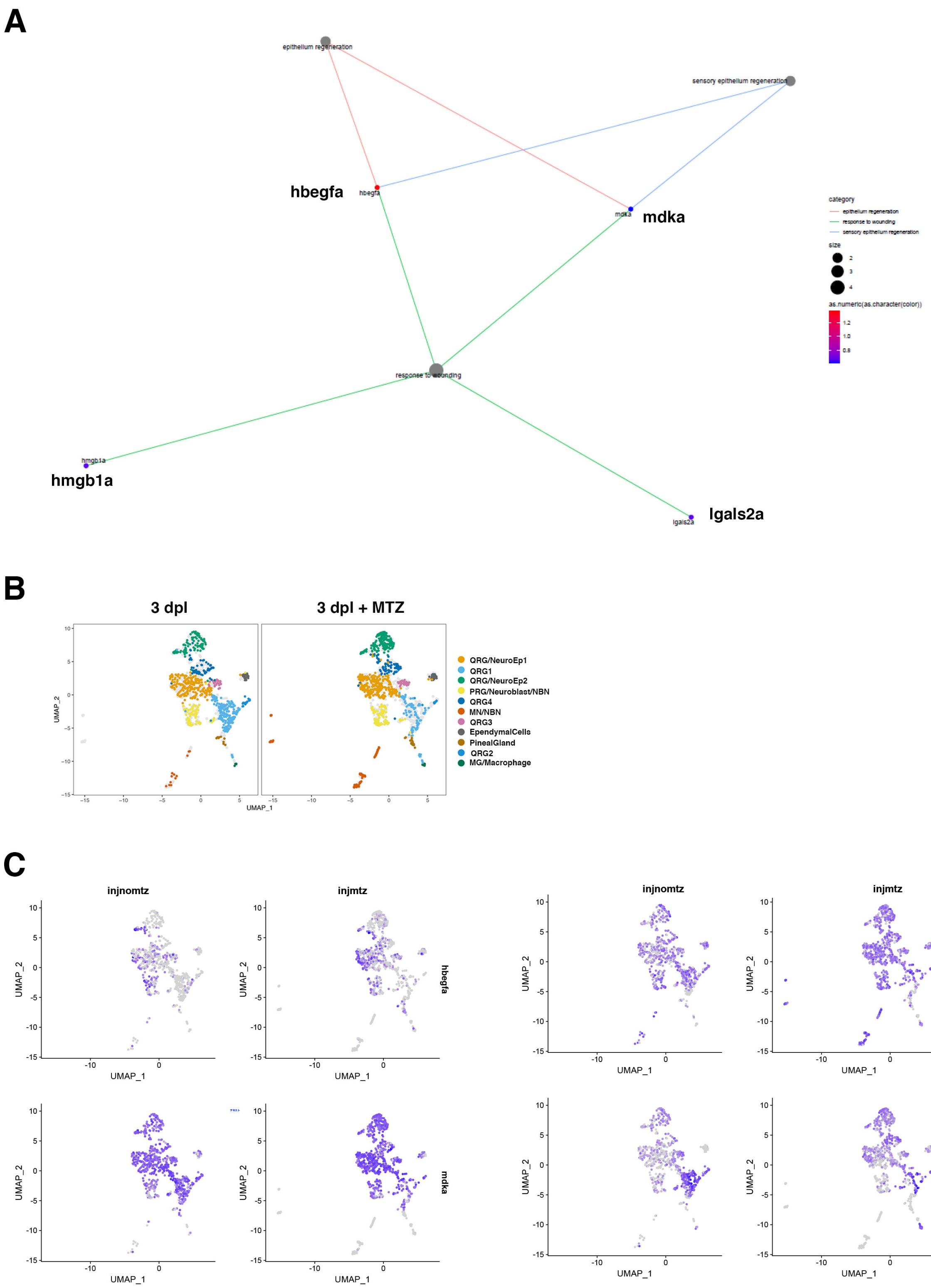


**Fig. 8-figure supplement 2**
